# The structure of the human cell cycle

**DOI:** 10.1101/2021.02.11.430845

**Authors:** Wayne Stallaert, Katarzyna M. Kedziora, Colin D. Taylor, Tarek M. Zikry, Holly K. Sobon, Sovanny R. Taylor, Catherine L. Young, Juanita C. Limas, Jeanette G. Cook, Jeremy E. Purvis

## Abstract

The human cell cycle is conventionally depicted as a five-phase model consisting of four proliferative phases (G1, S, G2, M) and a single state of arrest (G0). However, recent studies show that individual cells can take different paths through the cell cycle and exit into distinct arrest states, thus necessitating an update to the canonical model. We combined time lapse microscopy, highly multiplexed single cell imaging and manifold learning to determine the underlying “structure” of the human cell cycle under multiple growth and arrest conditions. By visualizing the cell cycle as a complete biological process, we identified multiple points of divergence from the proliferative cell cycle into distinct states of arrest, revealing multiple mechanisms of cell cycle exit and re-entry and the molecular routes to senescence, endoreduplication and polyploidy. These findings enable the visualization and comparison of alternative cell cycles in development and disease.

**One-sentence summary:** A systems-level view of single-cell states reveals the underlying architecture of the human cell cycle

## Main text

Our current understanding of the human cell cycle has been assembled piecemeal from decades of individual biochemical and genetic experiments (*1*). The earliest studies focused on the identification of discrete phases of DNA synthesis and cell division (*2*). Advances in molecular biology led to the identification of key cell cycle effectors (e.g. cyclins and cyclin-dependent kinases (CDKs)) and an understanding of the molecular events controlling the DNA replication, mitosis, and cell cycle arrest. This accumulated knowledge has resulted in the adoption of a model that separates the cell cycle into four functionally distinct proliferative phases (G1, S, G2 and M) and cell cycle arrest (G0). This five-phase cell cycle model, which has shaped our thinking for over 70 years (*3*), provides a useful framework for mapping key molecular events that govern the progression of a typical cell through the cell cycle.

However, a wave of recent single-cell studies in cultured cells has revealed that progression through the cell cycle cannot be reduced to a single, fixed sequence of molecular events. Cells can progress at different rates through each of the proliferative phases (*4*–*7*) and vary in the precise molecular paths taken through these phases (*8*–*11*). For example, some cells re-enter the cell cycle immediately following mitosis and begin the molecular preparations for DNA replication in S phase. Other cells divert temporarily to a reversible cell cycle arrest known as quiescence and re-enter the cell cycle after some variable amount of time. Under some conditions, cells can loop back to an earlier point in the cell cycle in response to certain stresses encountered during G1 (*9, 10*) or divert to a different mechanistic route when the canonical path is blocked by pharmacological inhibition (*8*). These studies collectively imply that the cell cycle is a plastic process in which cells may traverse different mechanistic routes *en route* to DNA replication and cell division.

It has been especially difficult to study the behavior of cells once they have exited the proliferative cell cycle and entered a state of cell cycle arrest. It is known, for example, that after entering quiescence individual cells re-enter the cell cycle after variable lengths of time (*11*–*16*), but the mechanisms that regulate this decision remain unclear. It also appears that certain quiescent states can become “deeper” with time, requiring a larger or longer stimulus in order to reverse this arrest and resume the cell cycle (*17*–*19*). In some cases, cells may become irreversibly arrested or “senescent” (*20*–*22*). Precisely how this state varies at the molecular level from reversible arrest has been a longstanding question for the field.

Recent work has begun using single cell measurements to describe the heterogeneity in cell cycle progression. For example, studies using single-cell mRNA sequencing have revealed considerable variability in the transcriptional states of individual cells around a single, cyclical cell cycle trajectory (*23*– *25*). One advantage of these methods is the ability to capture the expression of thousands of different transcripts in the same cell in an unbiased manner. However, models based on transcriptional measurements alone cannot capture many of the core mechanisms known to regulate cell cycle progression. It is widely appreciated that progression through the cell cycle is primarily driven by changes in protein turnover, post-translational modifications or the subcellular localization of key effectors (*1*) — features that cannot be captured by transcriptomics. Other studies using protein-based measurements to study cell cycle progression in individual cells have been focused on mapping the dynamics of cell cycle effectors along a single, one-dimensional trajectory through the proliferative cell cycle (*26, 27*). These approaches are useful for understanding the temporal profile of cell cycle proteins but cannot detect the higher order relationships among effectors in individual cells (e.g., the relationships between cyclins, CDKs, and CDK inhibitors) that are necessary to understand the overall architecture of fate trajectories.

To look for evidence of heterogeneity in cell cycle progression and to better understand the mechanistic basis of this plasticity, we performed time lapse imaging to record the individual cell cycle histories of human epithelial cells followed by iterative immunofluorescence of 48 core cell cycle regulators to obtain high-dimensional, protein-based cell cycle signatures of >30,000 individual cells. We used manifold learning to project these cells onto two- and three-dimensional surfaces to visualize the cell cycle as a sequence of continuous molecular states, ultimately revealing the underlying “structure” of the cell cycle. We also mapped the architecture of cell cycle arrest using various cell cycle stresses, identifying multiple points of divergence from the proliferative cell cycle into distinct states of arrest with unique molecular signatures. In addition, we revealed the fate trajectory through which sustained replication stress can generate polyploidy following mitotic skipping and a general mechanism through which cells can overcome p21 induction and re-enter the cell cycle by increasing the expression of proliferative effectors such as cyclin D.

### Probing the structure of the cell cycle

Starting with an asynchronous population of non-transformed human retinal pigmented epithelial cells (hTERT-RPE-1, abbreviated hereafter as RPE cells, **Fig. 1A**) expressing a single fluorophore cell cycle reporter (PCNA-mTurquoise2 (mTq2)) (*28*), we performed time lapse imaging to record the cell cycle histories of individual cells, including the cell cycle phase (i.e. G0/G1, S, G2 or M) and age (i.e. time elapsed since previous mitosis) of each cell (**Fig. 1B)**. Following time lapse imaging, cells were fixed and then subjected to multiple rounds of immunofluorescence using iterative indirect immunofluorescence imaging (4i) to obtain measurements of 48 core cell cycle effectors (**Table S1**) in a total of 8850 individual cells. From this imaging dataset we extracted 246 single-cell features including the expression and localization of each protein (i.e. in the nucleus, cytosol, perinuclear region and plasma membrane), cell morphological features (e.g. the size and shape of the nucleus and cell), and features of the microenvironment (e.g. local cell density), culminating in a multivariate cell cycle signature for each cell in the entire population.

**Figure 1.**
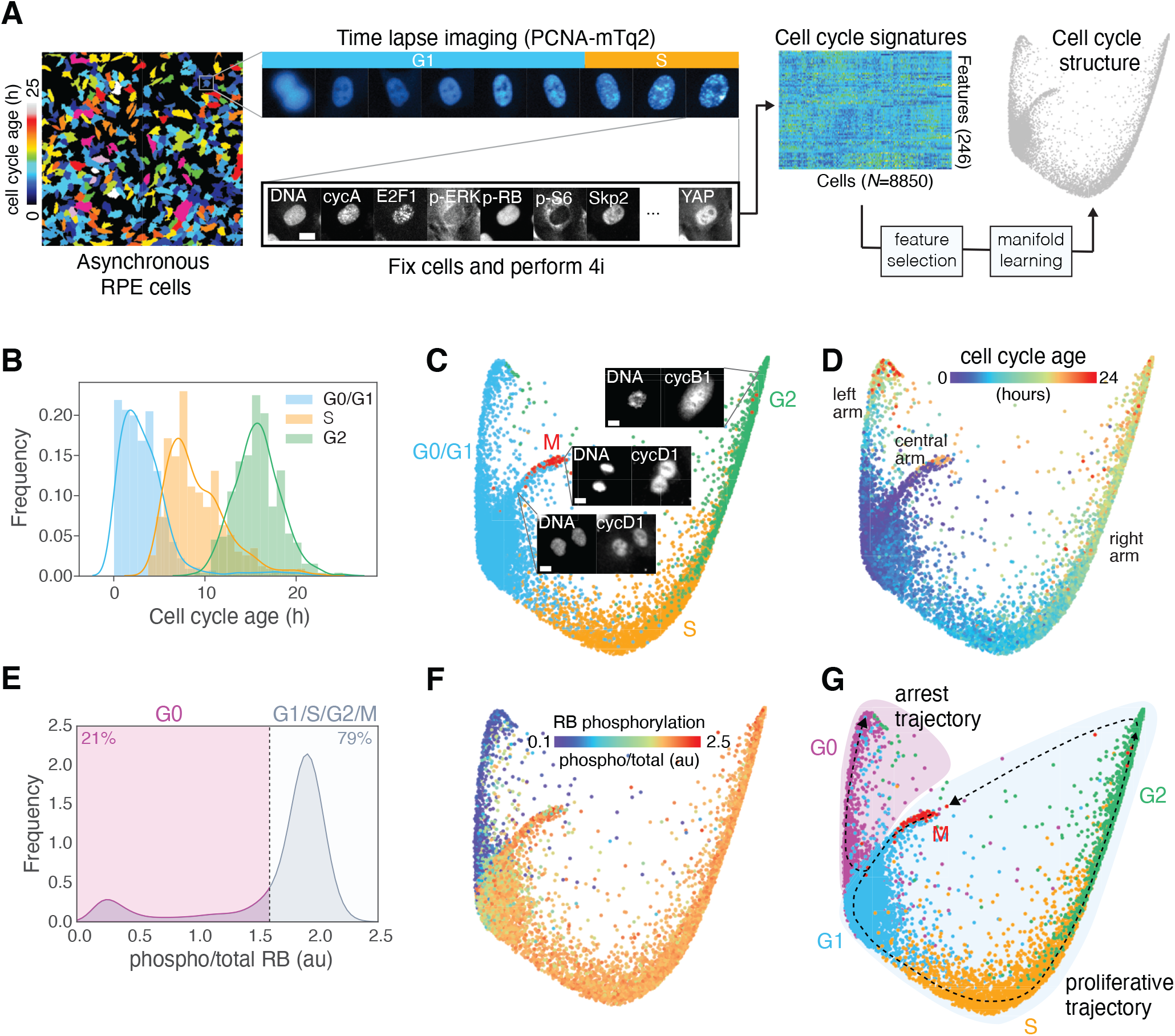
Probing the structure of the cell cycle. (**A**) Schematic of the experimental approach. (**B**) Distributions of cell cycle phase and age (time since mitosis) of all cells annotated by time-lapse imaging. (**C-D**) Cell cycle phase (C) and age (D) annotations/predictions (see Methods) of 8850 individual cells mapped onto the cell cycle structure. Representative images of mitotic cells and their locations on the structure are shown in (C). (**E-F**) Distribution of RB phosphorylation (phospho/total nuclear intensity) in individual cells (E) and mapped on the structure (F). (**G**) Proliferative (G1/S/G2/M) and arrest (G0) trajectories through the five canonical phases. Scale bars = 10 μm.

We then used machine learning to (1) enrich our feature set for variables that best predict the cell cycle state of each cell and (2) eliminate features that vary among cells in a cell cycle-independent manner (see Methods). To do so, we used ground truth annotations of cell cycle phase and age obtained by time lapse imaging to train random forest classifiers to predict these measures of cell cycle ‘state’ solely from the 4i signatures of individual cells (with a 95.5% accuracy for phase and an RMSE=125.8 min and R^2^=0.862 for age predictions). We then averaged the rank order of feature importance from the phase and age models to obtain a combined ranking for each feature. Successive random forests were generated for both phase and age by increasing the number of training features in order of this combined ranking to determine the minimal feature set necessary to accurately predict cell cycle state (i.e. both phase and age) and minimize redundancies (**fig. S1A-C, Table S2**).

We reasoned that cells at similar positions in the cell cycle should possess similar cell cycle signatures (as defined by 4i), and thus be located close to one another on a lower dimensional manifold within the high-dimensional space of this minimal feature set. To identify and visualize this manifold, we used Potential of Heat-diffusion for Affinity-based Transition Embedding (PHATE), a nonlinear manifold learning approach that performs well for continuous and branched trajectories in high-dimensional space (*29*).

This data-driven approach produced a continuous, three-armed structure representing the collective cell cycle states of every cell in the population (2D structure in **Fig. 1**; 3D structure in **Movies 1-4**; virtual reality-compatible dataset available (see Acknowledgments)). We found that this three-armed structure was reproducible across PHATE parameter space (**fig. S2A**), individual replicates (**fig. S2B**) and in another human epithelial cell line (human pancreatic epithelial cells, HPNE, **fig. S2C**).

Cell cycle annotations were obtained by time lapse imaging for only a subset of cells (33%) in order to preserve a temporal resolution that can accurately delineate cell cycle phase transitions (1 frame/16 min) (**fig. S1D-E**, see Methods). We used the random forest models that were trained to predict cell cycle phase and age based on the 4i signatures of cells (described above) to infer the cell cycle phase and age of cells that were not captured by time lapse imaging (**Fig. 1C-D, Movies S2** and **S3**). Similar inferences were obtained using a convolutional neural network trained on the same data (Phase: 95.7% accuracy, concordance with RF model = 94.8%; Age: RMSE = 123.7 min; **fig. S1F-K**). Mapping the cell cycle phase of each cell onto the structure, we observed that G0/G1 cells encompass most of the left and central arms, while S and G2 cells progressively reside along the rightmost arm, respectively (**Fig. 1C**). While cell cycle progression towards mitosis is clearly represented along the right arm of the structure, M phase cells were located at the tip of the central arm. Visual inspection of the immunofluorescence images of cells along the central arm revealed a progression through nuclear envelope reformation and cytokinesis, events consistent with the late stages of mitosis. Similarly, cells at the tip of the right arm possessed features of late G2 and early M, including DNA condensation and nuclear envelope breakdown. Mapping cell cycle age onto the structure indicated that the youngest cells lie near the tip of the central arm where newborn cells emerge out of mitosis and into G1, and that age increases as cells progress up both the left and right arms (**Fig. 1D**). The close correspondence between cell cycle age (determined by time lapse imaging) and the continuum of molecular states in the structure (determined by the similarity of 4i cell cycle signatures) provided confirmation that this structure represents an accurate depiction of the temporal trajectories of the cell cycle.

While the PCNA-mTq2 reporter allows delineation of the proliferative phases of the cell cycle (G1/S/G2/M) (*28*), it cannot distinguish G0 from G1 cells. The phosphorylation and inactivation of the retinoblastoma protein (RB), on the other hand, represents a major checkpoint controlling cell cycle re-entry and is often used to define the boundary between G0 (i.e. cell cycle arrest) and G1 (i.e. cell cycle commitment) (*30*– *32*). In our 4i data, we observed bimodality in RB phosphorylation (**Fig. 1E**), and a clear delineation between these two cellular states (high vs. low RB phosphorylation) along the central arm, thus distinguishing G0 from G1 cells (**Fig. 1F, Movie 4**). We use this RB phosphorylation status to identify actively proliferating versus arrested cells throughout the manuscript.

We therefore infer two principal trajectories within this cell cycle structure: (1) a cyclical, *proliferative trajectory* starting at the base of the central arm and progressing along the right arm through G1, S and G2, before looping back to the central arm during mitosis, and (2) an *arrest trajectory* along the left arm (**Fig. 1G**).

### Mapping the mechanisms of the proliferative cell cycle (G1/S/G2/M)

To validate our approach, we used cell cycle age annotations to order cells temporally along the proliferative trajectory and examined key mechanisms previously shown to drive cell cycle progression (**Auxiliary fig. S1**). This approach successfully ordered key molecular events regulating the G1/S transition (**Fig. 2A**) (*33*). The core molecular unit regulating this decision is the RB-mediated inhibition of E2F transcription factors, which control the expression of various S phase genes (*34*). Commitment to DNA replication is triggered by the phosphorylation and inhibition of RB by cyclin:CDK complexes. In early G1, RB is primarily phosphorylated by cyclin D:CDK4/6 (*16*), and we observed that cells begin their cell cycle with high cyclin D1 expression (**Fig. 2B**). This cyclin D:CDK4/6-driven phosphorylation of RB in early G1 de-represses E2F-regulated genes important for DNA replication including expression of E2F1 itself (**Fig 2C**) (*35, 36*). E2F activity also stimulates the production of cyclin E (**Fig. 2D**), which activates CDK2 to maintain RB phosphorylation as cyclin D levels begin to decrease (**Fig. 2B**) (*16, 27*). Another important event in the G1/S transition is the inactivation of APC/C complexes, which degrade and prevent the accumulation of S phase proteins during G1. The increase in cyclin E:CDK2 activity in late G1 stimulates the destruction of the Cdh1 subunit (**Fig. 2E**), switching off APC/C and permitting S phase initiation (*9*). The inactivation of APC/C also allows cyclin A to accumulate as S phase begins (**Fig. 2F**) to replace cyclin E and maintain CDK2-dependent RB phosphorylation through to mitosis (**Fig. 2D**).

**Figure 2.**
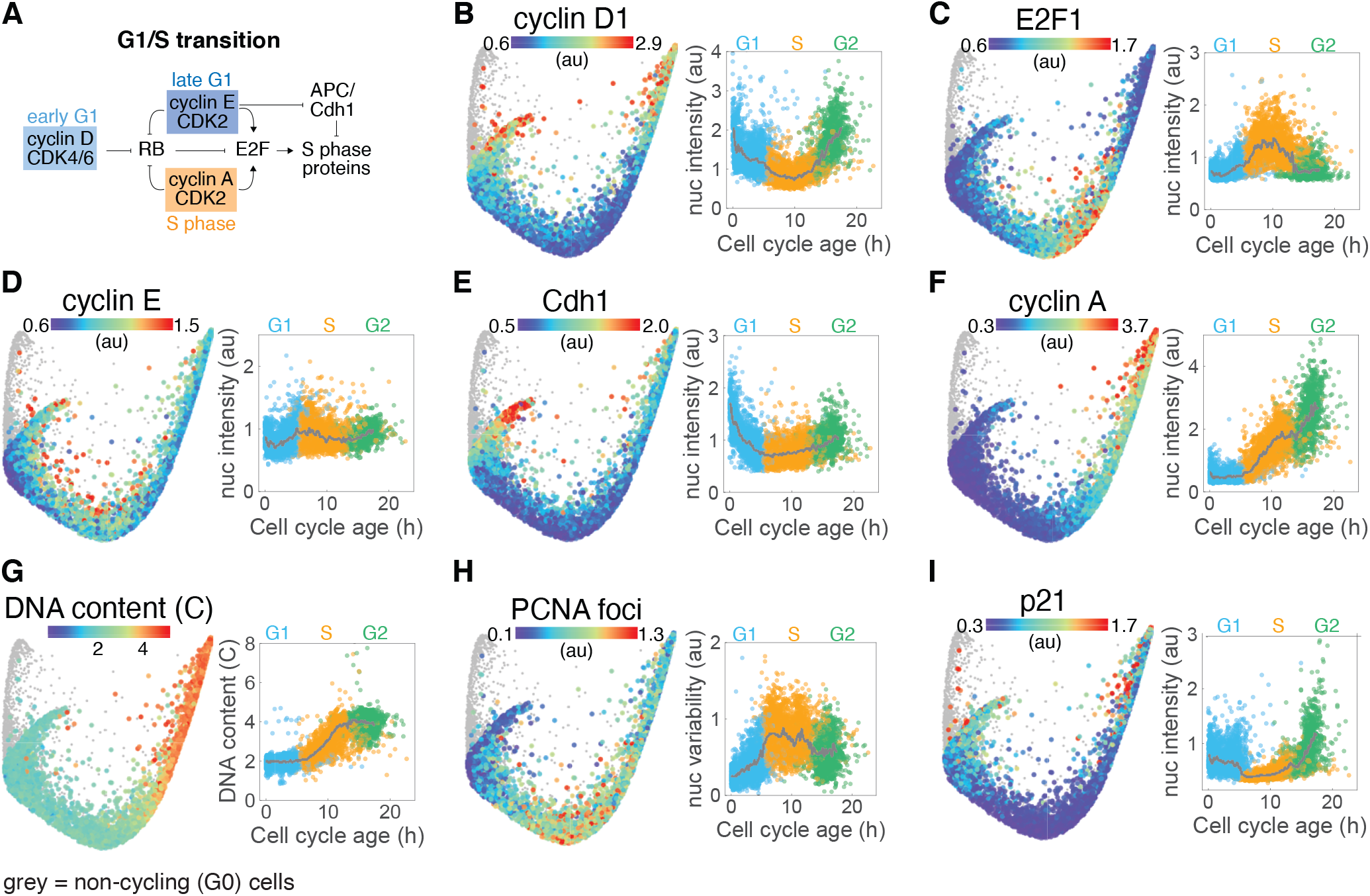
Visualizing the mechanisms of the G1/S transition. (**A**) Mechanistic model of the core events regulating progress through G1 and the transition to S. (**B-I**) Median nuclear intensity of cyclin D1 (B), E2F1 (C), cyclin E (D), Cdh1 (E), cyclin A (F) and p21 (I), DNA content (copy number) (G), and variability in nuclear PCNA intensity (H) of cells in the proliferative trajectory are mapped onto the cell cycle structure (left panels) and plotted against cell cycle age (right panels). Population medians in time courses indicated by solid grey lines and individual cells are colored by cell cycle phase (G1: blue, S: orange, G2: green). Non-cycling (G0) cells (phospho/total RB < 1.6) are shown in grey on the structure and are excluded from time courses.

DNA replication was clearly visible on the structure, with DNA content doubling over the course of S phase (**Fig. 2G**). Similarly, we localized additional S phase events important for DNA replication including the appearance of PCNA foci at replication complexes (quantified as variability in nuclear intensity) (**Fig. 2H**) and the replication-coupled destruction of p21 (**Fig. 2I**). Upon entry into G2, cyclin B expression increases first in the cytoplasm and then in the nucleus (**fig. S3A-B**) as cells move toward mitosis (*37*). We also observed increases in cyclin D1 (**Fig. 2B**) and its transcription factor cMyc (**fig. S3C**) as well as NF-kB activation (**fig. S3D**) during G2, which then remained elevated through mitosis and into the subsequent G1 phase of daughter cells, resulting in U-shape dynamics along the proliferative trajectory (*27*). Taken together, this approach accurately recapitulated many of the key molecular events governing the progression of cells through the four proliferative cell cycle phases.

### Trajectories into and out of cell cycle arrest (G0)

We used the same approach to examine the molecular changes that occur as cells progress along the arrest trajectory (**Auxiliary fig. S2**), for which comparatively less is known. We noted that cells exited mitosis with high RB phosphorylation along a single path down the central arm and then quickly diverged along two distinct trajectories within the first 2-3 hours following mitosis: either progressing directly into G1 with sustained RB phosphorylation or exiting the cell cycle into G0 following the loss of RB phosphorylation (**Fig. 3A, Fig. 1F, fig. S4A**). In each daughter cell, this fate decision was found to be governed primarily by the balance between cyclin D and p21 expression. Cyclin D and p21 activate and inhibit CDK4/6 activity, respectively, to control the phosphorylation state of RB in a stoichiometric and ultrasensitive manner (**fig. S4A-D**) (*13, 14, 38*). We found that the decrease in the cyclin D1:p21 ratio that precedes cell cycle exit was not due to a decrease in cyclin D1 expression, but rather an increase in p21 expression in early G1 (**Fig. 3B, fig. S4A**), consistent with previous observations (*38*). This induction of p21 in early G1 occurred simultaneously with the loss of RB phosphorylation as cells diverted to the arrest trajectory (**fig. S4A**).

**Figure 3.**
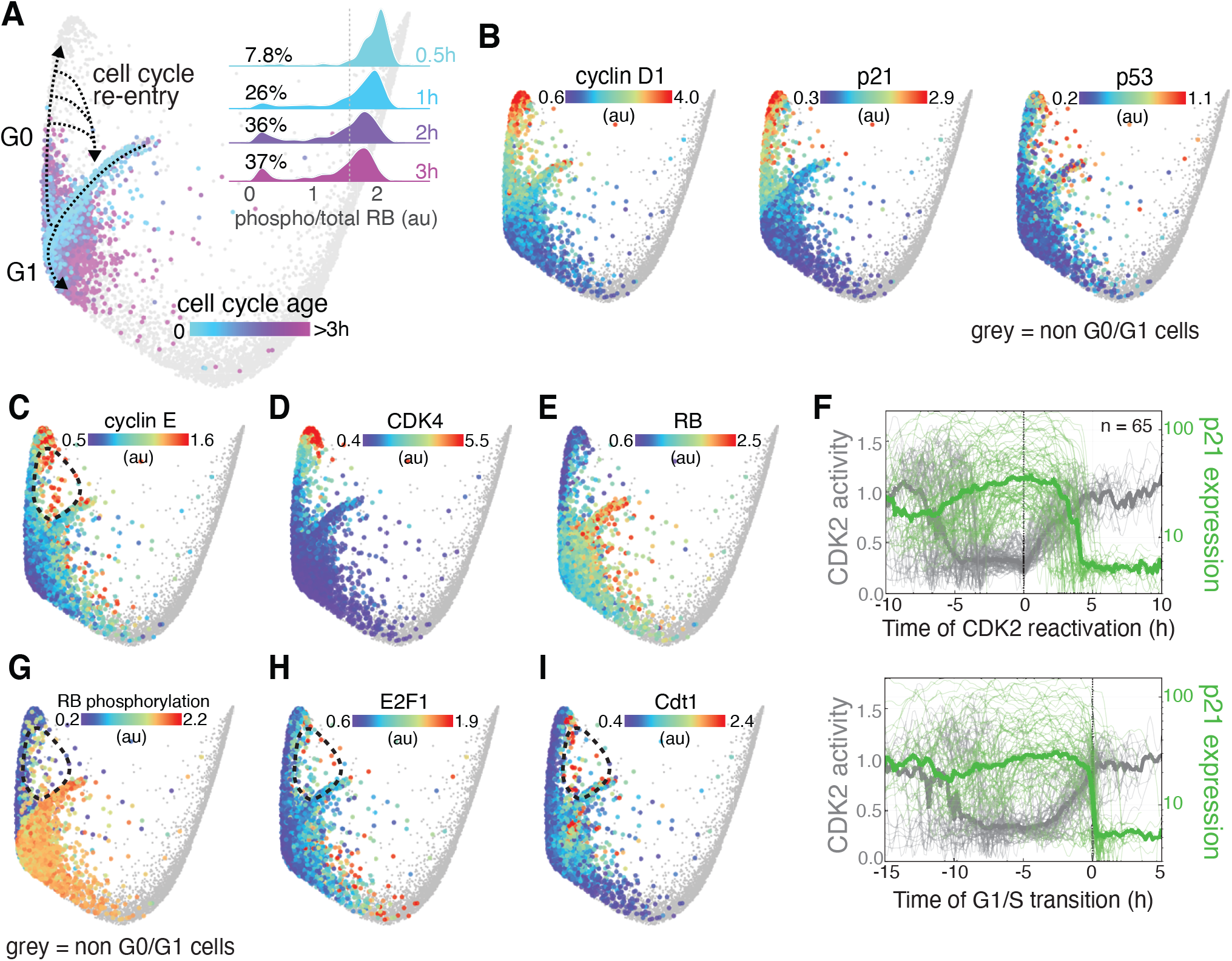
Cell cycle exit and re-entry from spontaneous arrest (G0). (**A**) Divergence of cell cycle trajectories into G1 or G0 following mitosis and subsequent cell cycle re-entry from G0. Cell cycle age of newborn cells (< 3h since mitosis) is mapped onto the structure. *Inset:* Time course of RB phosphorylation in individual cells following mitosis. Cells were binned by cell cycle age (less than 0.5, 1, 2 and 3 h post-mitosis). (**B**) Median nuclear expression of cyclin D1 (left), p21 (middle) and p53 (right) mapped onto G0/G1 cells. (**C-E**) Median nuclear expression of cyclin E (C), CDK4 (D) and RB (E) mapped onto G0/G1 cells. (**F**) Single cell traces from time-lapse imaging of CDK2 activity (grey, quantified by the cytoplasm:nuclear ratio of DHB-mCherry) and p21-YPet expression (green) aligned at the time of CDK2 reactivation (upper panel) and the G1/S transition (lower panel). Thick lines represent population medians. *N* = 65 cells. (**G-I**) RB phosphorylation (G) and median nuclear expression of E2F1 (H) and Cdt1 (I) mapped onto G0/G1 cells. Note the dotted circle representing cell cycle re-entry in C, G, H and I.

Further examination of the arrest trajectory revealed that cells do not exit the cell cycle into a single, static arrest state awaiting cell cycle re-entry. Instead, cells progressed further along this trajectory with time (**Fig. 1D**) accompanied by a progressive change in molecular signature suggesting an increasing depth of arrest (*18, 19, 22, 27*). Cells with longer durations of arrest exhibited increases in the expression of p21, p53 (**Fig. 3B**), p38 and p16 (**fig. S5**). Surprisingly, progression further up the arrest arm was also accompanied by increases in cyclin D1 (**Fig. 3B**), cyclin E, and CDK4 expression, along with loss of RB expression (**Fig. 3C-E**), molecular changes that typically promote cell cycle progression, not arrest.

We therefore hypothesized that cells may re-enter the cell cycle not by reversing the increase in p21 expression that drove cell cycle exit but instead by overcoming it with excessive cyclin and/or CDK expression. To test this hypothesis, we performed time lapse imaging of RPE cells engineered using a CRISPR-Cas9 knock-in approach to express fluorescently-tagged endogenous p21 (p21-YPet), a marker of cell cycle phase (PCNA-mTq2) and a CDK2 activity sensor (DHB-mCherry) to monitor cell cycle exit and re-entry (*12*). We observed that the re-activation of CDK2 as cells emerged from arrest occurred prior to a reduction in p21 (**Fig. 3F**). Instead, a decrease in p21 expression was observed only later, as cells transitioned into S phase due to its replication-coupled destruction (**Fig. 3F**) (*15*). Furthermore, the majority of cells (83%) re-entered the cell cycle (as identified by the reactivation of CDK2) at equal or higher p21 expression than was required to inhibit CDK2 activation during cell cycle exit immediately after cell division (**fig. S4E**), indicating that a reduction in p21 expression to baseline is not a prerequisite for cell cycle re-entry and that the CDK activity required to progress through G1 and into S phase can be achieved instead by mechanisms that overcome elevated p21 expression.

Cells re-enter the cell cycle along a funnel-shaped trajectory on the inside of the structure, connecting various points along the arrest arm with the base of the central arm (**Fig. 3A**). Cells in this region showed increased expression of cyclin D1 (**Fig. 3B**), a high cyclin D1:p21 ratio (**fig. S4C**) and high RB phosphorylation (**Fig. 3G, Movie S4**), consistent with cell cycle re-entry. Having spent variable amounts of time in G0, these cells are relatively old compared to other G1 cells (**Fig. 1D, Movie S2**), and exhibit characteristics of cells preparing for DNA replication including increased expression of cyclin E (**Fig. 3C**), E2F1 (**Fig. 3H**) and Cdt1 (**Fig. 3I**). Cell cycle re-entry therefore is not simply a reversal of the mechanisms that drive cells into arrest, but instead occurs along a distinct molecular trajectory back to the proliferative cell cycle.

### Cellular senescence

In addition to changes in their molecular state, cells also increased in size as they progressed further along the arrest trajectory with time (**fig. S6A, Fig. 1D**), a phenomenon first observed in yeast cell cycle arrest (*39*). In particular, cells residing at the end of the arrest trajectory (**Fig. 4A**) were at least 3-4 times larger than the average arrested cell and possessed a high cytoplasm-to-DNA ratio (**Fig. 4B**), a hallmark of cellular senescence (*40*). Cells in this terminal arrest region also showed the longest durations of arrest (>18 h, **Fig. 1D**) and exhibited a unique molecular signature that was distinct from other cells in the arrest arm. This molecular profile included increased expression of DNA damage markers phospho-H2AX, phospho-CHK1, p38 (**fig. S6B-D**) and p53 (**Fig. 4C**), CDK inhibitors p21, p27 and p16 (**fig. S6E-G**), and loss of the proliferative effectors PCNA and Fra1 (**fig. S6H-I**). These cells also exhibited a cytoplasmic dilution of CDK2 (**Fig. 4D**) and increased nuclear expression of β-catenin (**Fig. 4E**), an effector known to mediate signaling during senescence induction (*41*). Cells located in this region possessed either 2 or 4 copies of DNA (2C or 4C, **Fig. 4F**) (and no multinucleated cells were observed), indicating that some cells must have exited the proliferative cell cycle along a second arrest trajectory that diverts after DNA replication (**Fig. 4A**). Indeed, we identified a population of cells residing along a path between G2 and the arrest terminus with 4 copies of DNA (**Fig. 4F, Movie S1**), elevated markers of a DNA damage response (**Fig. 4C, fig. S6B-E**) and a loss of RB phosphorylation (**Fig. 1F**).

**Figure 4.**
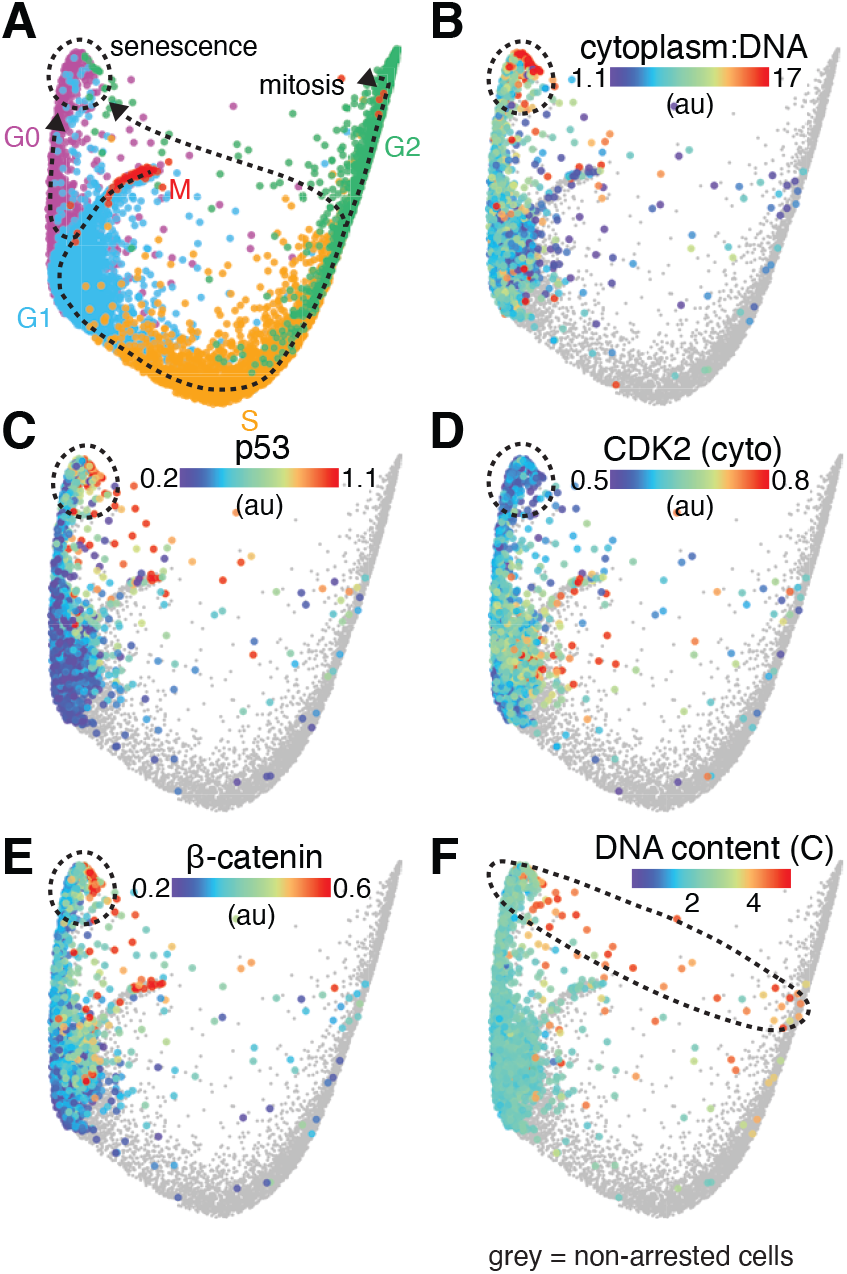
Signature of cellular senescence. (**A**) Model showing two distinct molecular trajectories into senescence, diverging from the proliferative cell cycle in either G1 or G2. (**B-F**) Ratio of cytoplasmic area:DNA content (B), median nuclear expression of p53 (C) and β-catenin (E), median cytoplasmic expression of CDK2 (D) and DNA content (copy number) (F) of arrested cells are mapped onto the structure. Non-arrested cells (phospho/total RB > 1.6) are shown in grey.

### The molecular architecture of cell cycle arrest

To further explore the architecture of cell cycle arrest, we performed 4i and manifold learning on >23,000 cells following treatment with a variety of stresses known to induce cell cycle arrest: hypomitogenic stress induced by serum starvation, replication stress following etoposide treatment and oxidative stress in response to hydrogen peroxide (**Fig. 5**). Plotting the arrest trajectories in response to each of these stresses on the same map alongside unperturbed cells provided a clear visualization of the complex architecture of cell cycle arrest and its connectivity with the proliferative cell cycle (**Fig. 5A**).

**Figure 5.**
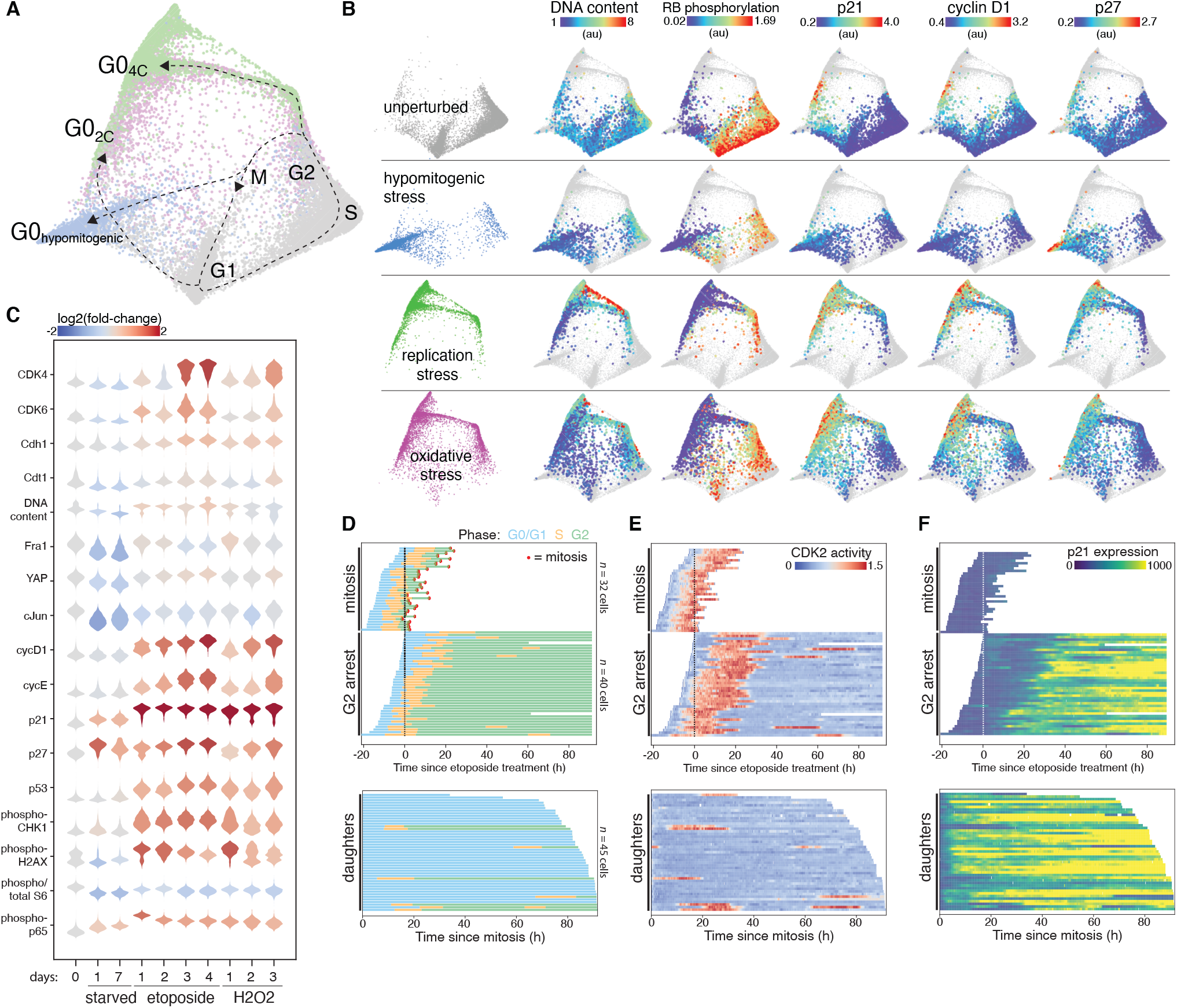
The architecture of cell cycle arrest. (**A**) Expanded cell cycle structure depicting the proliferative phases (G1/S/G2/M) and multiple states of arrest (G0). Unperturbed (grey, *N* = 11,268 cells), serum-starved (“hypomitogenic stress”, 1 or 7 days, blue, *N* = 3007 cells), etoposide-treated (“replication stress”, 1 μM for 1-4 days, green, *N* = 4315 cells) and H2O2-treated (“oxidative stress”, 200 μM for 1-3 days, purple, *N* = 5015 cells) cells are mapped onto the structure. (**B**) DNA content, RB phosphorylation and median nuclear expression of p21, cyclin D1 and p27 are mapped onto unperturbed and stressed cells. For panels A and B, all time points for each treatment are mapped on the structure. (**C**) Molecular signatures of cell cycle stresses over time. Single cell distributions are colored by relative expression versus control. (**D-F**) Single cell tracks derived from time lapse imaging depicting cell cycle phase (D, manually annotated based on presence of PCNA foci in S phase), CDK2 activity (E, quantified by the cytoplasm:nuclear ratio of DHB-mCherry) and p21-YPet expression (F, median nuclear intensity) following 1 μM etoposide treatment. G2 arrested and mitotic fates (upper panels) and daughters of mitotic cells (lower panels) are shown. *N* = 117 cells.

In response to hypomitogenic stress (i.e. serum starvation for 1 or 7 days), cells diverged from the proliferative cell cycle in G2, undergoing cell division directly into a state of arrest with 2C DNA content (“G0_hypomitogenic_”, **Fig. 5A**) identified by the loss of RB phosphorylation (**Fig. 5B**). However, unlike the spontaneous arrest observed in unperturbed cells (**Fig. 3B**), exit to the hypomitogenic arrest state was accompanied by only a modest increase in p21, and instead was driven primarily by a loss of cyclin D1 expression (**Fig. 5B, fig. S7**). Following serum starvation, cells do not exhibit the characteristic increase in cyclin D1 during G2 that is observed in the cell cycle of unperturbed cells (**Fig. 5B, Fig. 2B**). Consequently, RB phosphorylation is rapidly lost due to a lack of cyclinD1:CDK4/6 activity (*16*) as cells complete mitosis. Progression further along this trajectory was accompanied by an increase in p27 (**Fig. 5B**) and loss of CDK4 and CDK6 (**fig. S8**), consistent with previous observations that hypomitogenic arrest is not a single static state but rather a continuum of states of deeper quiescence (*18, 19, 22*). The hypomitogenic arrest state was also characterized by decreased expression of key proliferative effectors Cdh1, Cdt1, Fra1 and cJun, as well as decreased nuclear YAP and mTOR signaling (S6 phosphorylation) (**Fig. 5C, fig. S8**).

We next assessed how the cell cycle responds to replication stress following treatment with etoposide (for 1, 2, 3 or 4 days), an inhibitor of DNA topoisomerase II that interferes with DNA re-ligation during replication. Within a single population of cells treated with etoposide, we observed that individual cells exited the proliferative cell cycle along two distinct arrest trajectories: (1) from a G2-like state after DNA replication was complete (DNA content = 4C), or (2) in the subsequent G1 phase of daughter cells immediately following mitosis (DNA content = 2C) (**Fig. 5A**). Cell cycle exit to the 4C state (“G0_4C_”) was accompanied by the activation of the DNA damage checkpoint in G2, including an increase in DNA damage itself (pH2AX), activation of CHK1, NF-κB signaling (phospho-p65) and increased expression of p53 (**fig. S9**) and p21 (**Fig. 5B, fig. S7**), while daughter cells that exit to the 2C state following mitosis (“G0_2C_”) possessed elevated p21 expression but showed no increase in other DNA damage markers (**Fig. 5B, fig. S9**). While a few unperturbed cells were found to populate both the hypomitogenic and 4C states of arrest, the majority of spontaneously arrested cells (with low RB phosphorylation) were observed along a trajectory towards the 2C arrest state driven by an increase in p21 but exhibiting very weak expression of DNA damage markers (**Fig. 5B, fig. S9**), consistent with observations that spontaneous cell cycle arrest results from low levels of replication stress (*38*).

While the PCNA-mTq2 reporter allows delineation the cell cycle along two distinct arrest trajectories in response to replication stress and to investigate the mechanisms that govern this fate decision, we performed single cell time lapse imaging of RPE cells expressing a cell cycle sensor (PCNA-mTq2), a CDK2 activity sensor to detect cell cycle arrest (DHB-mCherry) (*12*) and endogenous p21 fused to a fluorophore (p21-YPet) for ∼4 days after etoposide treatment (**Fig 5D-F, fig. S10**). We observed a similar bifurcation of cell fate in live cells in response to replication stress, with 56% of cells arresting in G2 and 44% proceeding through to mitosis following etoposide treatment (**Fig. 5D, upper panel**). For each of these cells that successfully completed mitosis, however, we observed that their daughter cells arrested immediately following cell division (as indicated by a sustained decrease in CDK2 activity) (**Fig. 5D, lower panel**). Choice between these two fate trajectories was dependent on the timing of p21 induction in individual cells. While G2 arrest coincided with a loss of CDK2 activity and an increase in p21 expression soon after the S/G2 transition (**Fig. 5E-F, upper panel; fig. S10A**), in cells that proceeded through to mitosis, p21 remained low in G2 (**Fig. 5F, upper panel; fig. S10B**) but increased abruptly following cell division in daughter cells (**Fig. 5F, lower panel; fig. S10C**). Furthermore, daughter cell cycle arrest was found to be reversible, with some cells re-entering the cell cycle and transitioning into S phase after variable lengths of time in G0 (**Fig. 5D, bottom panel**). These cells, however, invariably arrested as they transitioned into G2, due to the persistent stress of etoposide during DNA replication, consistent with the observation that cells progressively escaped from the 2C arrest state and accumulated in the 4C state over time following etoposide treatment (**Fig. 6A-B**).

**Figure 6.**
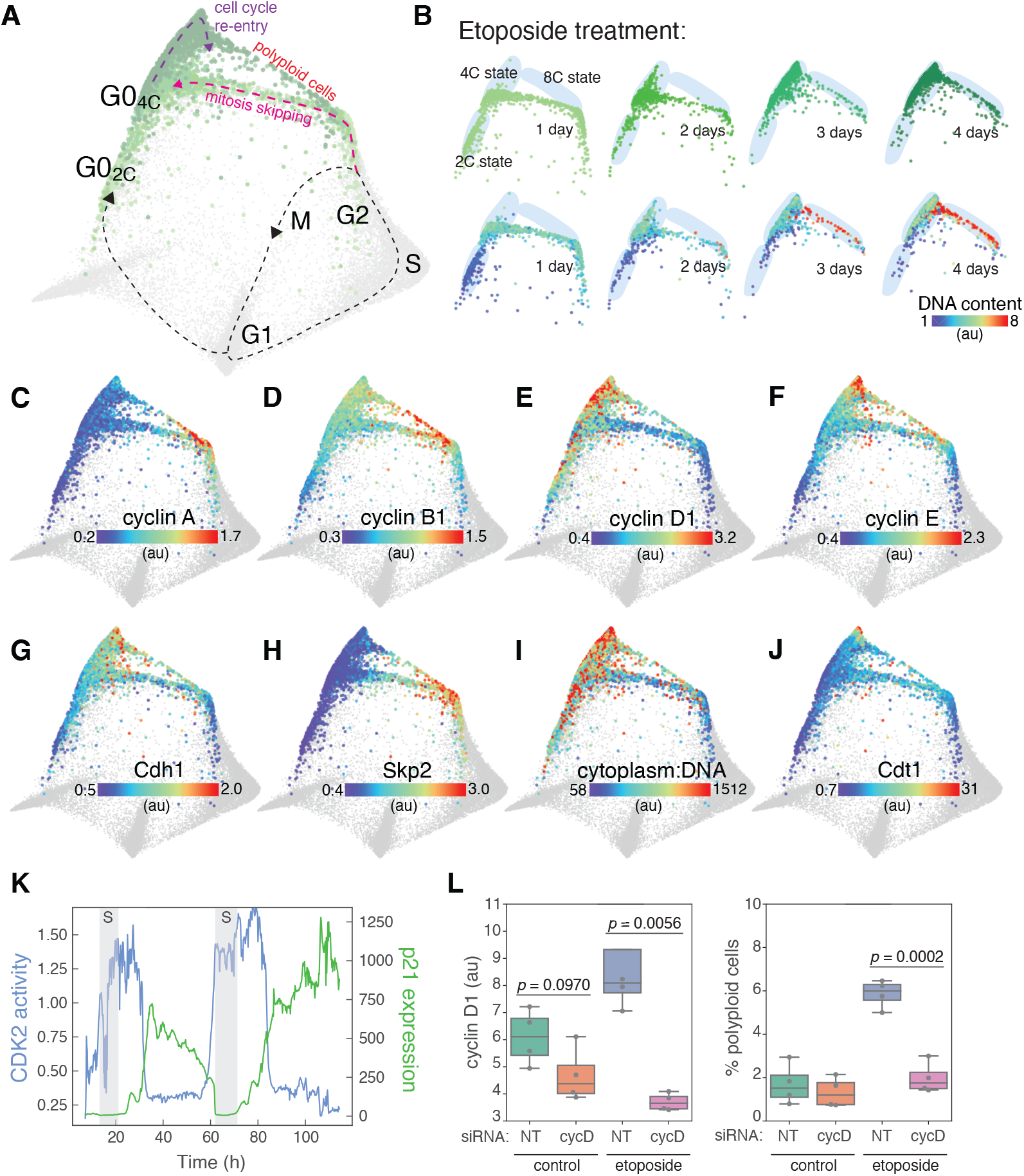
Mitosis skipping and polyploidization. (**A**) Cell fate trajectories in response to replication stress. Unperturbed (grey) and etoposide-treated (1 μM, green) cells are mapped onto the structure. (**B**) Changes in the proportion of cells with 2C, 4C and 8C DNA content over time following etoposide treatment. (**C-J**) Median nuclear intensities of cyclin A (C), cyclin B1 (D), cyclin D1 (E), cyclin E (F), Cdh1 (G), Skp2 (H) and Cdt1 (J) and the cytoplasm:DNA ratio (I) of etoposide-treated cells are mapped onto the expanded cell cycle structure. Data from all time points (1-4 days) are plotted together on each structure. (**K**) Representative single cell trace showing mitotic skipping and endoreduplication observed during time lapse imaging. Two S phases were identified by the appearance of PCNA nuclear foci and are depicted by grey bars. CDK2 activity (quantified by the cytoplasm:nuclear ratio of DHB-mCherry, blue) and p21-YPet expression (green) are shown. (**L**) Cyclin D1 expression (left) and the proportion of polyploid cells (right) measured by Hoechst staining and immunofluorescence, respectively, following siRNA-mediated knockdown of cyclin D in control and etoposide pre-treated cells. Bars represent population medians from four independent replicates (grey circles) and statistical significance was determined using a two-tailed Student’s *t* test.

To assess the cell cycle response to oxidative stress, we treated cells with hydrogen peroxide (H_2_O_2_) for 1, 2 or 3 days. We observed very similar arrest architecture to replication stress (**Fig. 5A-B**), indicating that the dominant cell cycle response to exogenous oxidative stress is primarily related to its known ability to induce DNA damage (**Fig. 5B-C, fig. S9**)(*42*). Similar to etoposide treatment, H_2_O_2_ induced cell cycle exit along two distinct trajectories diverging from either G1 or G2, into arrest states with 2C or 4C DNA content, respectively, both accompanied by p21 induction (**Fig. 5B, fig. S7**). However, unlike etoposide, H_2_O_2_ is rapidly metabolized following addition to cells (*43*) and consequently induces only temporary stress soon after treatment. Consistent with a transient DNA damage response, markers of DNA damage decrease more rapidly (**Fig 5C, fig. S11A-B**) and a higher proportion of cells remain in the cell cycle over time (**fig. S11C-D**).

### Mitosis skipping and polyploidization

While the 2C state of cell cycle arrest following etoposide-induced replication stress was found to be reversible (**Fig. 5D-E, bottom panels**), the accumulation of cells in the 4C state over time (**Fig. 6A-B**) suggested that it may represent an irreversible state of cell cycle arrest. As cells progressed along the arrest trajectory from G2 to the 4C arrest state (**Fig. 6A, pink arrow**), there was an abrupt degradation of the S/G2 cyclins A and B (**Fig. 6C-D**) that coincided with the loss of RB phosphorylation (**Fig. 5B**). Degradation of these cyclins normally occurs during mitosis (*44*). However, we observed no change in DNA content (**Fig. 6B**) nor any visual evidence of mitotic events in the images of cells along this trajectory. Progression further along this trajectory was accompanied by an increase in G1 cyclins D1 and E, as well as an increase in Cdh1 and a loss of Skp2 (**Fig. 6E-H**), all consistent with the transition into a G0/G1-like state and a phenomenon known as “mitotic skipping”, which often precedes the transition into senescence (*45*–*47*). Indeed, after 3 days of etoposide treatment, most cells resided in a G0/G1-like state of cell cycle arrest with 4C DNA content (**Fig. 6B**) and a high cytoplasm-to-DNA ratio (**Fig. 6I**), consistent with a senescent state (*40, 47*).

However, following a trajectory upward from the 4C arrest state (**Fig. 6A**, *purple arrow*), we observed molecular changes consistent with cell cycle re-entry and progress toward S phase, including sequential increases in cyclin D1, cyclin E and the DNA licensing factor Cdt1 (**Fig. 6E,F,J**). Furthermore, there was a gradual accumulation of polyploid cells with 8C DNA content over time following etoposide treatment (**Fig. 6B**), supporting a hypothesis that cells might be able to re-enter the cell cycle from this 4C state of arrest and undergo a second round of DNA replication, or “endoreduplication” (*48*), without prior cell division. To validate these observations, we performed time lapse imaging for 4 days following etoposide treatment. We observed rare instances of cells re-entering the cell cycle from the 4C state of arrest (as indicated by an increase in CDK2 activity) and transitioning into a second S phase (as indicated by an increase in PCNA foci) without undergoing mitosis (5/117 cells over 4 days of imaging; **Fig. 6K, Fig. 5D**). Similar to cell cycle re-entry in unperturbed cells (**Fig. 3**), the re-emergence of cells from the 4C state of arrest following mitotic skipping occurred despite elevated p21 expression (**Fig. 5B,6K**) in cells with very high expression of cyclin D1 (**Fig. 6E**). To test if increased cyclin D expression can overcome p21 expression and drive endoreduplication, we treated cells with siRNA against all three cyclin D isoforms following etoposide treatment. Knockdown of cyclin D completely abolished the appearance of polyploid cells in response to replication stress (**Fig. 6L**), confirming its role in the generation of polyploidy, and indicating that increased expression of G1 cyclins may represent a general mechanism through which cells can escape p21-induced cell cycle arrest.

## Discussion

We combined time lapse microscopy, multiplexed imaging, and manifold learning to reveal the underlying structure of the human epithelial cell cycle. This data-driven approach revealed the molecular architecture of proliferative and arrested states as well as a detailed map of the connectivity between them.

Under standard cell culture conditions, we observed a single proliferative trajectory through DNA replication and cell division. The primary source of heterogeneity observed among unperturbed cells occurred shortly after cell division, when the molecular trajectories of newborn daughter cells diverged along two distinct paths, either immediately re-entering the cell cycle or diverting into a spontaneous arrest state driven by an induction of p21 expression (**Fig. 3**), as previously described (*12, 14, 32, 49*). Although an alternative mechanistic route through G1 has recently been observed when the canonical route is blocked (i.e. by CDK4/6 inhibition)(*8*), our results indicate that, when grown under ideal culture conditions, cells primarily traverse a single trajectory *en route* to mitosis. This topology was reproducibly obtained from replicate cell populations, across PHATE parameter space and in a second human epithelial cell line (**fig. S2**).

We mapped the architecture of cell cycle arrest states using various cell cycle stresses, revealing multiple molecular trajectories diverging from different points in the proliferative cell cycle into distinct arrest states (**Fig. 5**). In response to mitogen starvation, for example, cells followed a graded trajectory into a single arrest state characterized by a lack of cyclin D and an increase in p27. In contrast, replication or oxidative stress caused cells to exit the proliferative cell cycle along two distinct arrest trajectories: diverting either in G2 or immediately after cell division. The choice between these divergent arrest trajectories was determined by distinct p21 dynamics in response to DNA damage.

In addition, we found that increased expression of proliferative effectors such as cyclin D1, cyclin E and CDK4/6 was a general feature of all arrest states driven by p21 induction, promoting cell cycle re-entry from the spontaneous arrest observed in control cells (**Fig. 3**), the 2C arrest state following replication and oxidative stress (**Fig. 4**) and from the 4C arrest state that follows mitotic skipping, which in this particular case leads to endoreduplication and polyploidization (**Fig. 6**). As a general principle, these findings indicate that cell cycle re-entry does not necessarily require a reversal of the mechanisms that induced arrest, but can occur instead along a distinct molecular trajectory, in some cases by overcoming sustained arrest signals with countervailing proliferative signals.

The systems-level approach described in the current study allows many of the core regulatory events governing cell cycle progression and arrest to be quantified and visualized simultaneously on a single representation and using a single assay. Here, we demonstrated how this approach can identify how the cell cycle responds to different stresses. This comparative cell cycle analysis may prove particularly powerful in the context of cancer cell biology. Identifying differences between normal and oncogenic cell cycle structures may provide novel insights into the mechanisms of tumorigenesis and lead to the development of new therapeutic targets.

## Supporting information

Movie S1. DNA content mapped onto the 3-dimensional structure of the cell cycle.

Movie S2. Cell cycle age mapped onto the 3-dimensional structure of the cell cycle.

Movie S3. Cell cycle phase mapped onto the 3-dimensional structure of the cell cycle.

Movie S4. RB phosphorylation status mapped onto the 3-dimensional structure of the cell cycle.

Auxiliary fig S1. Effector dynamics along the proliferative trajectory

Auxiliary fig S2. Effector dynamics along the arrest trajectory

## Acknowledgments

We would like to thank Sonja Mihailovic (UNC Chapel Hill), Chi Pham (UNC Chapel Hill) and Margaret Redick (Oregon State University) for assistance with cell culture and Dr. Sam Wolff (UNC Chapel Hill) for support with imaging. We would like to thank Dr. Robert Duronio (UNC Chapel Hill) and Dr. Adam Palmer (UNC Chapel Hill) for careful reading of the manuscript.

## Authors contributions

WS and JEP conceived of the project. WS, KMK, JGC and JEP designed the experiments. WS, HKS, SRT and CLY performed the 4i experiments. WS and SRT performed the time lapse imaging. WS, KMK and CDT performed image analysis. TMZ trained the random forest models. KMK trained the neural networks. HKS and CLY performed Western blots. JCL created the RPE-dox-KRAS cell line. WS wrote the manuscript with the help of all authors.

## Data and materials availability

Single cell datasets, including data compatible for exploration in virtual reality, are available at http://doi.org/10.5281/zenodo.4525425.

## Supplementary Materials

Materials and Methods

Table - S2

Fig S1 - S12

Auxiliary Figures S1 - S2

Movies S1 - S4

## Materials and Methods

### Cell culture

Retinal pigment epithelial cells (hTERT RPE-1, ATCC, CRL-4000) were cultured in DMEM (Gibco, 11995-065) supplemented with 10% fetal bovine serum (FBS; Sigma, TMS-013-B), 2 mM L-glutamine (ThermoFisher Scientific, 25030081) and penicillin/streptomycin (P/S; ThermoFisher Scientific, 15140148). For time lapse imaging of RPE cells, FluoroBrite™ DMEM (Gibco, A18967-01) supplemented with 10% FBS and 2 mM L-glutamine was used. and Human pancreatic epithelial cells (hTERT-HPNE, ATCC, CRL-4023) were cultured in pyruvate-free DMEM (Gibco, 11965-092) supplemented with 10% FBS and P/S. All cells were cultured at 37 °C and 5% CO_2_.

### Cell line generation

The construction of the RPE-PCNA-mTurqoise2 (RPE-mTq2) cell line was previously described (*7*). The RPE-PCNA-mTq2/p21-YPet/DHB-mCherry cell line was engineered using a CRISPR-Cas9 knock-in approach to introduce the YPet fluorophore into the 3’ region of the endogenous p21 gene in a RPE-PCNA-mTq2/DHB-mCherry cell line (*50*). We synthesized the gene sequence for YPet-P2A-neoR (**fig. S12**) flanked by 900-950 bp homology arms into a pUC donor plasmid (Bio Basic Inc.) and targeted it to the endogenous p21 gene with the gRNA sequence ggaagccctaatccgcccac (Synthego). Recombinant Cas9 2.0 (Thermo Fisher Scientific, A36498) was mixed with gRNA and incubated for 15 min at room temperature. A total of 3×10^5^ cells were electroporated with 900 ng sgRNA and 3000 ng linearized donor plasmid using the Neon Transfection System (ThermoFisher Scientific). Positive clones were enriched using a low-dose of G418 (200 μg/ml), and individual clones were hand-picked and screened for successful fluorophore integration following overnight neocarzinostatin treatment to induce p21-YPet expression. The RPE-dox-KRAS cell line used in the cell stress 4i experiment was constructed by introducing the pInducer20 plasmid containing KRAS-G12V cDNA into RPE cells by viral transduction as described above. All cell lines were authenticated by STR profiling (ATCC) and confirmed to be mycoplasma free.

### Antibodies

High quality, previously published/validated primary antibodies were identified using BenchSci (http://app.benchsci.com) and are listed in Table S1.

### Time lapse imaging

Cells were plated in glass-bottom plates (Cellvis) coated with fibronectin (1 μg/cm^2^, Sigma, F1141). Fluorescence images were acquired using a Nikon Ti Eclipse inverted microscope with a Nikon Plan Apochromat Lambda 40x objective with a numerical aperture of 0.95 and an Andor Zyla 4.2 sCMOS detector. Autofocus was provided by the Nikon Perfect Focus System (PFS) and a custom enclosure (Okolabs) was used to maintain constant temperature (37°C) and atmosphere (5% CO_2_). For time lapse imaging, the following filter sets were used (excitation; beam splitter; emission filter; Chroma): CFP (425-445/455/465-495nm), YFP (490-510/515/520-550nm) and mCherry(540-580/585/593-668). Stitched 4-by-4 images were acquired every 10 min for RPE-PCNA-mTq2/p21-YPet/DHB-mCherry cells and every 16 min for RPE-PCNA-mTq2 cells. Uneven field illumination was corrected prior to stitching. NIS-Elements AR software was used for image acquisition and post-processing. CDK2 activation was quantified as the ratio of background corrected cytoplasmic to nuclear intensity of the DHB-mCherry sensor (cytoplasm signal quantified as a 40th percentile in a 15-pixel ring outside the nuclear segmentation, with a 2-pixel gap between the nucleus and the ring; nuclear signal quantified as median) and p21-YPet expression was calculated as the background corrected median nuclear intensity. Cell cycle phases were annotated manually from time lapse imaging using the appearance/disappearance of nuclear PCNA foci to mark the beginning and end of S phase, respectively.

For the time lapse imaging that preceded 4i (**see Fig. 1A**), approximately 25% of the total well area was imaged for a total of 24h, permitting ∼27% of the total cells to be tracked. Nuclear regions were segmented based on the PCNA-mTq2 signal using a modified U-Net neural network (https://github.com/fastai/fastai). Linking of regions into tracks was computed using TrackMate (*51*). Segmentation and tracking corrections were performed manually.

### Iterative indirect immunofluorescence imaging (4i)

Cells were plated in glass-bottom plates (Cellvis) coated with fibronectin (1 μg/cm^2^, Sigma, F1141), treated as required and prepared as follows. In between each step, samples were rinsed 3X times with phosphate-buffered saline (PBS) and incubations were at room temperature, unless otherwise stated. Cells were fixed with 4% paraformaldehyde (ThermoFisher Scientific, 28908) for 30 min, permeabilized with 0.1% Triton X-100 in PBS for 15 min and inspected for sample quality control following Hoechst staining in imaging buffer (IB: 700 mM N-acetyl-cysteine (Sigma, A7250) in ddH_2_O. Adjust to pH 7.4). Sample was rinsed 3X with ddH_2_O and incubated with elution buffer (EB: 0.5M L-Glycine (Sigma, 50046), 3M Urea (Sigma, U4883), 3M Guanidine chloride (ThermoFisher Scientific, 15502-016), and 70mM TCEP-HCl (Sigma, 646547) in ddH_2_0. Adjusted to pH 2.5) 3X for 10 min on shaker to remove Hoechst stain. Sample was incubated with 4i blocking solution (sBS: 100 mM maleimide (Sigma, 129585), 100 mM NH_4_Cl (Sigma, A9434) AND 1% bovine serum albumin in PBS) for 1h and incubated with primary antibodies diluted as required (**Table S1**) in conventional blocking solution (cBS: 1% bovine serum albumin in PBS) overnight at 4°C. Samples were rinsed 3X with PBS and then incubated in secondary antibodies (**Table S1**) and Hoechst for 1h on shaker, then rinsed 5X with PBS and imaged in IB. Samples were imaged using the Nikon Ti Eclipse microscope described above. Stitched 8×8 images were acquired for each condition using the following filter cubes (Chroma): DAPI(383-408/425/435-485nm), GFP(450-490/495/500-550nm), Cy3(530-560/570/573-648nm), Cy5(590-650/660/663-738nm). After imaging, samples were rinsed 3X with ddH2O, antibodies were eluted and re-stained iteratively as described above.

### Image processing and single cell analysis

Image registration was performed using a custom Python script (v3.7.1) using features common to multiple rounds (Hoechst or CDK2 staining) and StackReg library (*52*). Segmentation and feature extraction from registered images were performed using standard modules in CellProfiler (v3.1.8). Only cells that persisted through all rounds of 4i were included in subsequent analyses.

### Random forest models

Two random forest (RF) models were trained to predict cell cycle age and phase (obtained by time lapse imaging) from the multivariate 4i signatures of the same cells. 80% of our annotated data was used to train the RF and the remaining 20% was reserved as a test set. Classification accuracy was used as the error metric to train the phase model, while root mean squared error (RMSE) was used to train the age model. The phase model yielded 95.5% accuracy (95% CI: 93.5, 0.971) and a kappa of 0.925. The age model had an RMSE of 125.8 with an R^2^=0.862. Variable importance tables were obtained from each model (**fig. S2A, Table S2**) through calculation of unconditional permutation importance. All hyperparameter tuning was performed with 10-fold cross validation (Age: ntrees=500, mtry=246, Phase: ntrees=500, mtry=124).

To identify the optimal feature subset that best predicts the cell cycle state (i.e. age and phase), first a combined ranking was calculated for each feature as the average of the individual rankings from the variable importance tables obtained from the age and phase models. Successive random forest models were generated for both age and phase starting with a single feature and adding additional features in order of combined ranking. Accuracy (for cell cycle phase) or error (RMSE for cell cycle age) were calculated at each iteration (**fig. S2B-C**). The optimal feature was defined as the top 40 features (by combined ranking).

All RF analyses were performed using R (v1.2.5001) with caret (v6.0-86) and ggplot2 (v3.3.2) packages.

### Convolutional neural network

The convolutional neural network (CNN) models predicting cell cycle phase and age were trained using as an input image stacks for individual cells extracted from 4i experiments (48 fluorescent channels and 4 channels of masks representing entire cell, cell nucleus, cytoplasm and cytoplasmic ring around the nucleus respectively; 52 frames total; 100 px x 100 px) and ground truth annotations obtained from the time lapse imaging. The Fastai (https://github.com/fastai/fastai) Python deep learning library was used for training and the initial pre-trained ResNet-50 convolutional networks were obtained directly from it. Both of the models were trained first using a low-resolution stack (50 px x 50 px) and then fine-tuned using full resolution stacks. The model predicting cell cycle phase was based on 2930 cells divided into a training set (2491 cells; 85%) and a validation set (439 cells, 15%). ResNet-50 CNN predicting phase was trained using a cross-entropy loss function. The model predicting cell cycle age was based on 2767 cells divided into a training set (2352 cells; 85%) and a validation set (415 cells, 15%). ResNet-50 CNN predicting age was trained using mean squared error loss function. Models were trained using Google Cloud VM (8 vCPUs, 52 GB memory, 1 x NVIDIA Tesla P100).

### Manifold learning

Manifold learning was performed using Potential of Heat-diffusion for Affinity-based Transition Embedding (PHATE) (*29*) using the optimal feature set described above as input variables. PHATE was run on z-normalized variables with the following parameter sets: k-nearest neighbor (knn)=200, t=12, gamma=1 for structures presented in **Figs. 1-4**, and knn=75, t=10, gamma=0.25 for structures presented in **Figs. 5-6**.

### Data Visualization

Data were visualized using custom Python scripts (v3.7.1) in Jupyter Notebooks (v6.1.4), GraphPad Prism (v8) and scanpy (v1.6) (*53*). Scanpy was also used to prepare data for visualization in virtual reality using the singlecellVR website (singlecellvr.com) (*54*).

### siRNA

RPE cells were treated with DMSO or etoposide (1 μM) for 24h then transfected with non-targeting (Dharmacon, D-001810-10-0) or cyclin D1/D2/D3 (SMARTPools L-003210-00-0005/L-003211-00-0005/L-003212-00-0005) siRNA pools using the DharmaFECT 1 transfection reagent (T-2001-01) as per manufacturer’s protocol and incubated for 3 days prior to fixation. Immunofluorescence was performed as per the 4i protocol described above.

---

**Figure S1.**
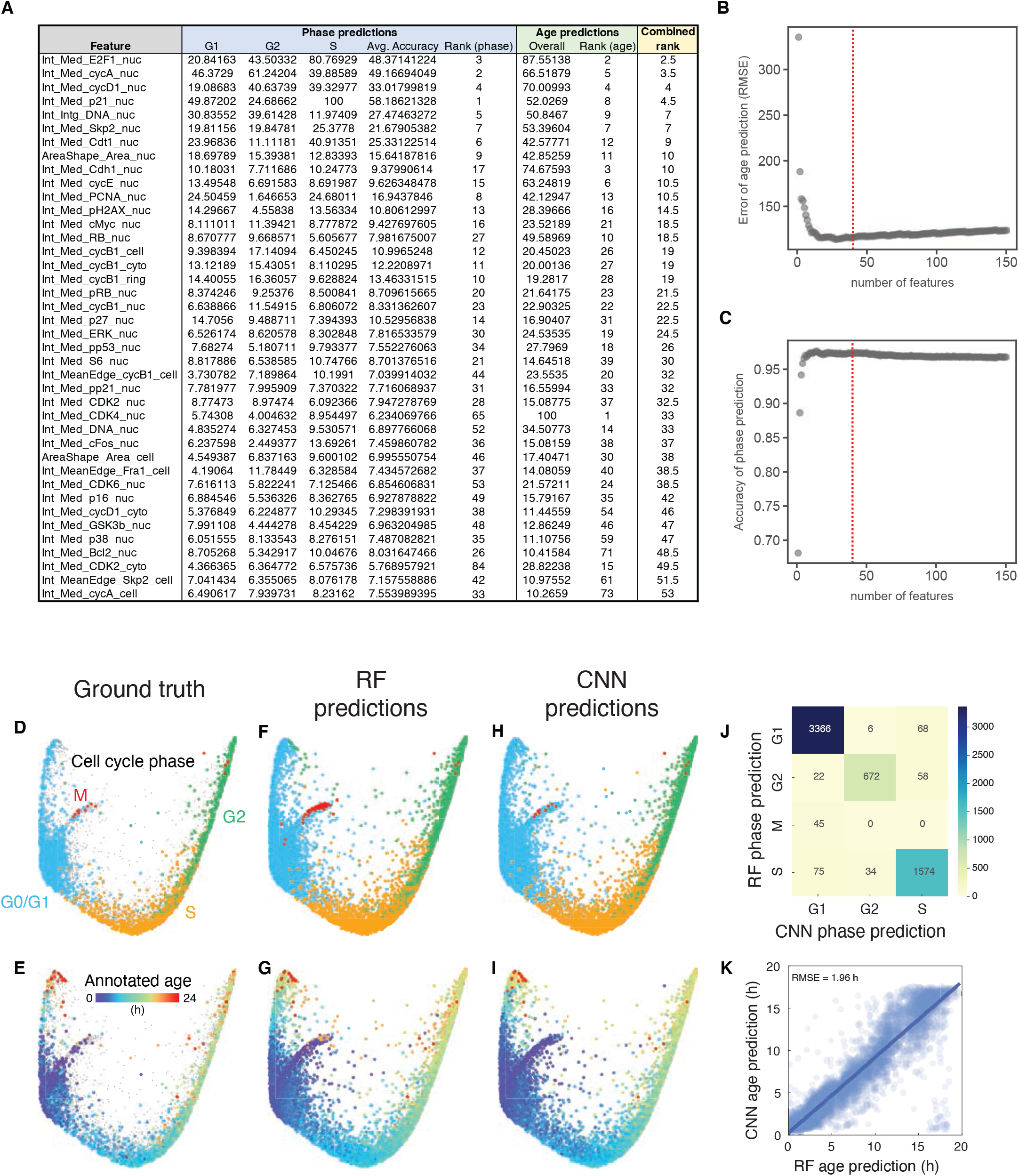
- Generation and annotation of the human cell cycle structure. (**A**) Random forest (RF) models were used to predict cell cycle age and phase annotations (obtained by time-lapse imaging) from the multivariate 4i signatures (see Methods). Variable importance tables were obtained that rank each feature by its importance in maximizing accuracy of phase predictions or reducing the error of age predictions (see full tables in Table 2). These rankings were averaged to obtain a combined rank for each feature. (**B-C**) Iterative random forest models were used to calculate prediction error for cell cycle age (B, RMSE: root mean squared error) and accuracy for phase (C) while increasing the number of features that were for training. Features were successively added in each iterative round, ordered by their combined rank as determined in A. Shown are iterations for the first 150 ranked features. The top 40 features were chosen as an optimal feature set and used as inputs for PHATE to generate the cell cycle structure. (**D-E**) Ground truth annotations of cell cycle phase (D) and age (E). (**F-G**) Random forest predictions of phase (F) and age (G) (same figures are presented in Fig. 1C-D, and are shown here for comparison). (**H-I**) Convolutional neural network (CNN) predictions of phase (F) and age (G). For panels F-I, phase and age ground truth annotations are also plotted on the structures. (**J**) Confusion matrix comparing RF and CNN phase predictions. The CNN model did not predict any M phase cells. (**K**) Concordance between RF and CNN predictions for age. Int: intensity; Med: median; nuc: nuclear; cyto: cytoplasm; ring: perinuclear area; MeanEdge: plasma membrane

**Figure S2.**
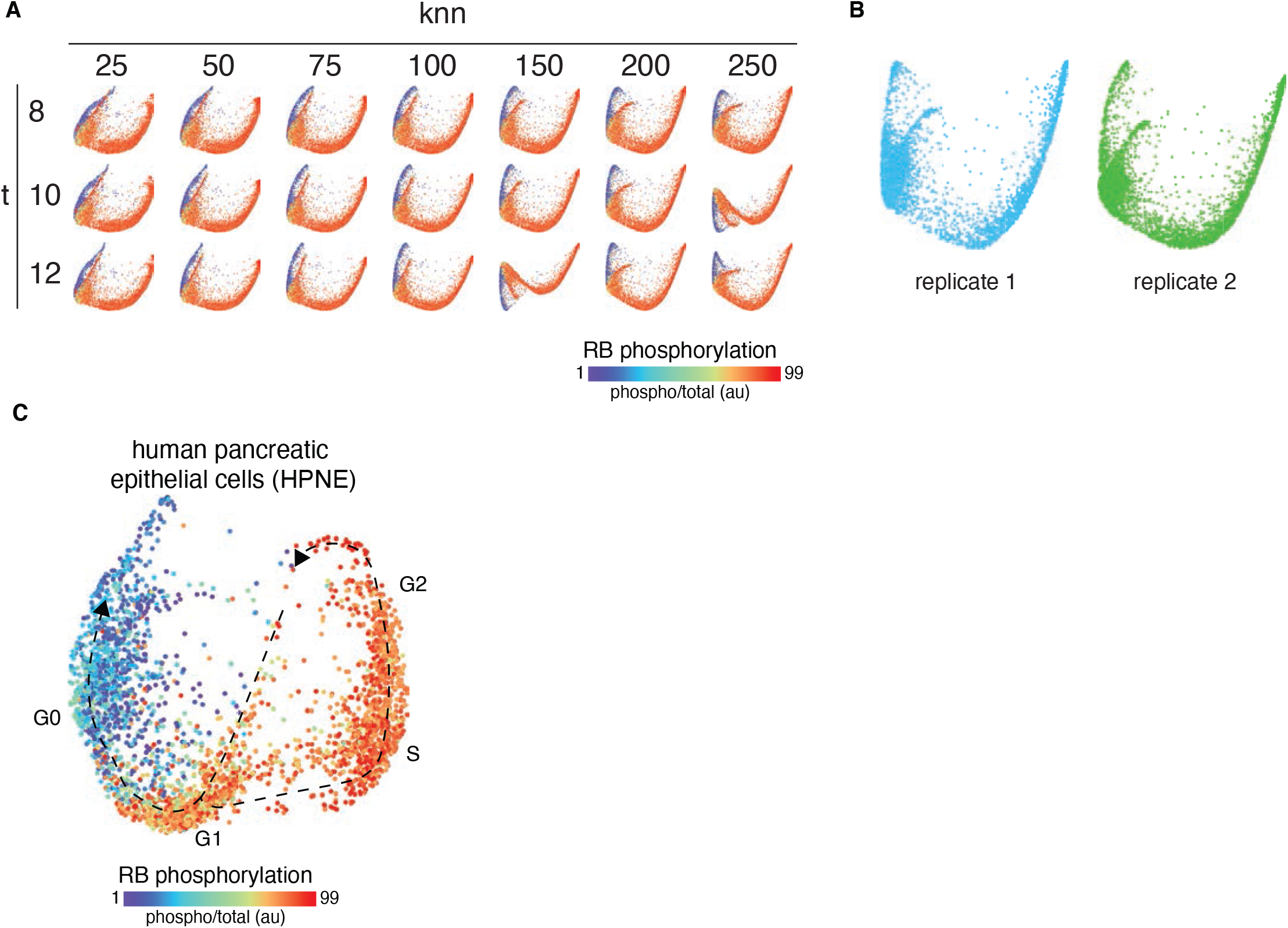
- Conservation of cell cycle structure. **(A)** Cell cycle structures generated from a PHATE parameter screen across a range of k-nearest neighbor (knn) and power (t) values. **(B)** Cell cycle structures generated from two replicate populations of RPE cells using the same feature sets and PHATE parameters. **(C)** Cell cycle structure obtained from the single-cell, multivariate cell cycle signatures obtained by 4i of human pancreatic epithelial cells (HPNE).

**Figure S3.**
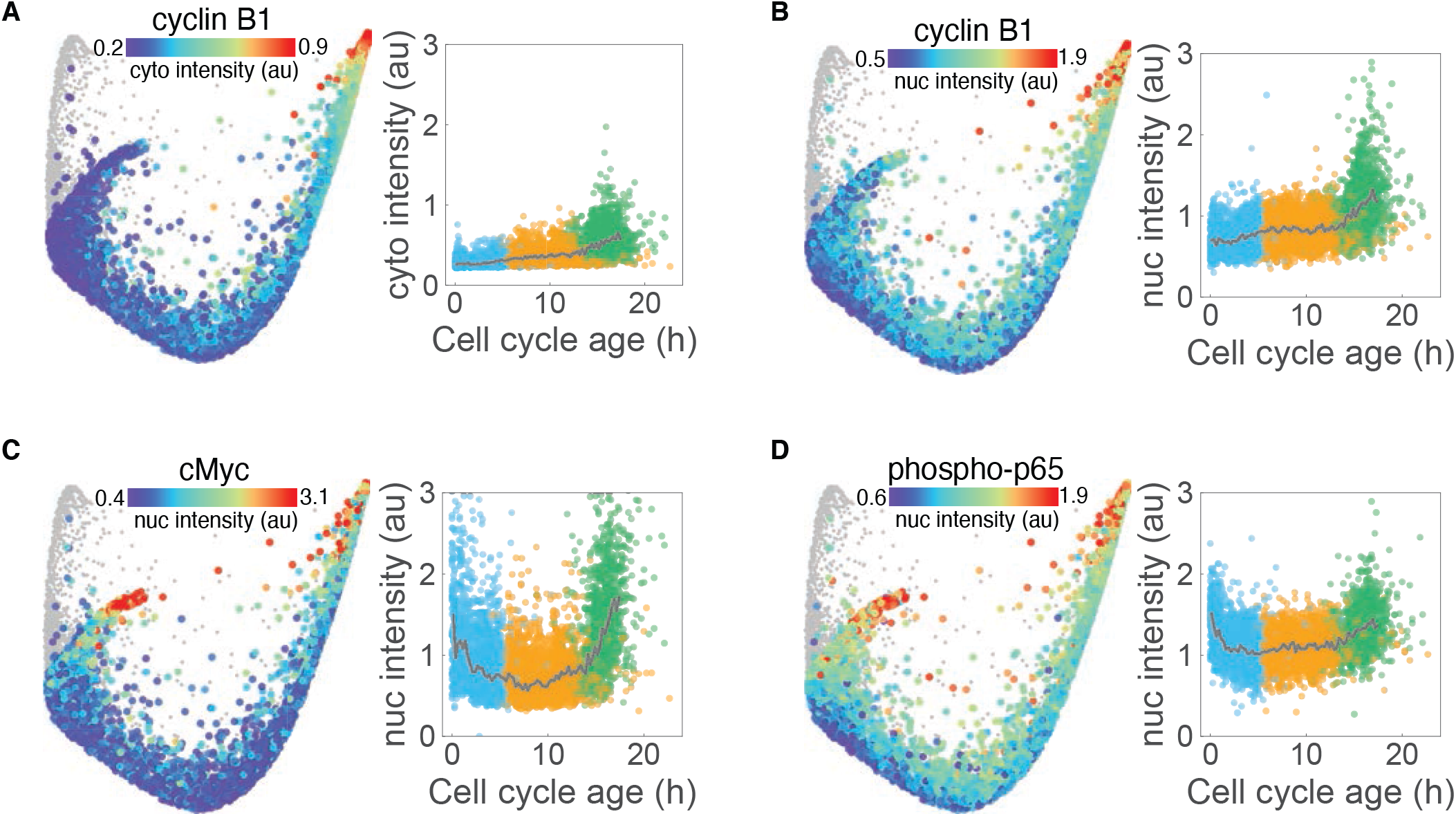
- Effector dynamics along the proliferative trajectory. Median cytoplasmic expression of cyclin B1 (A) and median nuclear expression of cyclin B1 (B), cMyc (C) and phospho-p65 (D) are mapped onto the proliferative trajectory of the cell cycle structure (left panels) and plotted against cell cycle age (right panels). Population medians in time courses indicated by solid grey lines and individual cells are colored by cell cycle phase (G1: blue, S: orange, G2: green). Non-cycling (G0) cells (phospho/total RB < 1.6) are shown in grey on the structure and are excluded from time courses.

**Figure S4.**
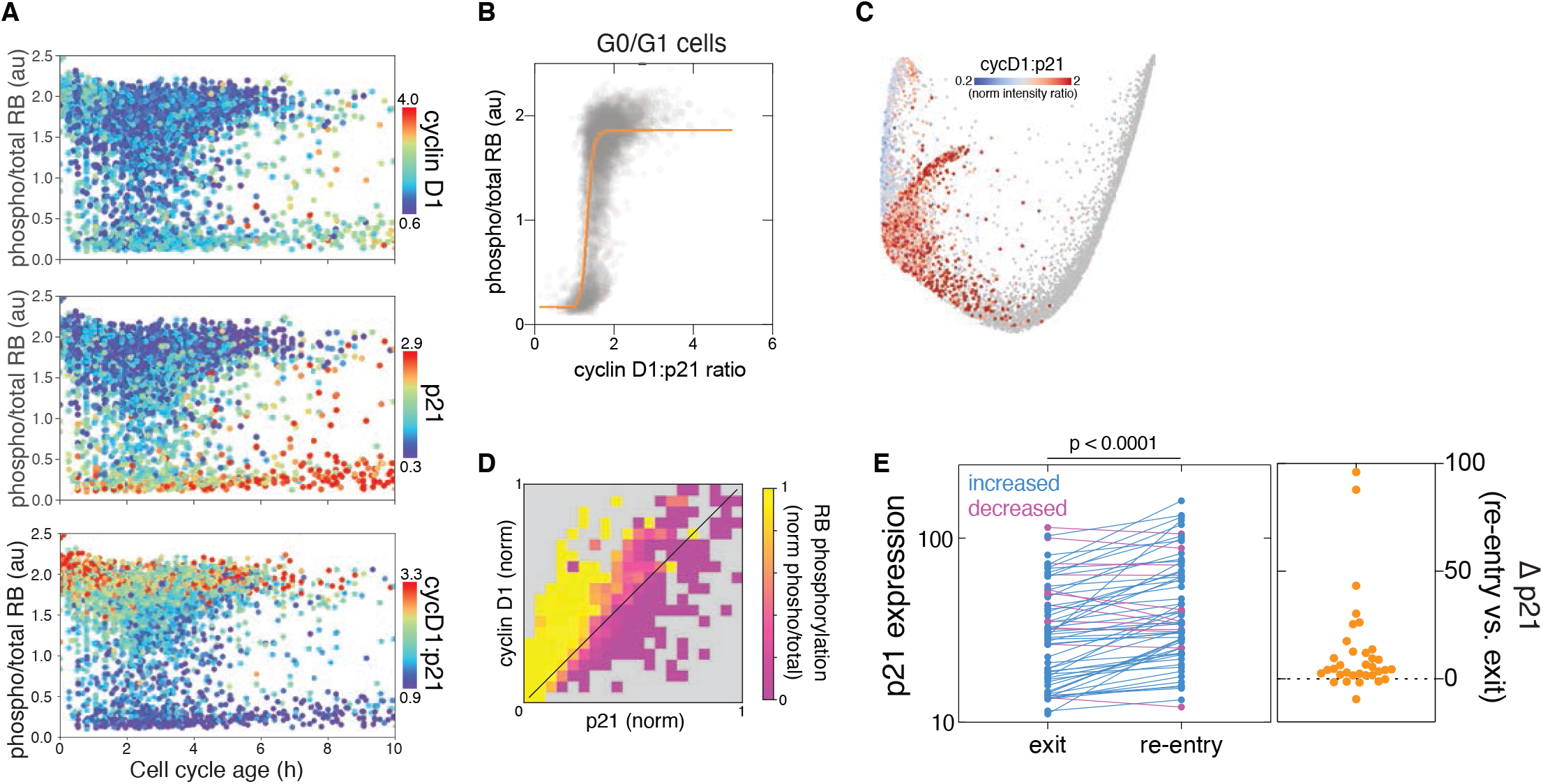
- The cyclin D1:p21 ratio regulates both cell cycle exit and re-entry. (**A**) Time courses of RB phosphorylation following mitosis are colored by median nuclear expression of cyclin D1 (upper panel) and p21 (middle panel) and the cyclin D1:p21 ratio (lower panel). Note the delay in RB dephosphorylation immediately following mitosis (<1h). (**B**) RB phosphorylation after cell division is controlled by the cyclin D1:p21 ratio in individual cells in an ultrasensitive manner (Hill coefficient = 12.2). Data from G0/G1 cells are shown. (**C**) The cyclin D1:p21 ratio of G0/G1 cells is mapped onto the cell cycle structure. Data is expressed as a z-normalized nuclear intensity ratio. (**D**) Heatmap showing the proportion of cells with high RB phosphorylation (phospho/total RB > 1.6) at a given ratio of z-normalized cyclin D1:p21 expression. (**E**) *Left panel*: Single cell p21 expression measured by time lapse imaging at the time of cell cycle exit (1h post-mitosis) versus re-entry (at time of CDK2 reactivation). *Right panel:* Difference in p21 expression at cell cycle re-entry versus exit. Statistical significance determined using a Student’s paired *t* test. *N* = 65 cells.

**Figure S5.**
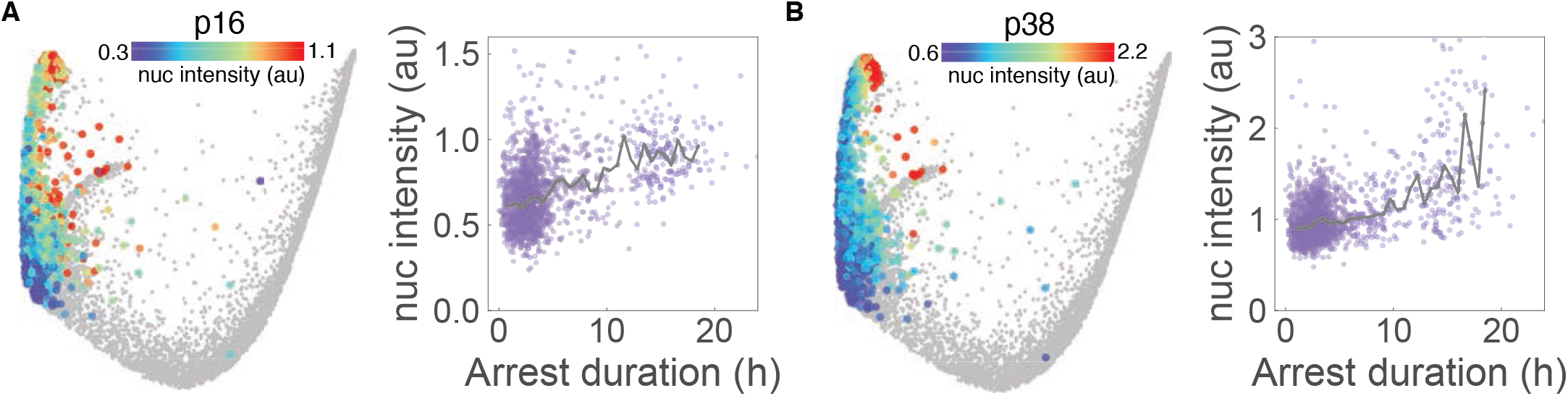
- Effector dynamics along the arrest trajectory. (**A-B**) Median nuclear expression of p16 (A) and p38 (B) are mapped onto the arrest trajectory of the cell cycle structure (left panels) and plotted against time of arrest (right panels). Population medians in time courses indicated by solid grey lines. Non-arrested cells (phospho/total RB > 1.6) are shown in grey on the structure and are excluded from time courses.

**Figure S6.**
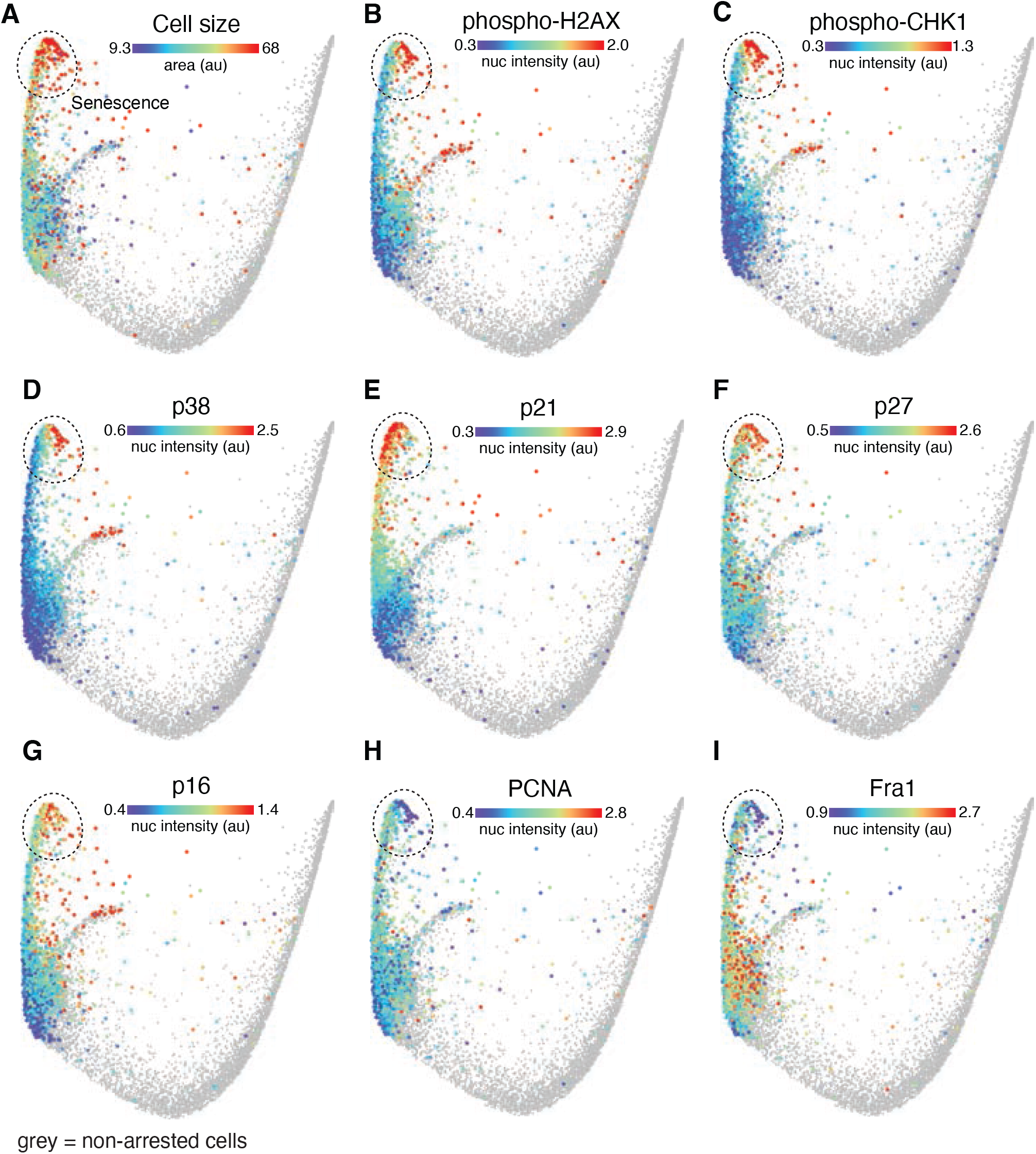
- Molecular signature of cellular senescence. **(A-I)** Cell size (A) and median nuclear intensities of phospho-H2AX (B), phospho-CHK1 (C), p38 (D), p21 (E), p27 (F), p16 (G), PCNA (H) and Fra1 (I) of arrested cells are mapped onto the cell cycle structure. Non-arrested cells (phospho/total RB > 1.6) are shown in grey.

**Figure S7.**
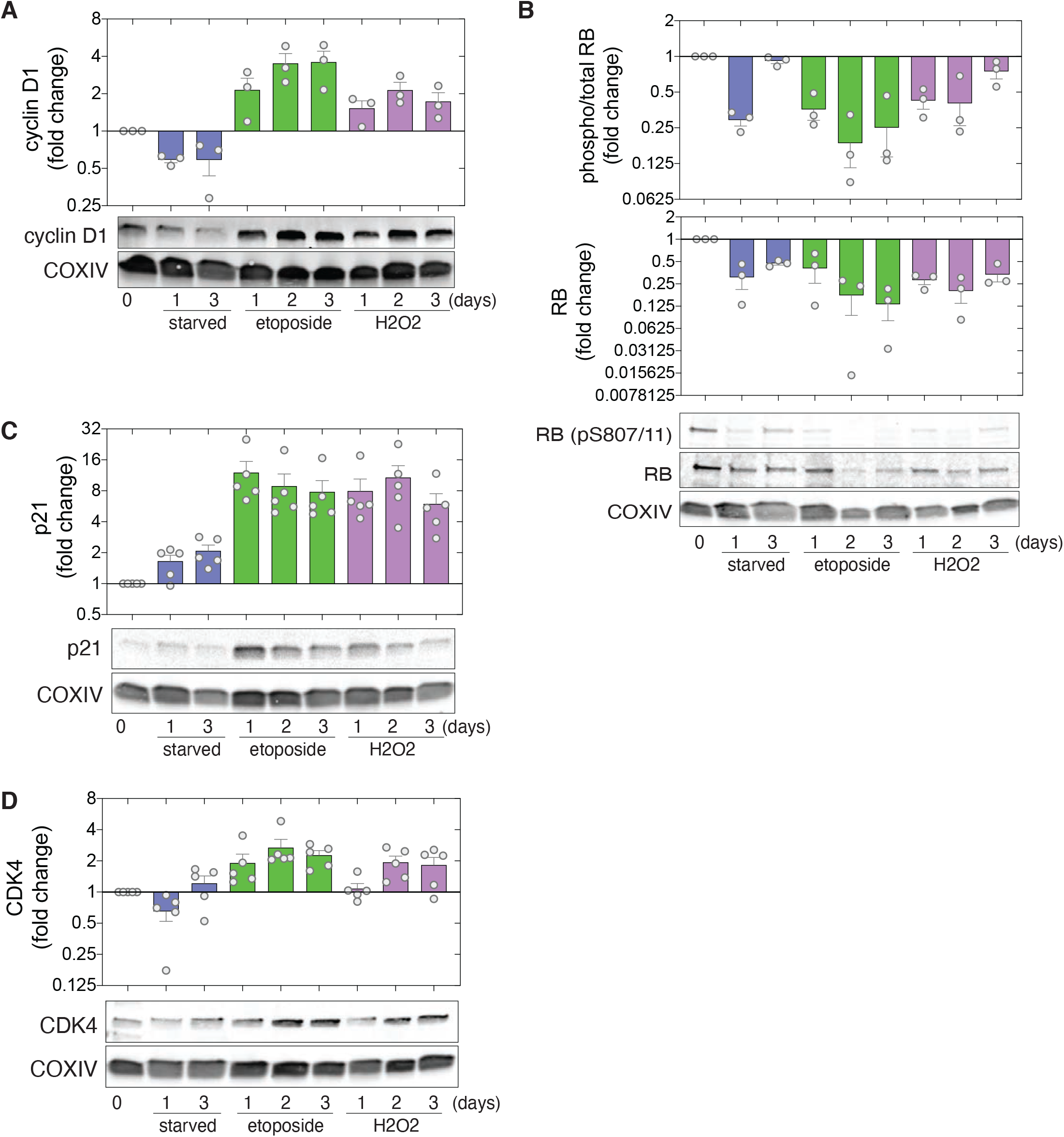
Population arrest signatures in response to cell cycle stresses. (**A-D**) Representative western blots (lower panels) and quantification (upper panels) of cyclin D1 (A), phospho/total and total RB (B), p21 (C) and CDK4 (D) in unperturbed, serum-starved (1-3 d), etoposide-treated (1 μM, 1-3 d) or H2O2-treated (200 μM, 1-3 d) cell populations. Data were normalized by COXIV expression (as a loading control). Data represent means +/- sem from at least 3 independent experiments. Experimental replicates are shown as filled circles.

**Figure S8.**
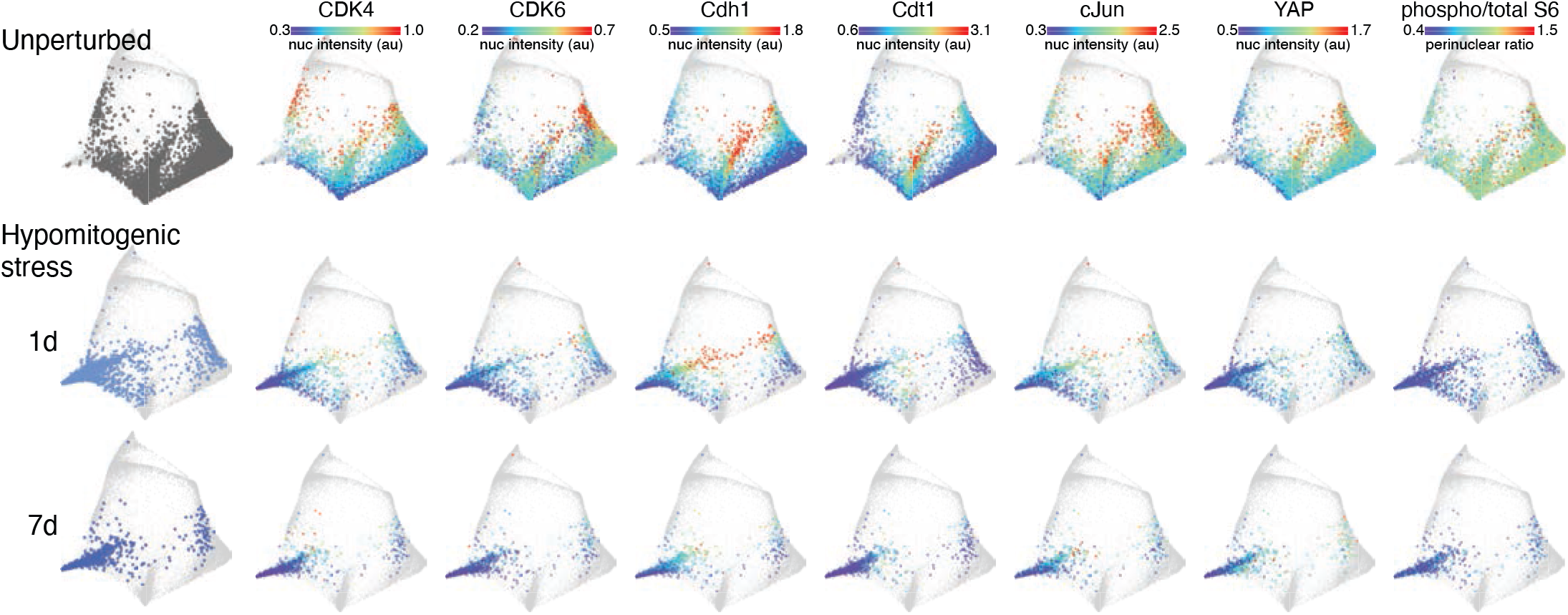
- Cell cycle signature of hypomitogenic stress. Median nuclear intensities of CDK4, CDK6, Cdh1, Cdt1, cJun and YAP, and perinucear S6 activation (phospho/total) are mapped onto unperturbed or serum-starved cells (“hypomitogenic stress”, 1 or 7 days).

**Figure S9.**
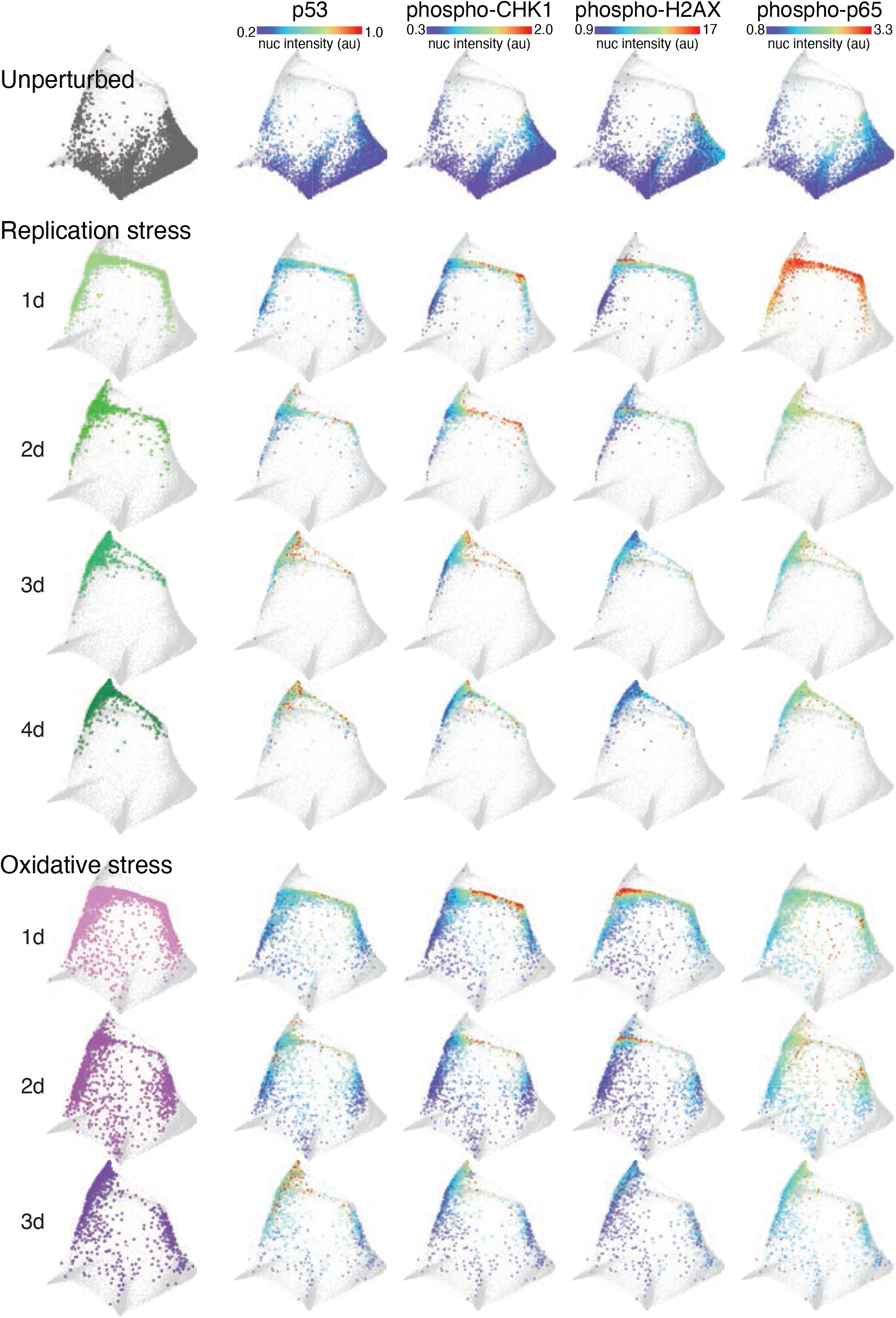
- Cell cycle signatures of replication and oxidative stress. Median nuclear intensities of p53, phospho-CHK1, phospho-H2AX and phospho-p65 are mapped onto unperturbed (grey), etoposide-treated (“replication stress”, 1 μM, 1-4 d) and H2O2-treated (“oxidative stress”, 200 μM, 1-3 d) cells.

**Figure S10.**
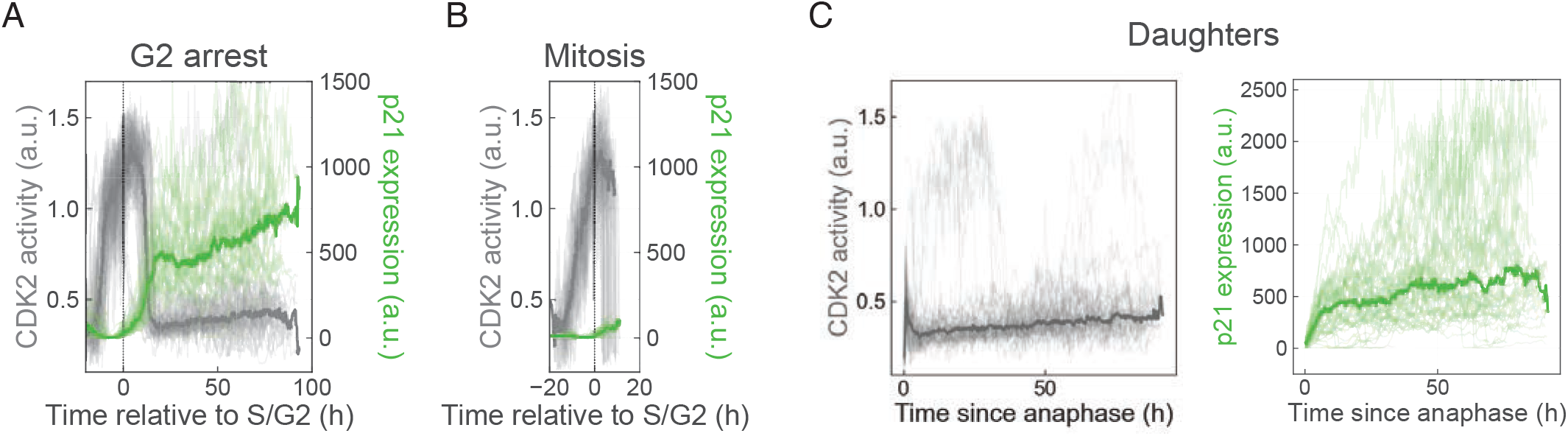
Time-lapse imaging of cell fate following etoposide treatment. (**A-B**) Single cell traces from time-lapse imaging of CDK2 activity (blue, quantified by the cytoplasm:nuclear ratio of DHB-mCherry) and p21-YPet expression (green) in cells undergoing G2 arrest (A) or mitosis (B) following etoposide treatment (1 μM) aligned at the S/G2 transition. (**C**) Single cell traces of the daughters of mitotic cells (from panel B). Thick lines represent population medians. These data are shown as heatmaps in Figure 5D-F. *N* = 117 cells

**Figure S11.**
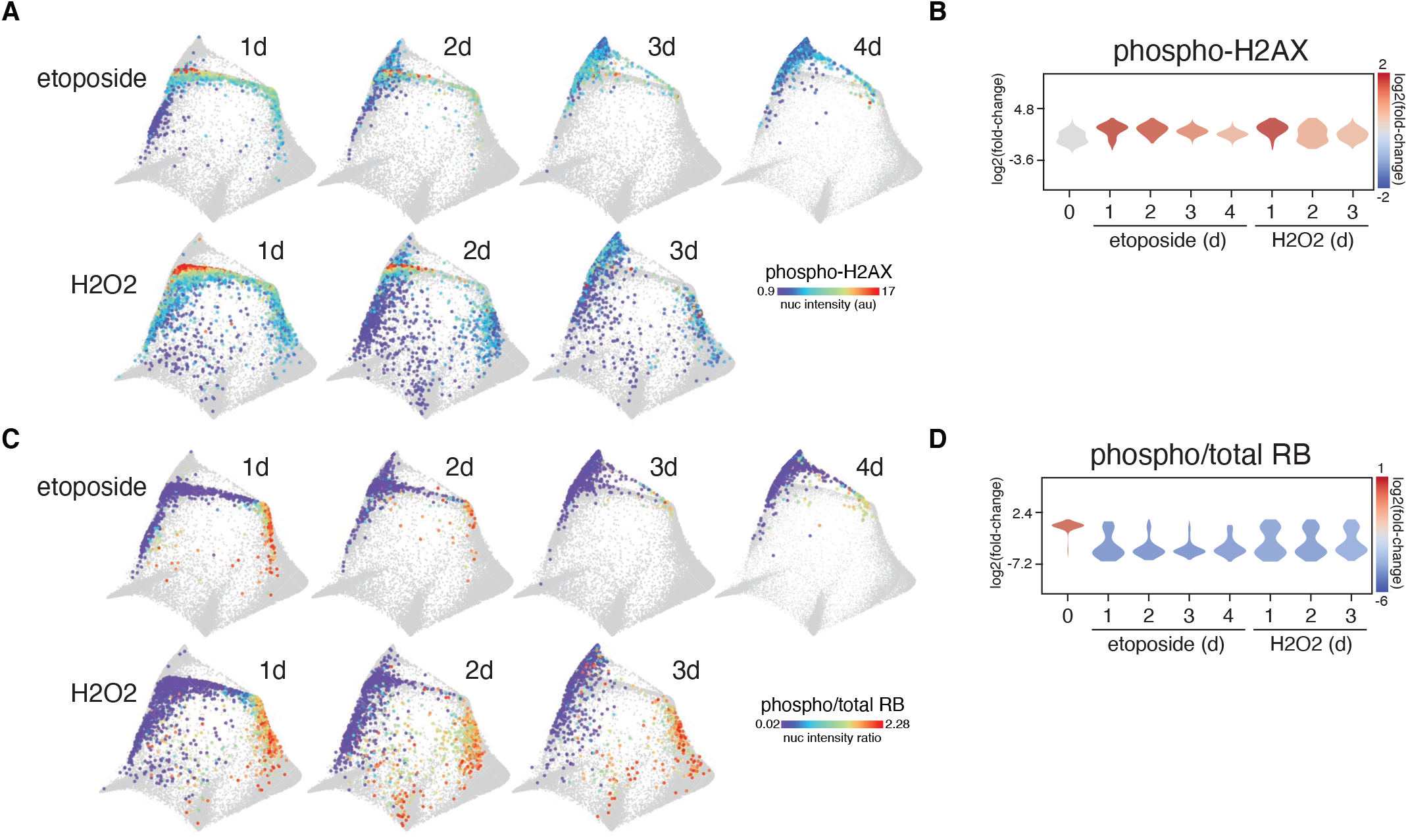
- Etoposide and H2O2 induce sustained and transient DNA damage responses, respectively. (**A-B**) Median nuclear phospho-H2AX intensity is mapped onto etoposide- and H2O2-treated cells (A) and plotted as single cell distributions (B). (**C-D**) RB phosphorylation (phospho/total) is mapped onto etoposide- and H2O2-treated cells (C) and plotted as single cell distributions (D).

**Figure S12.**
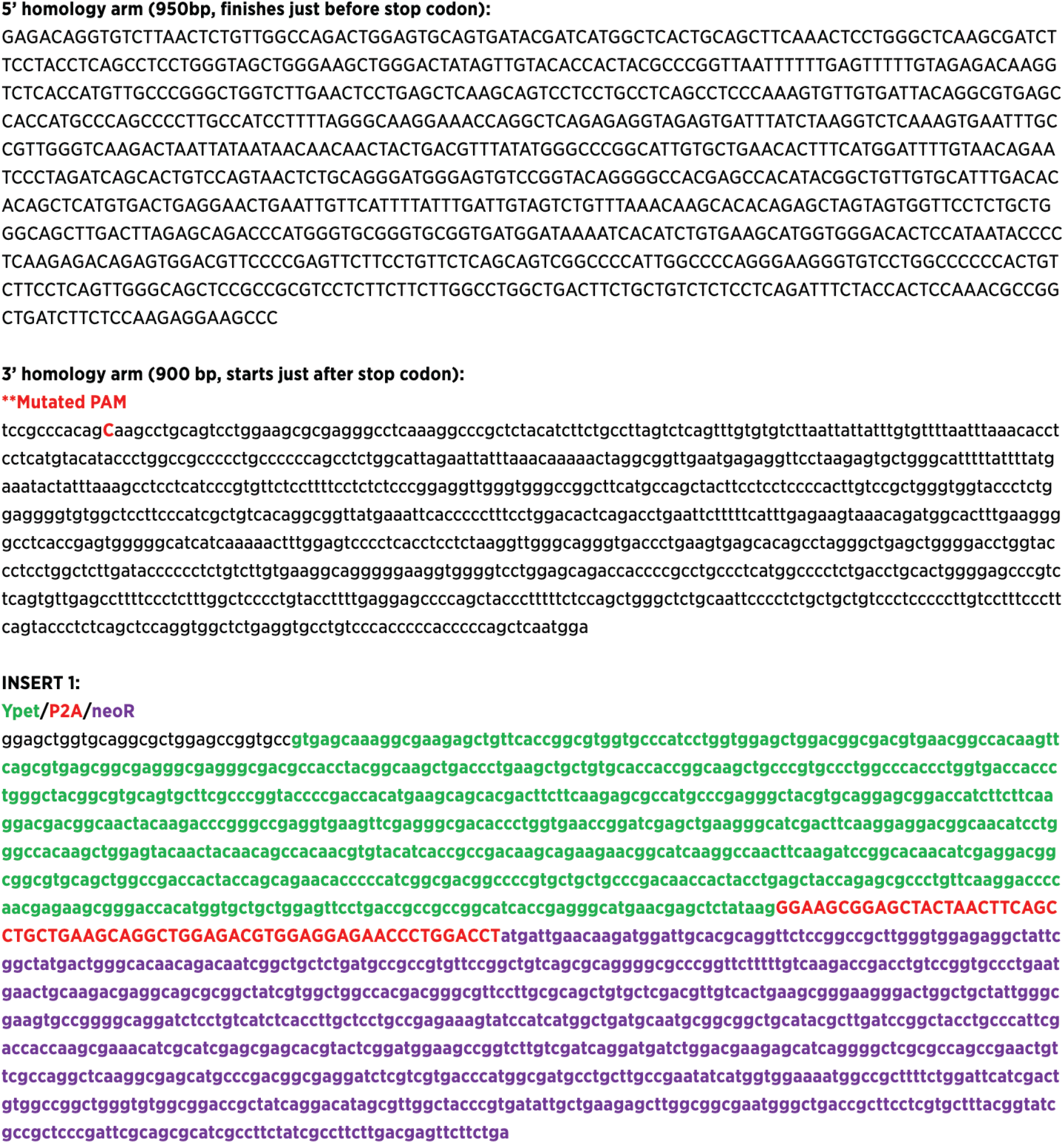
Donor cassette used to generate p21-YPet gene by CRISPR-Cas9-mediated knock-in.

**Table S1.**
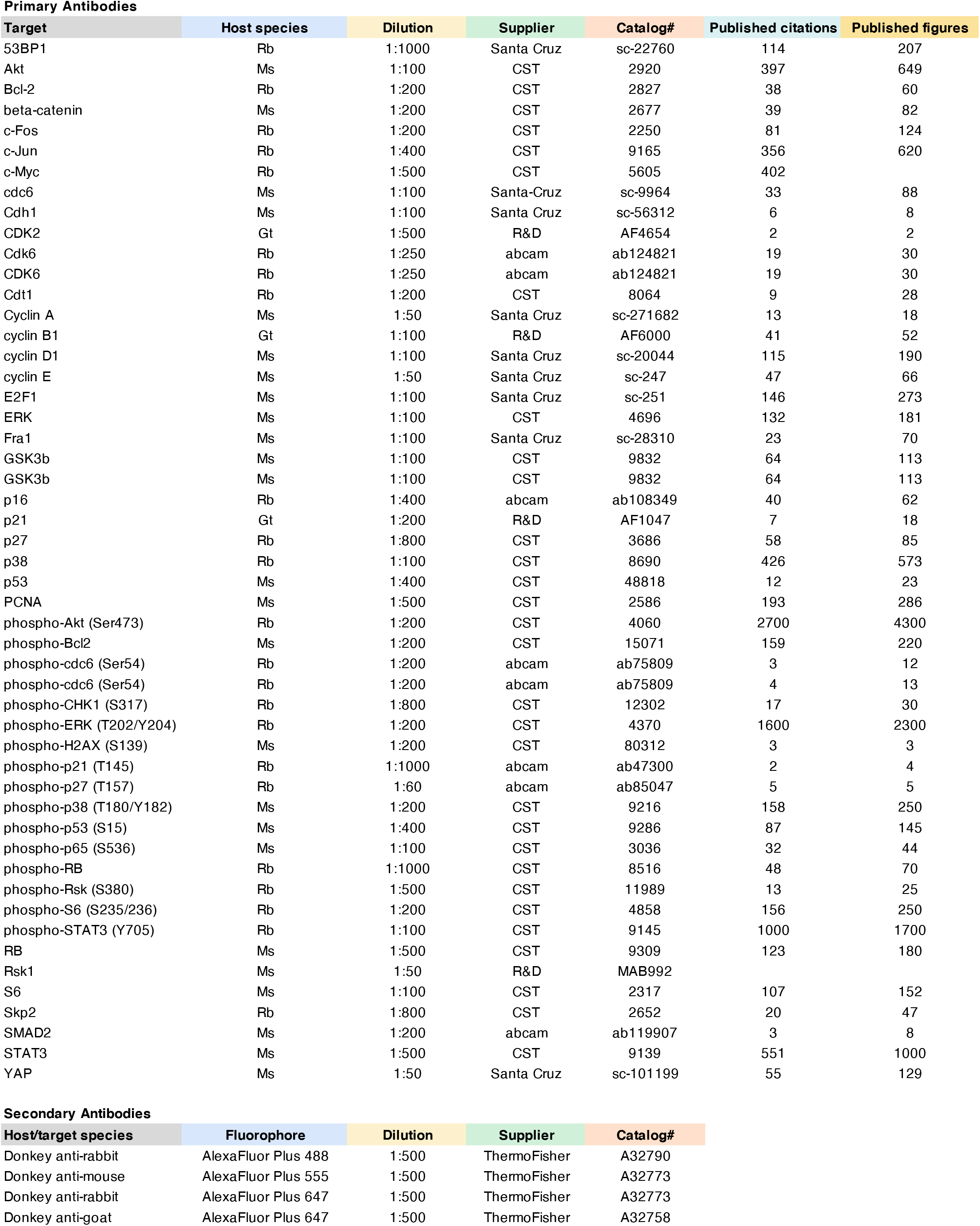
Antibodies.

**Table S2.**
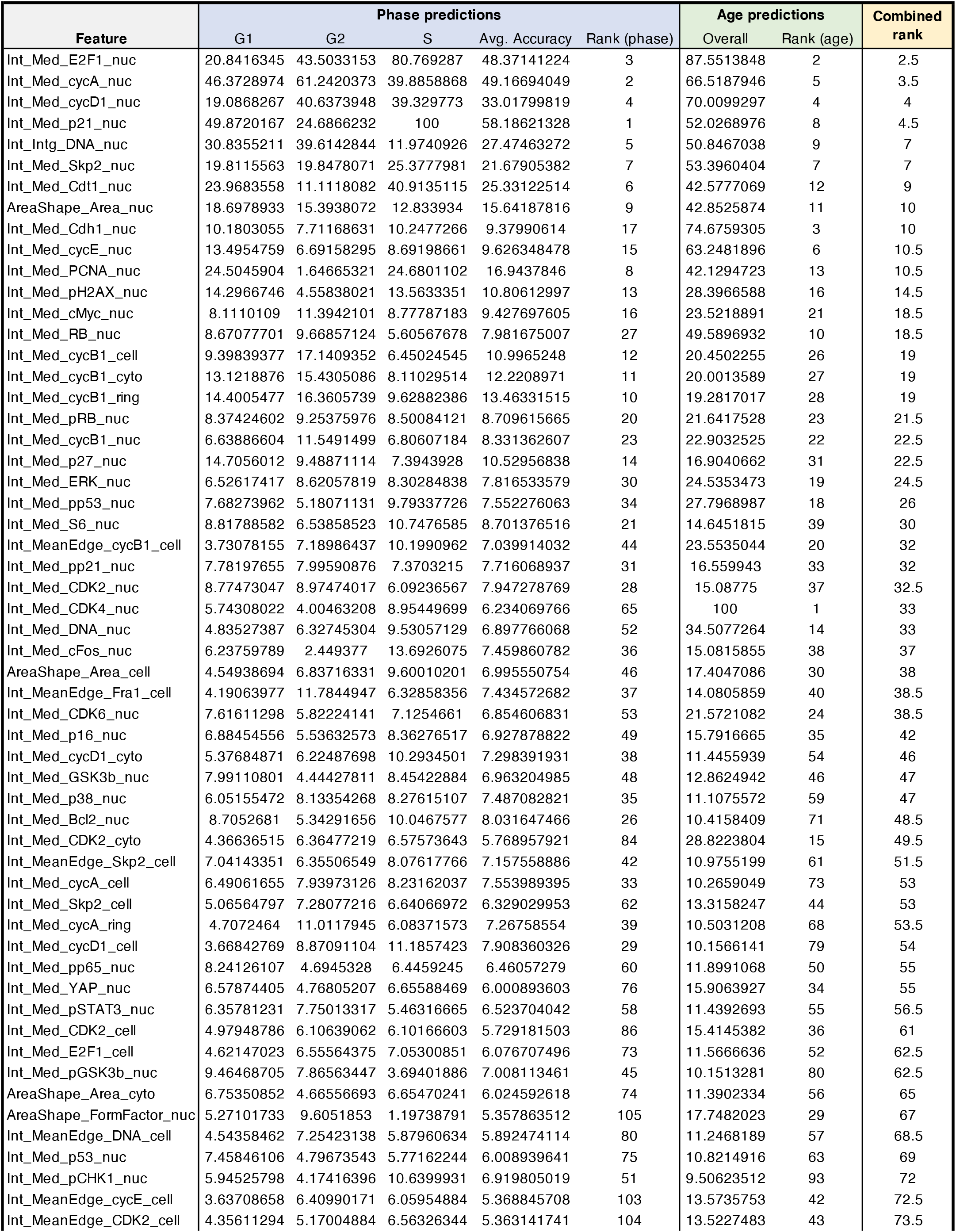
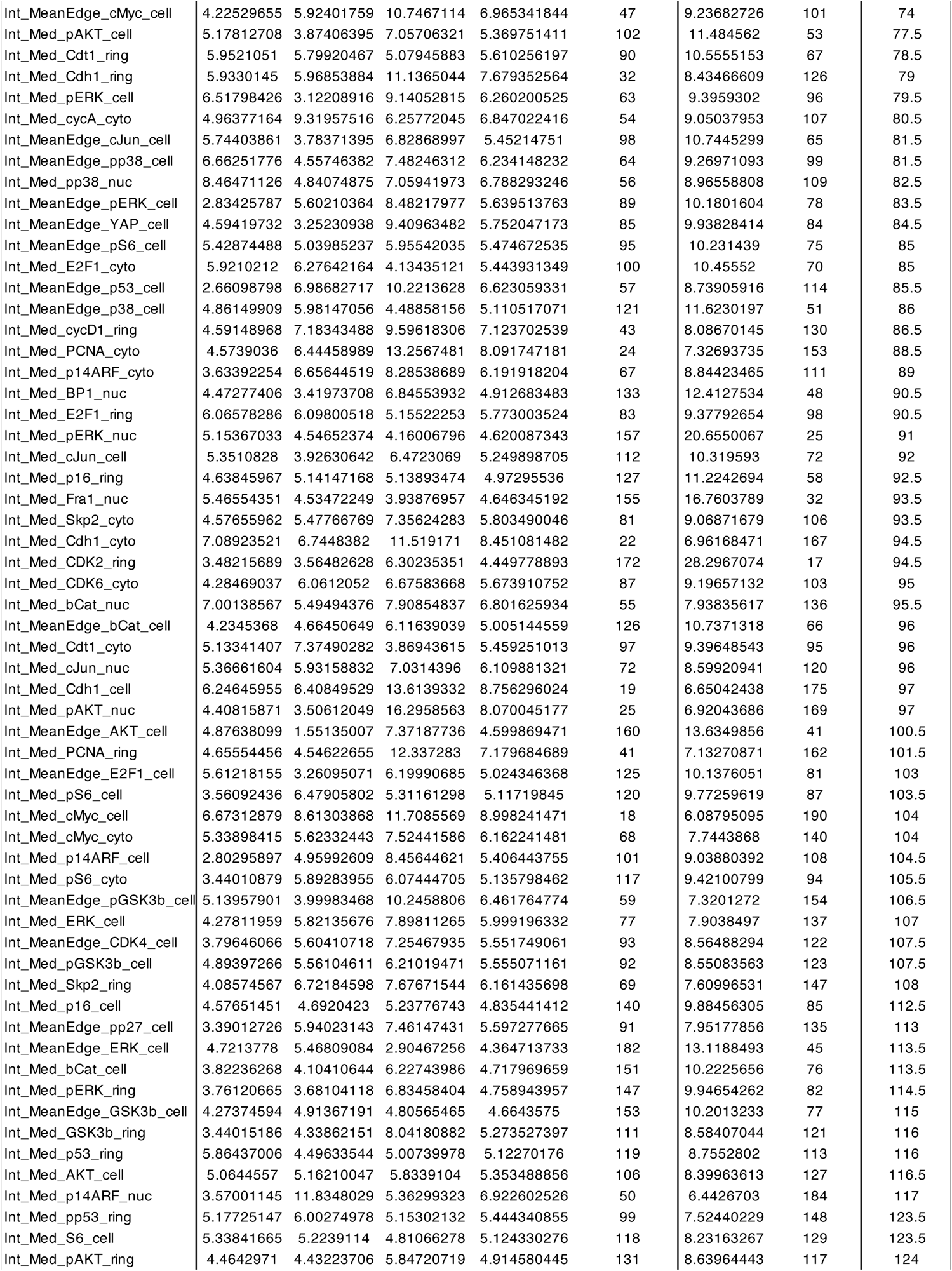
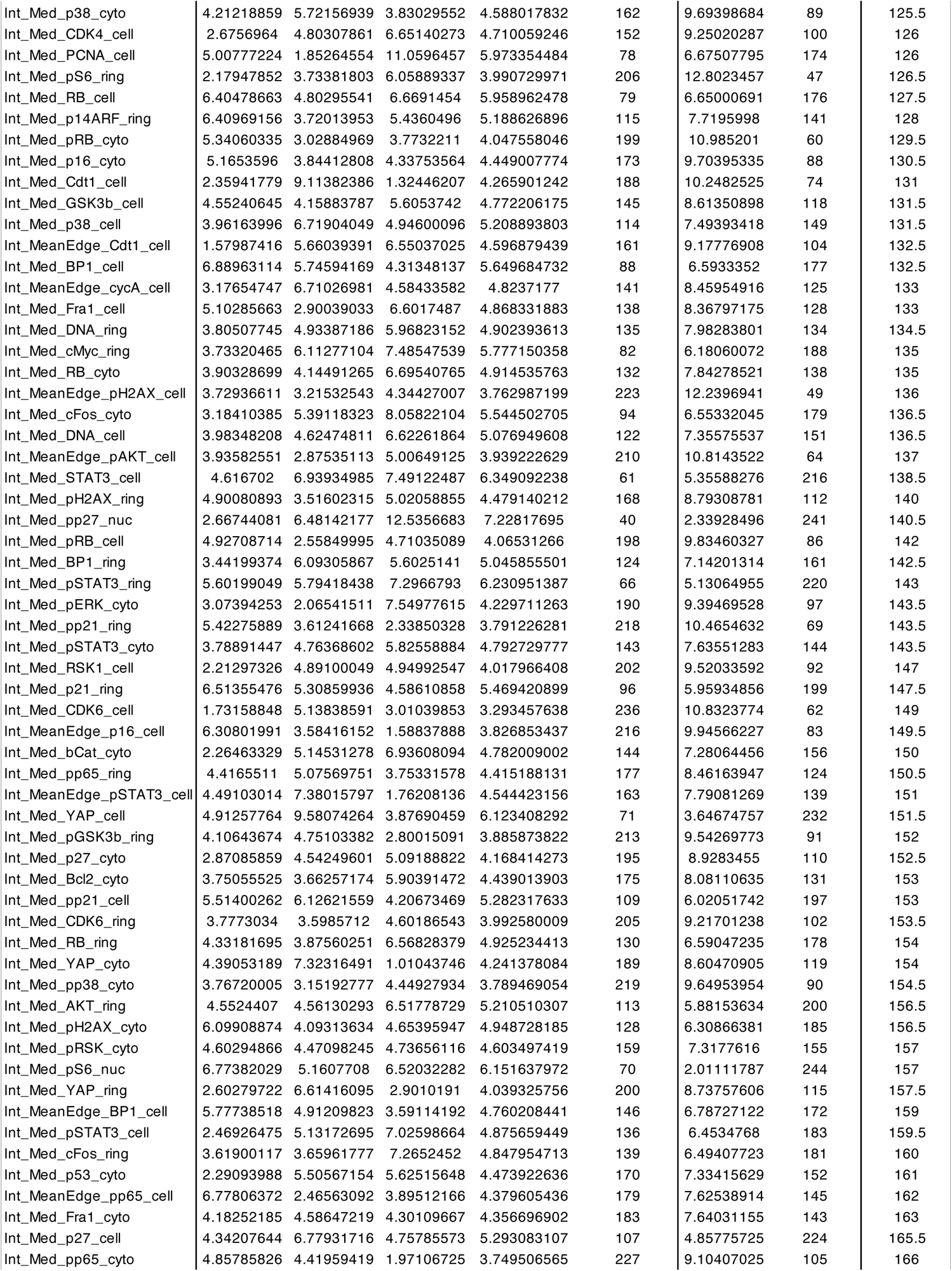
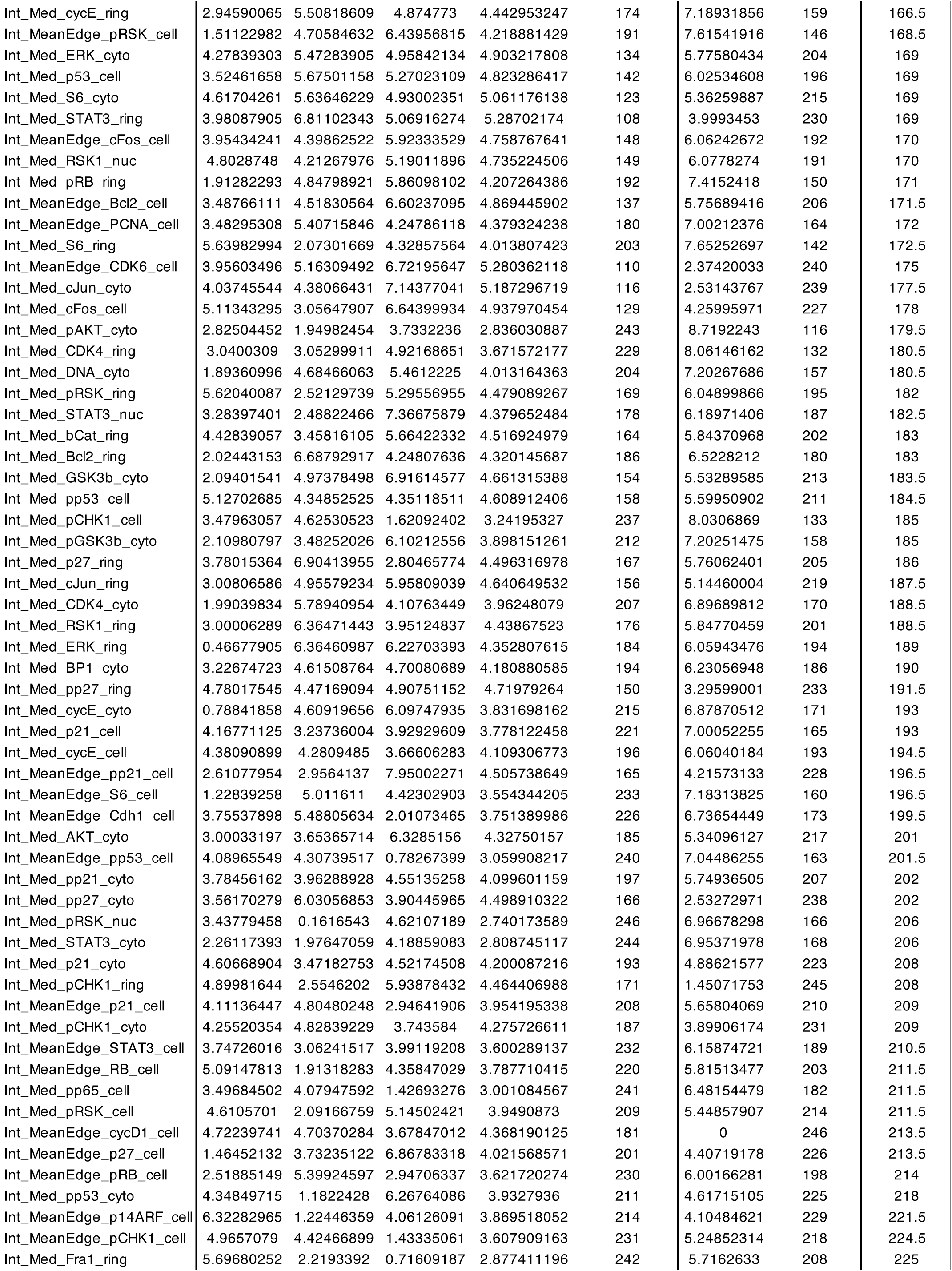
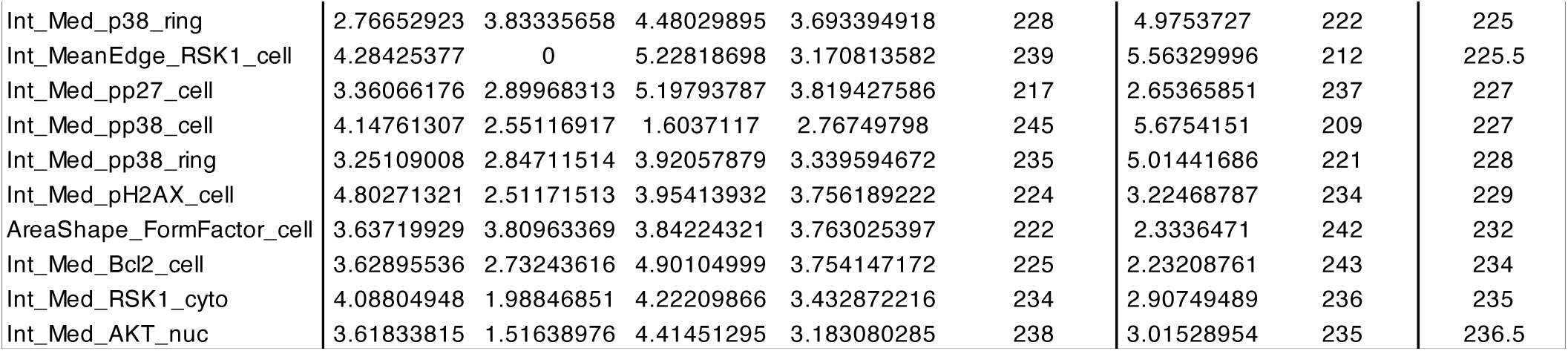
Variable importance for cell cycle phase and age determined by random forest modeling.

