## Supplementary material for "The structure of the human cell cycle": Auxiliary fig S1. Effector dynamics along the proliferative trajectory

**Auxiliary Figure S1. Effector dynamics along the proliferative trajectory.** Features are mapped onto the proliferative trajectory of the cell cycle structure (left panels) and plotted against cell cycle age (right panels). Population medians in time courses indicated by solid grey lines and individual cells are colored by cell cycle phase (G1: blue, S: orange, G2: green). Non-cycling (G0) cells (phospho/total RB $< 1.6$) are shown in grey on the structure and are excluded from time courses. Plasma membrane, PM; perinuclear, PN.

**Aux Fig. S1 - Proliferative trajectory - Page 1**

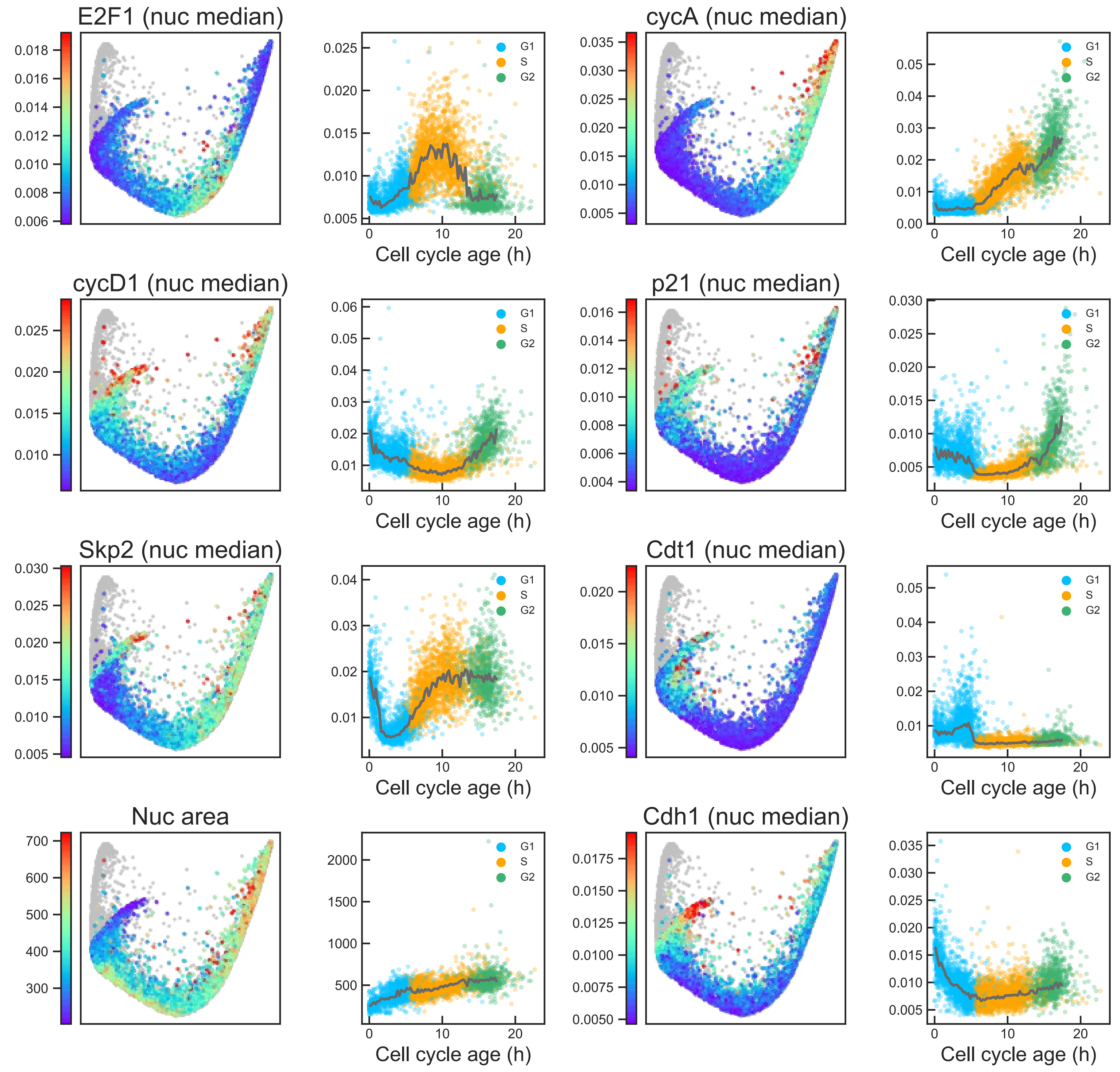

### Aux Fig. S1 - Proliferative trajectory - Page 2

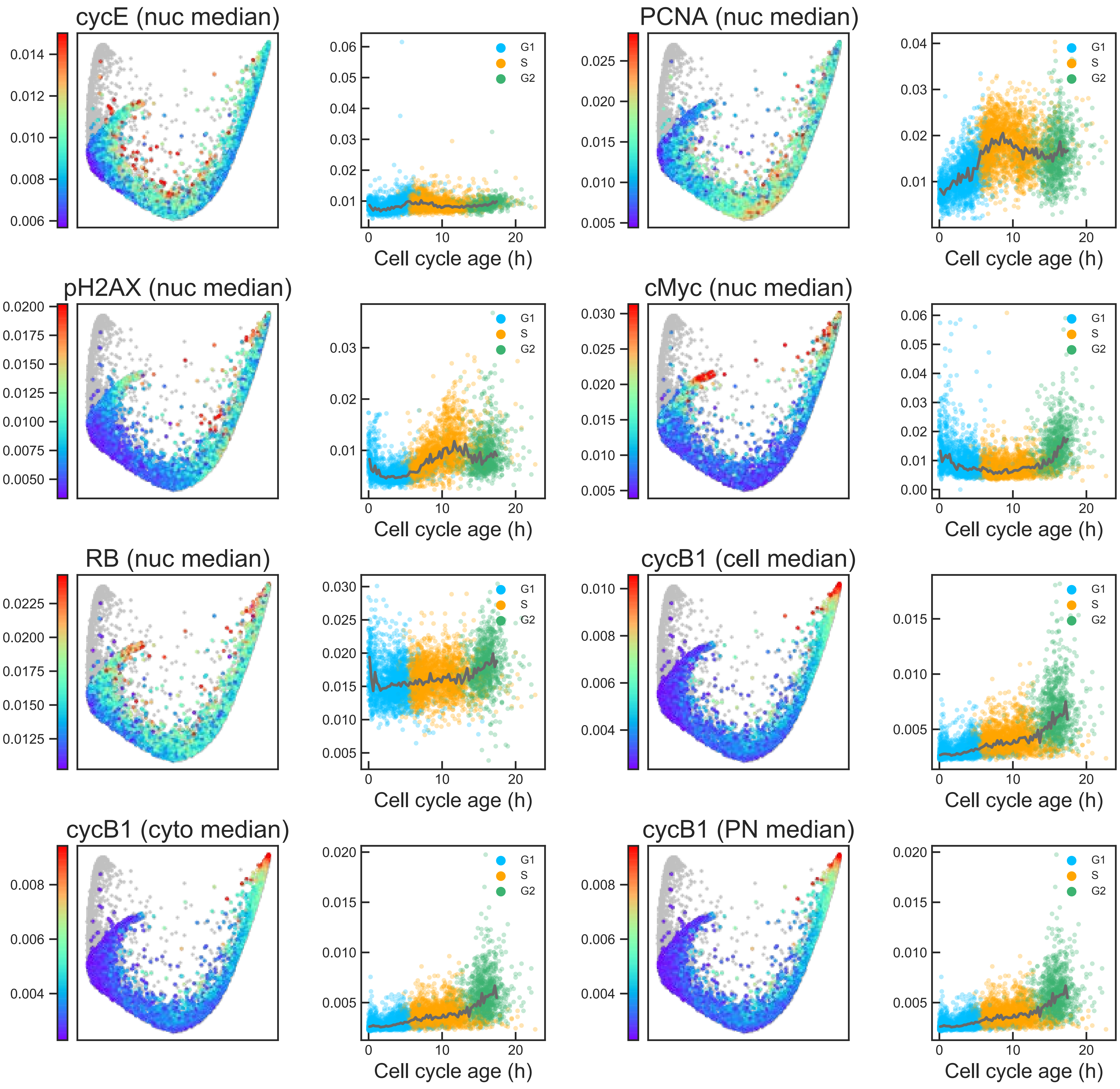

#### Aux Fig. S1 - Proliferative trajectory - Page 3

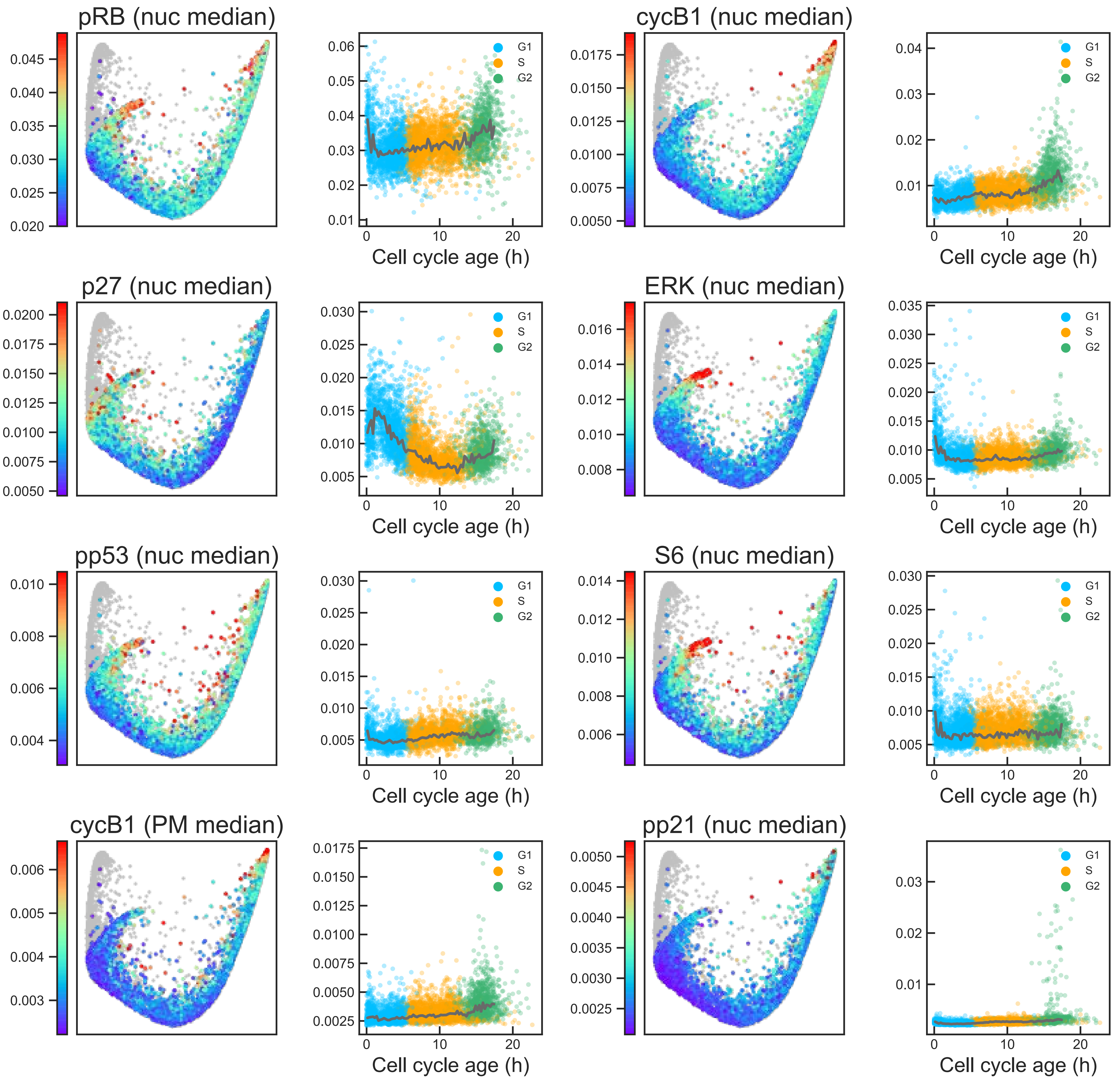

#### Aux Fig. S1 - Proliferative trajectory - Page 4

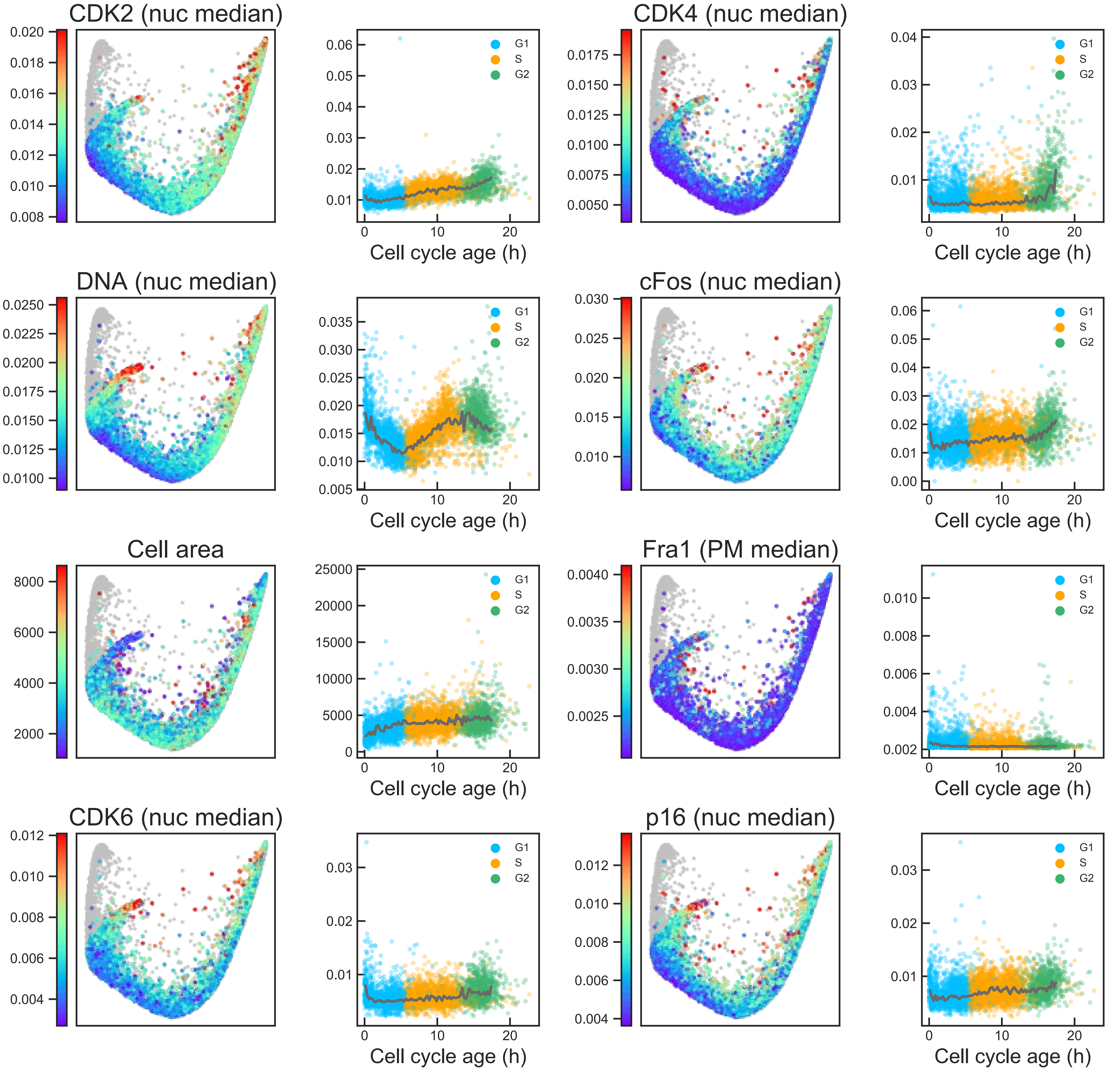

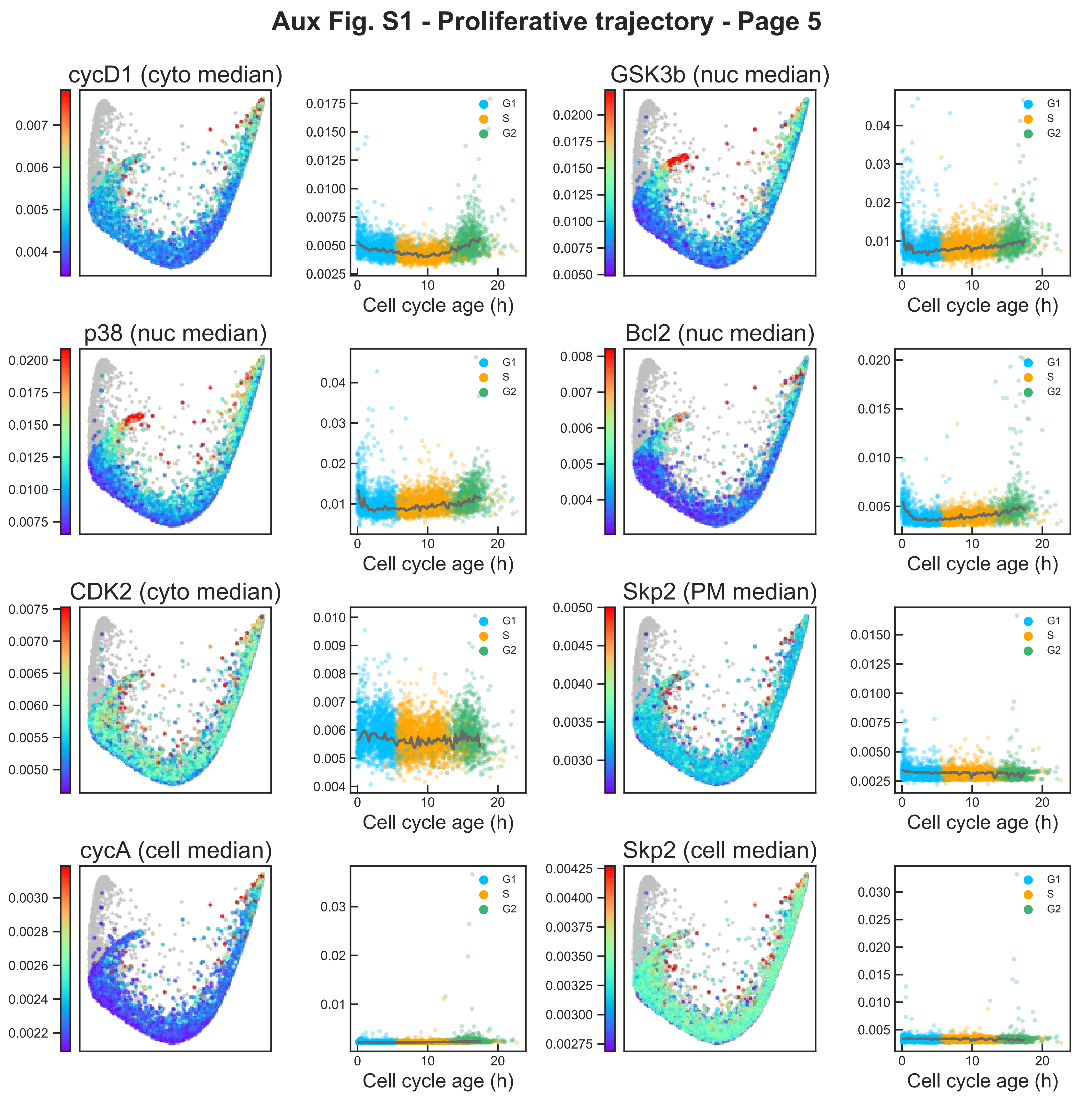
Aux Fig. S1 - Proliferative trajectory - Page 5
cycD1 (cyto median)
GSK3b (nuc median)
p38 (nuc median)
Bcl2 (nuc median)
CDK2 (cyto median)
Skp2 (PM median)
cycA (cell median)
Skp2 (cell median)
G1
S
G2
Cell cycle age (h)

### Aux Fig. S1 - Proliferative trajectory - Page 6

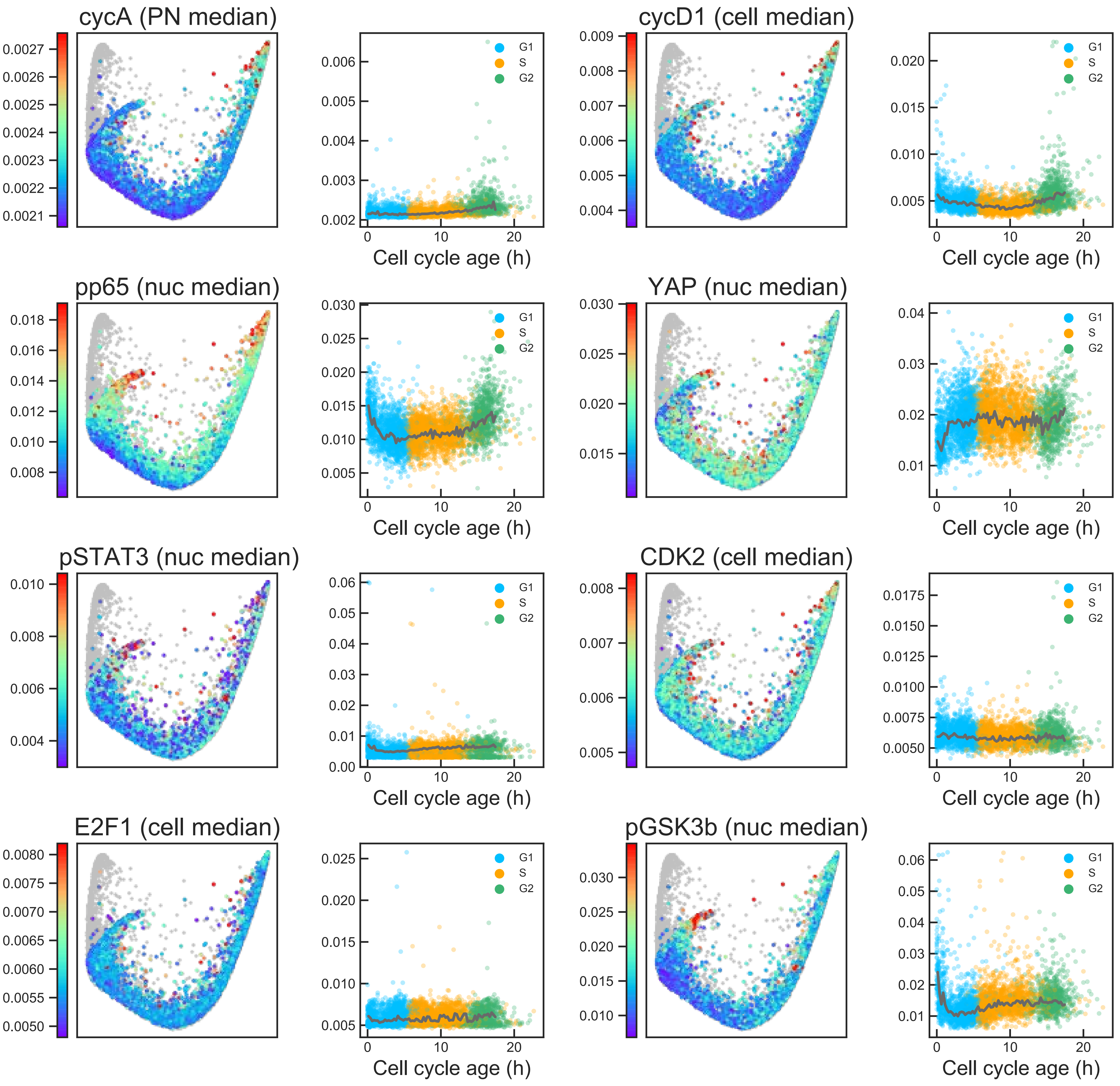

### Aux Fig. S1 - Proliferative trajectory - Page 7

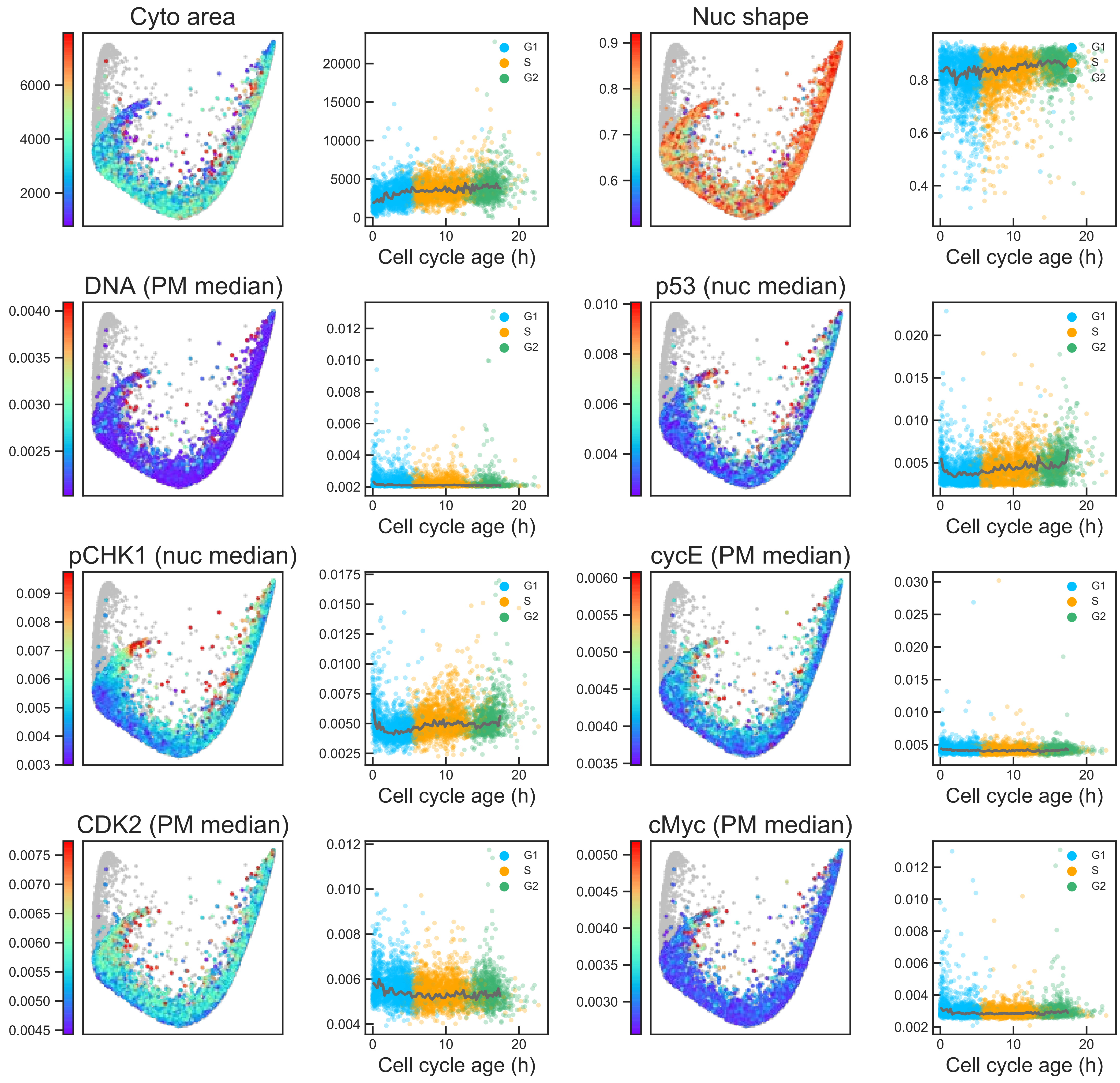

### Aux Fig. S1 - Proliferative trajectory - Page 8

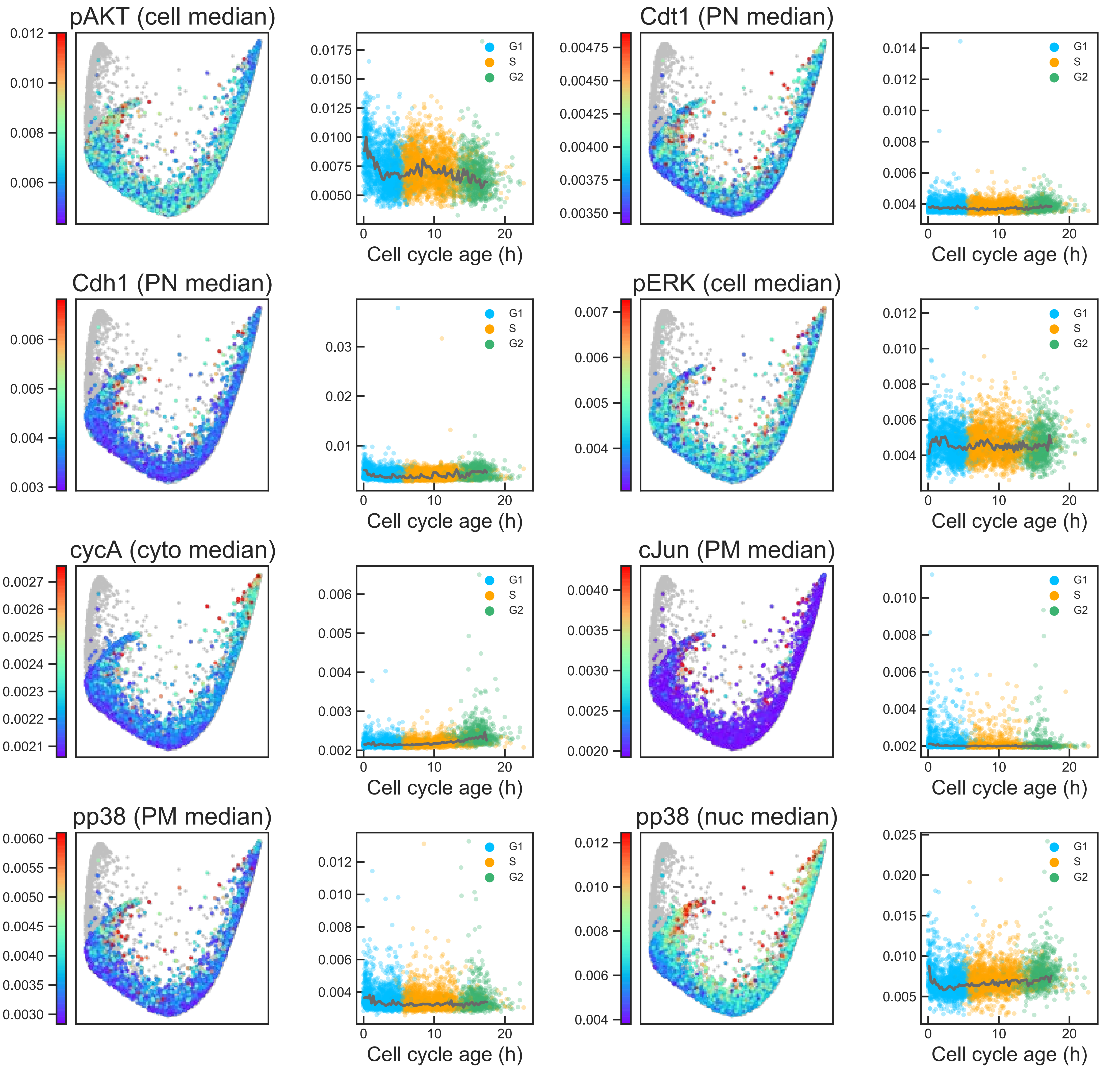

### Aux Fig. S1 - Proliferative trajectory - Page 9

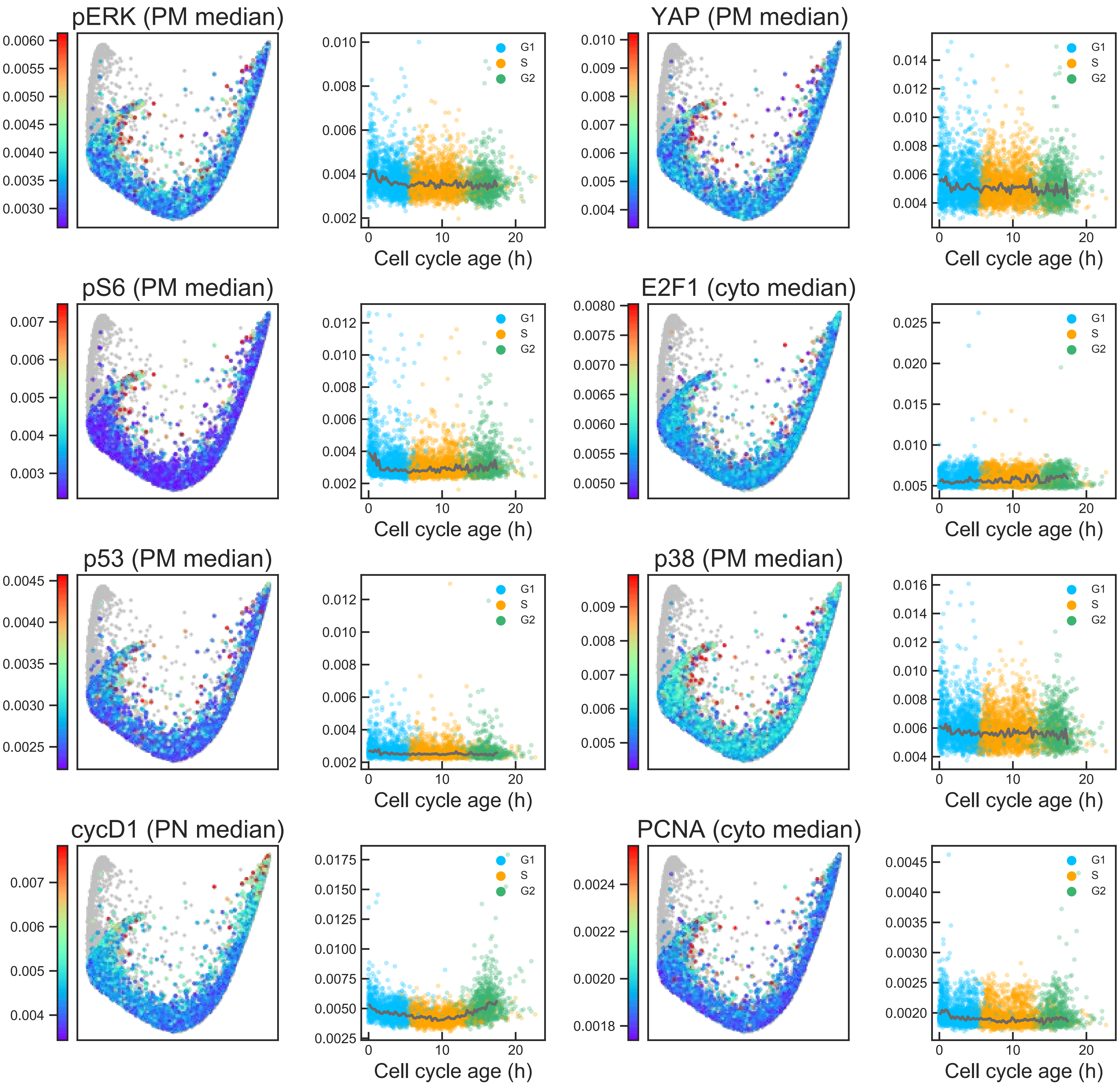

### Aux Fig. S1 - Proliferative trajectory - Page 10

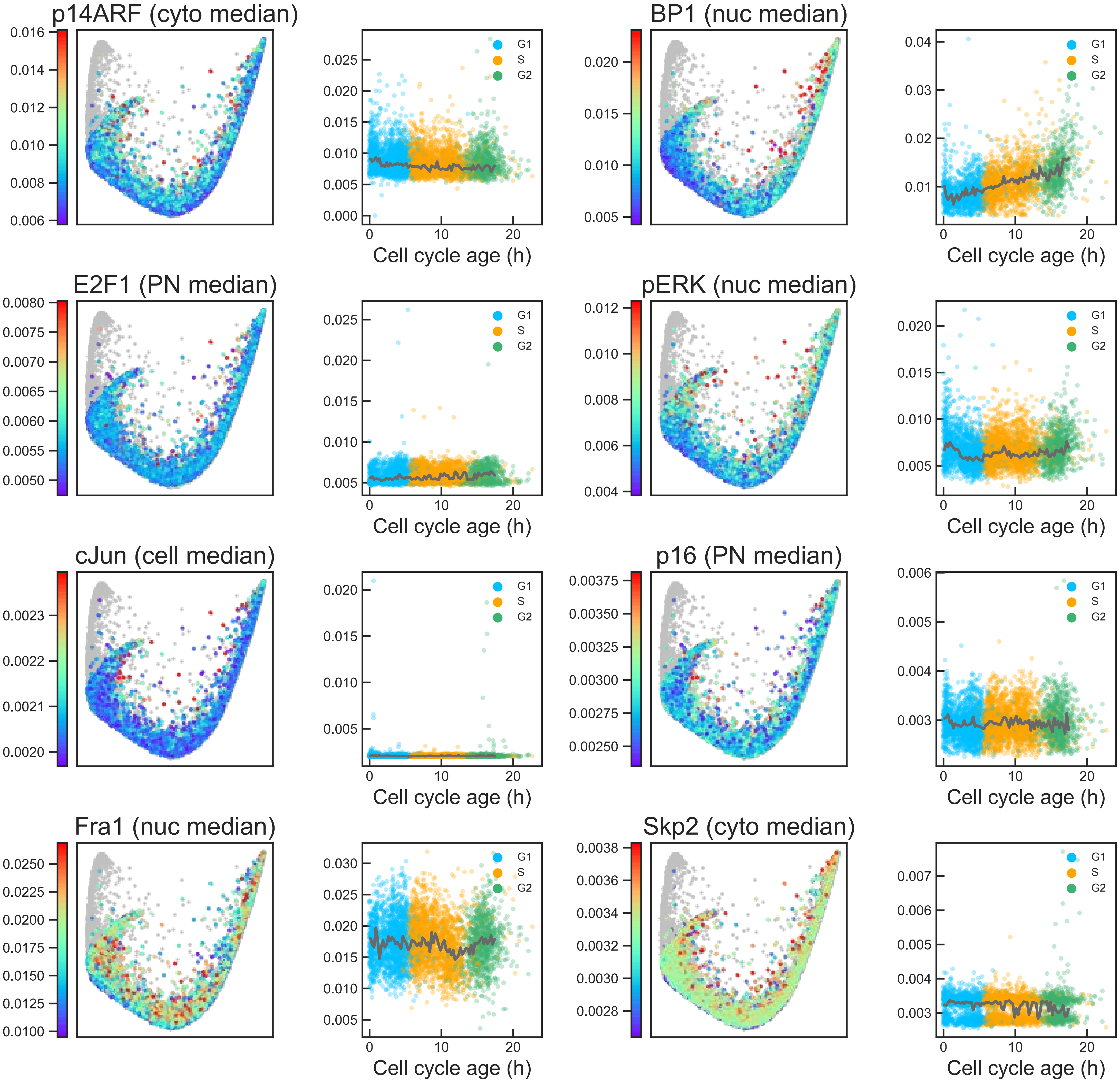

### Aux Fig. S1 - Proliferative trajectory - Page 11

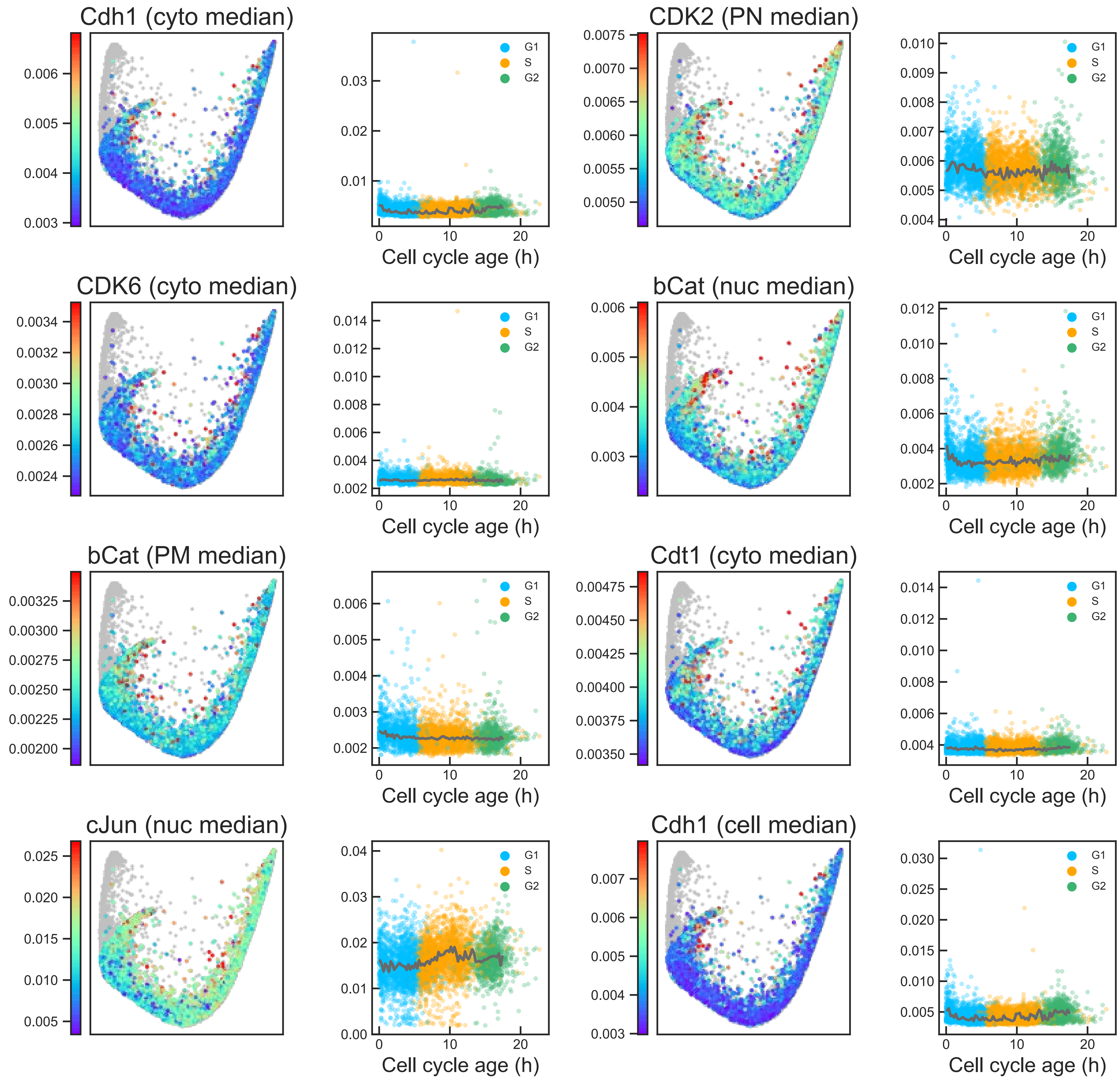

### Aux Fig. S1 - Proliferative trajectory - Page 12

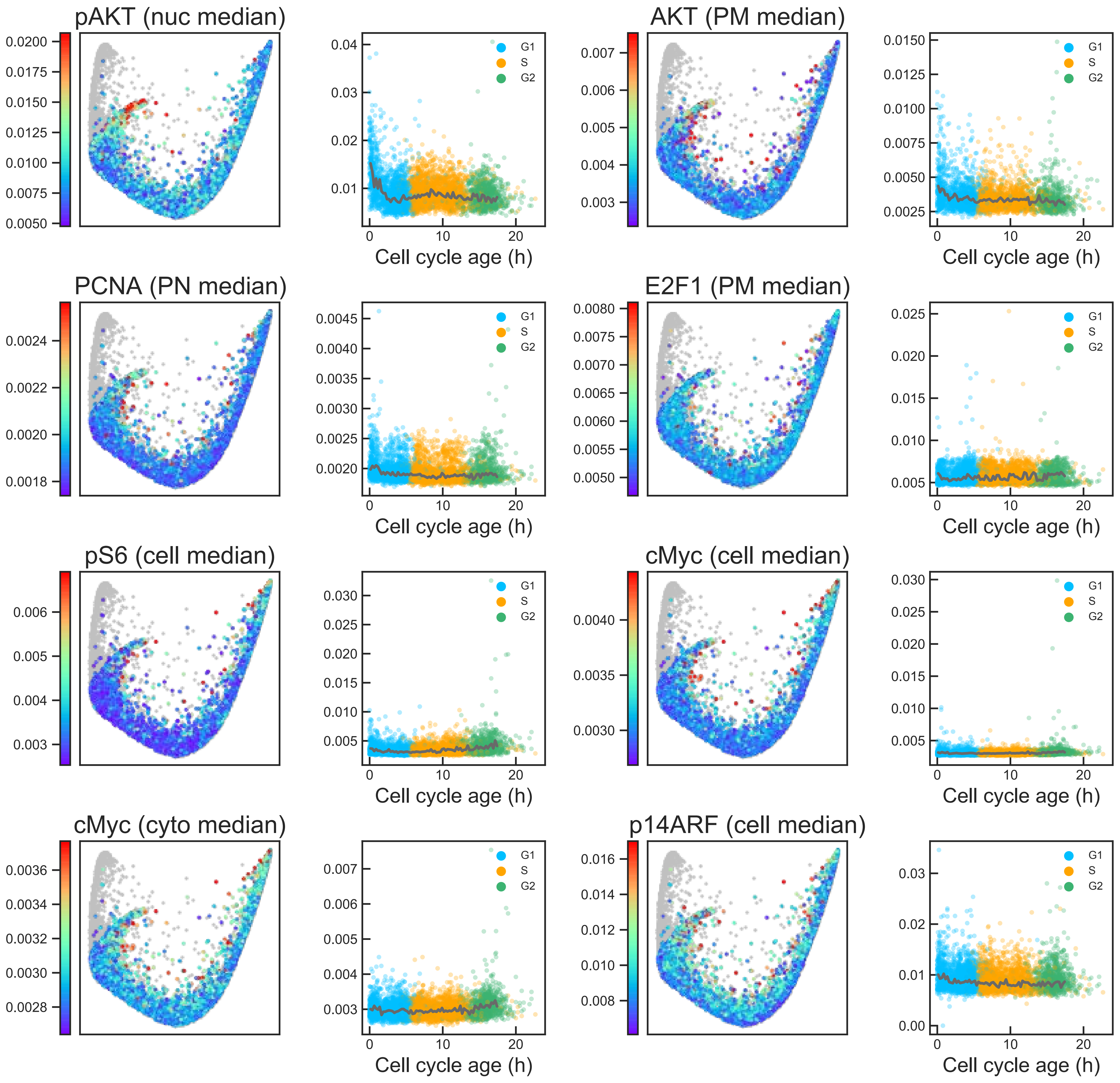

#### Aux Fig. S1 - Proliferative trajectory - Page 13

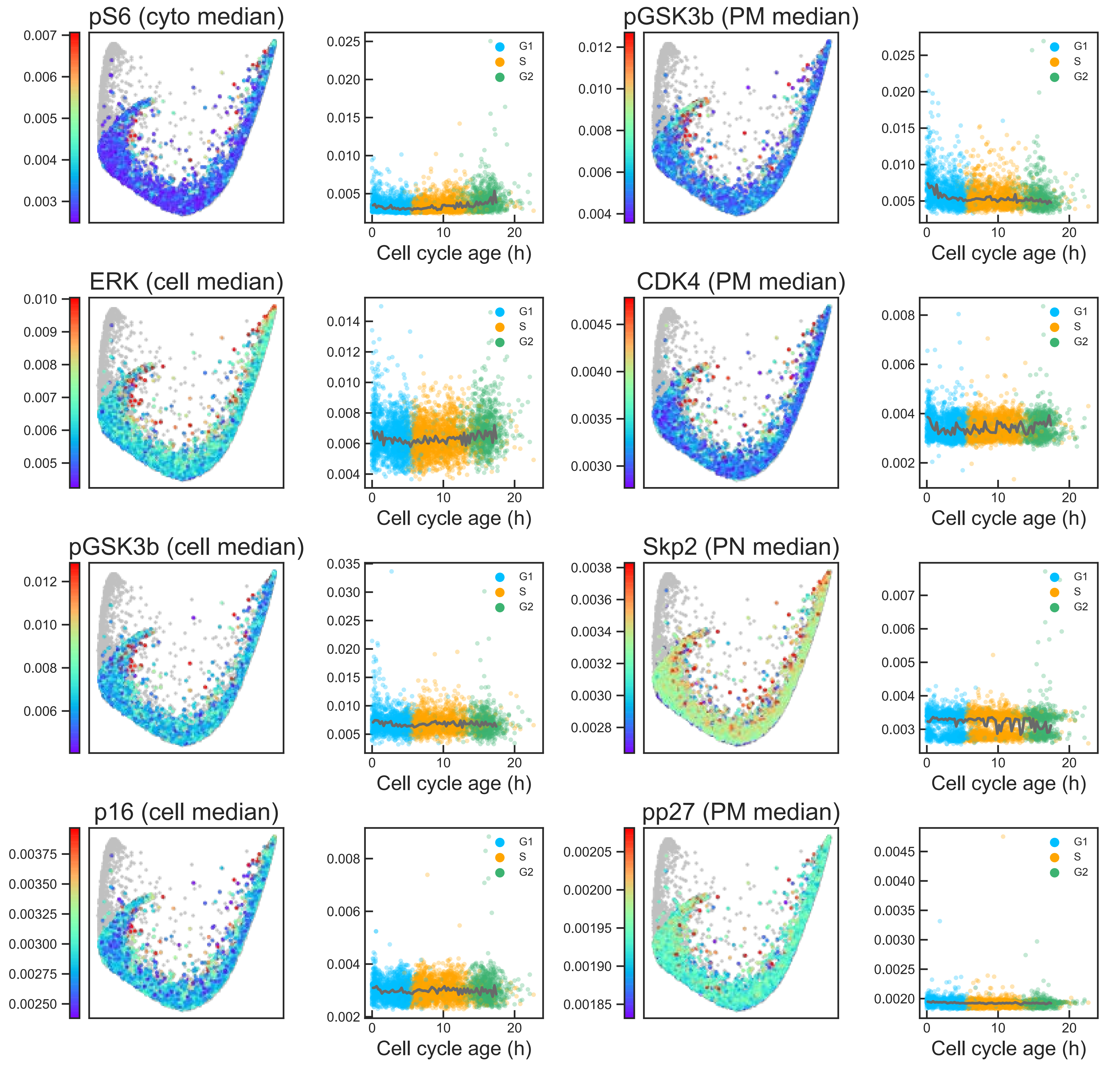

### Aux Fig. S1 - Proliferative trajectory - Page 14

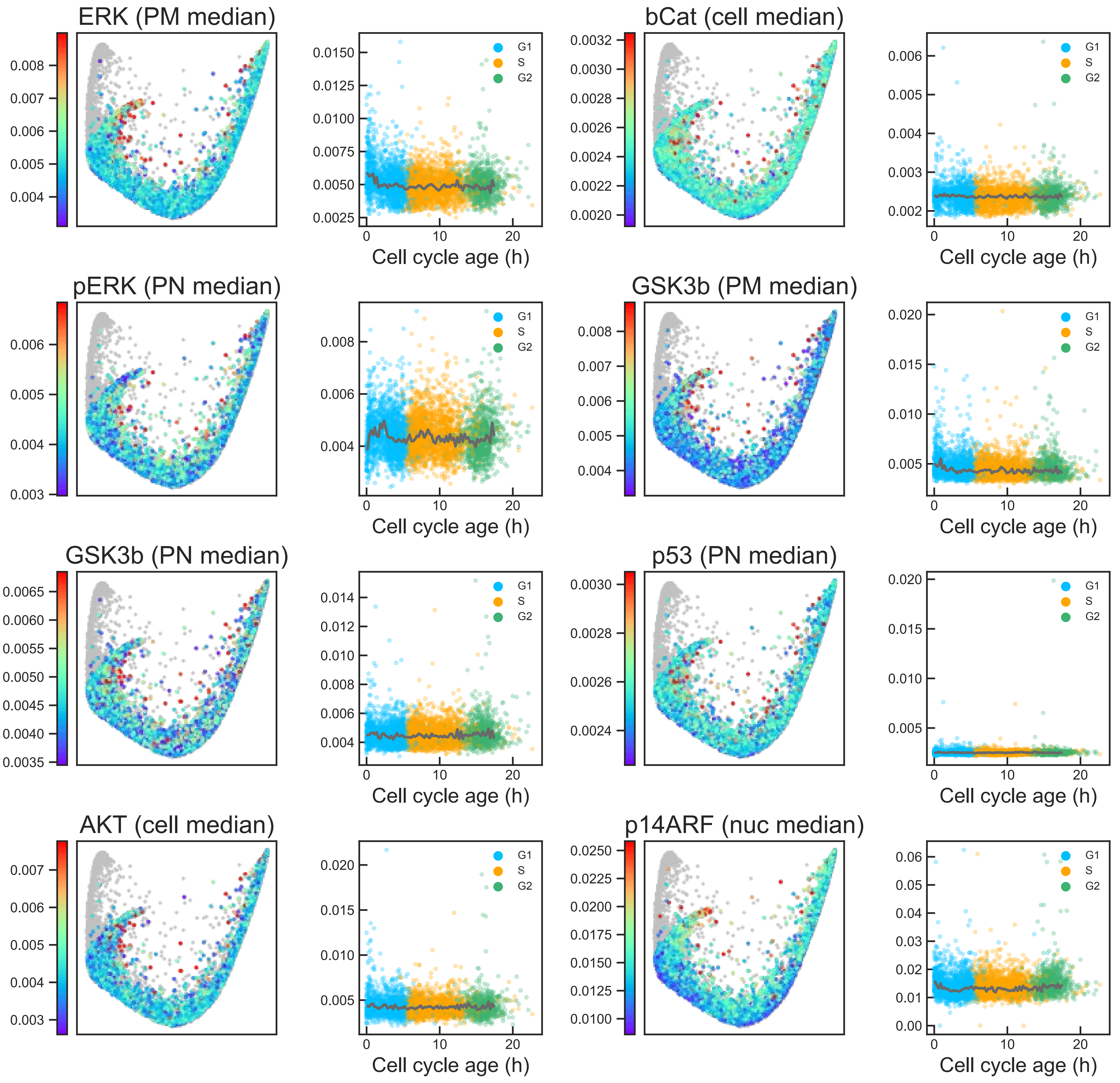

### Aux Fig. S1 - Proliferative trajectory - Page 15

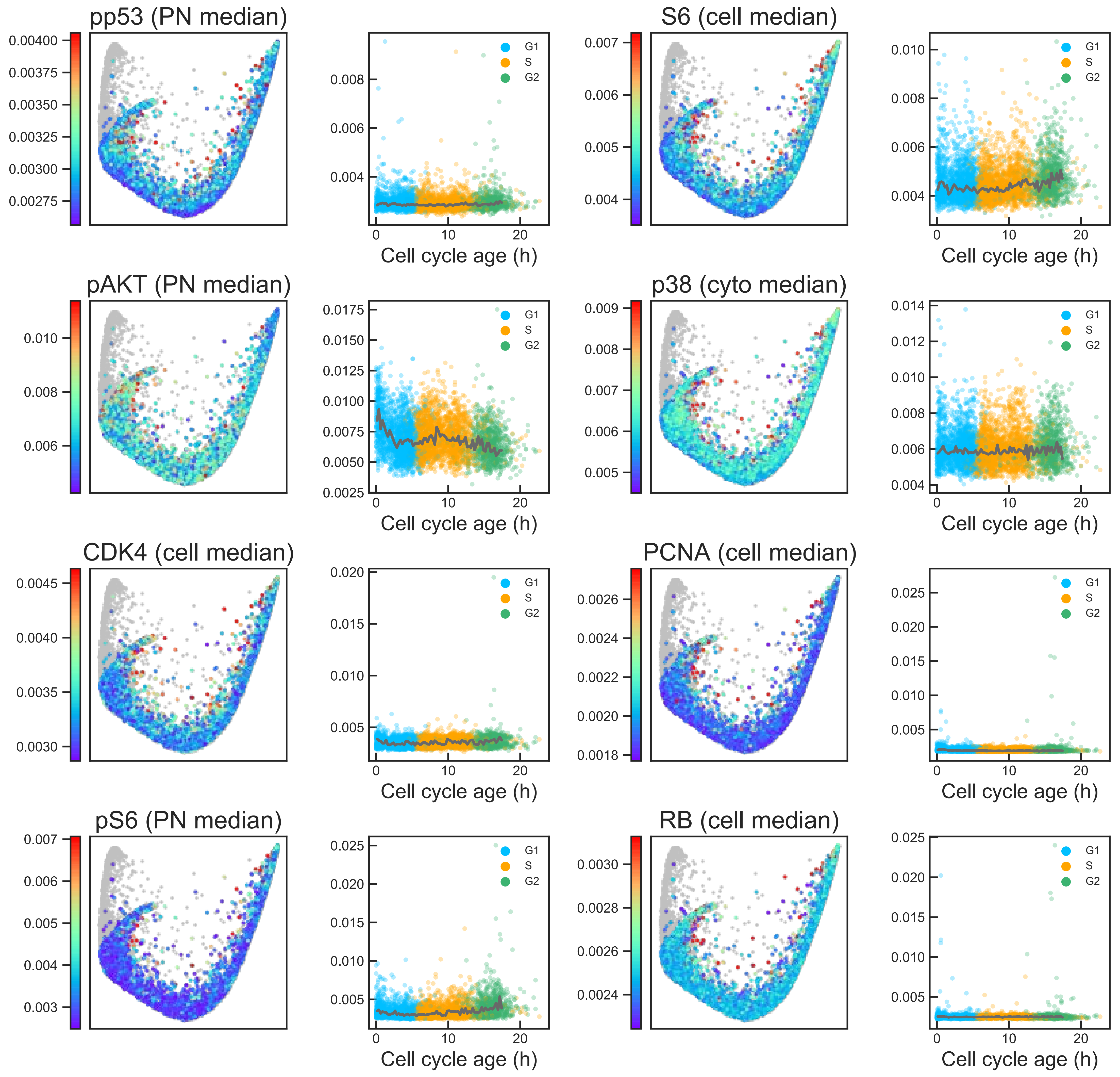

### Aux Fig. S1 - Proliferative trajectory - Page 16

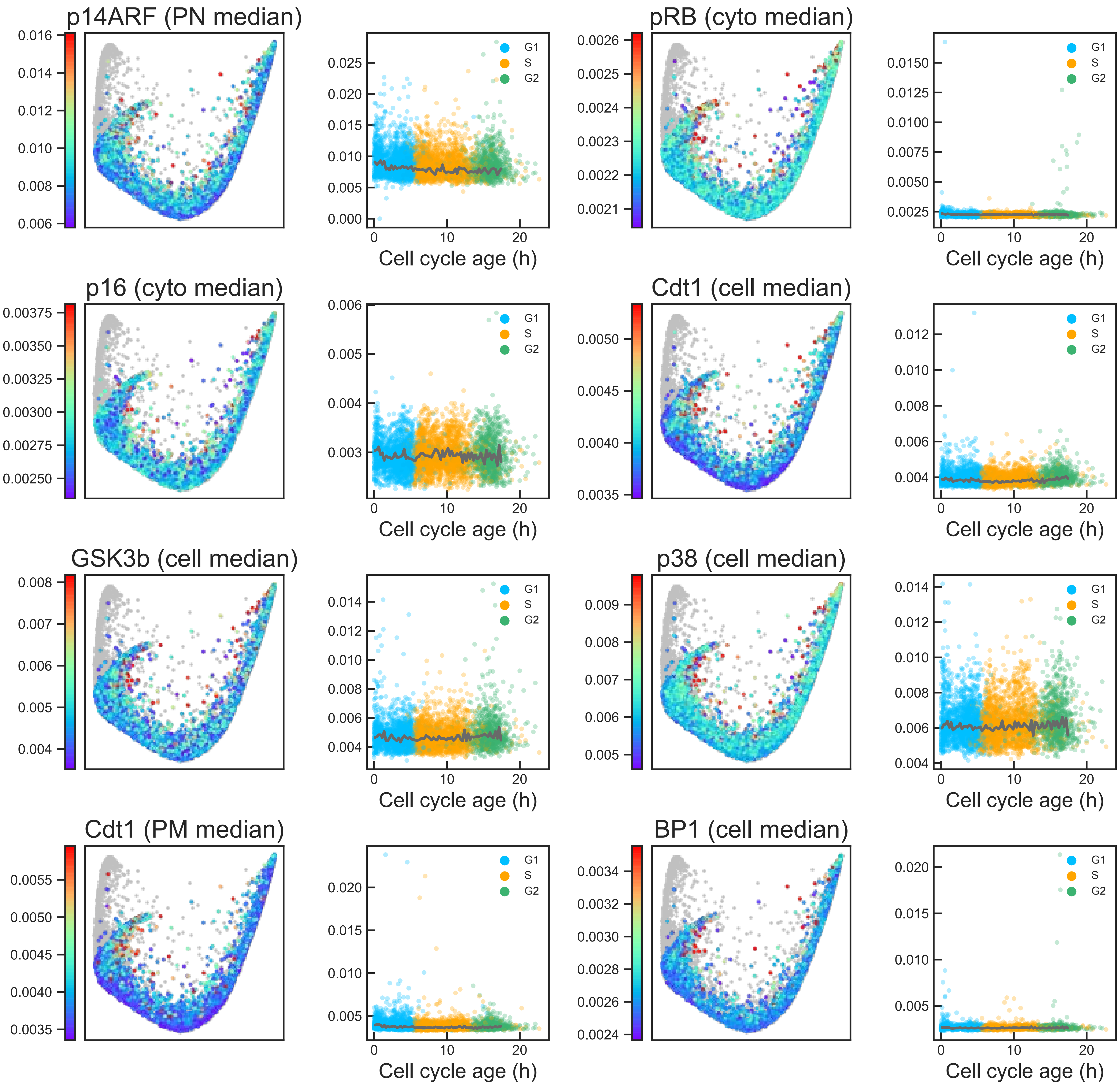

### Aux Fig. S1 - Proliferative trajectory - Page 17

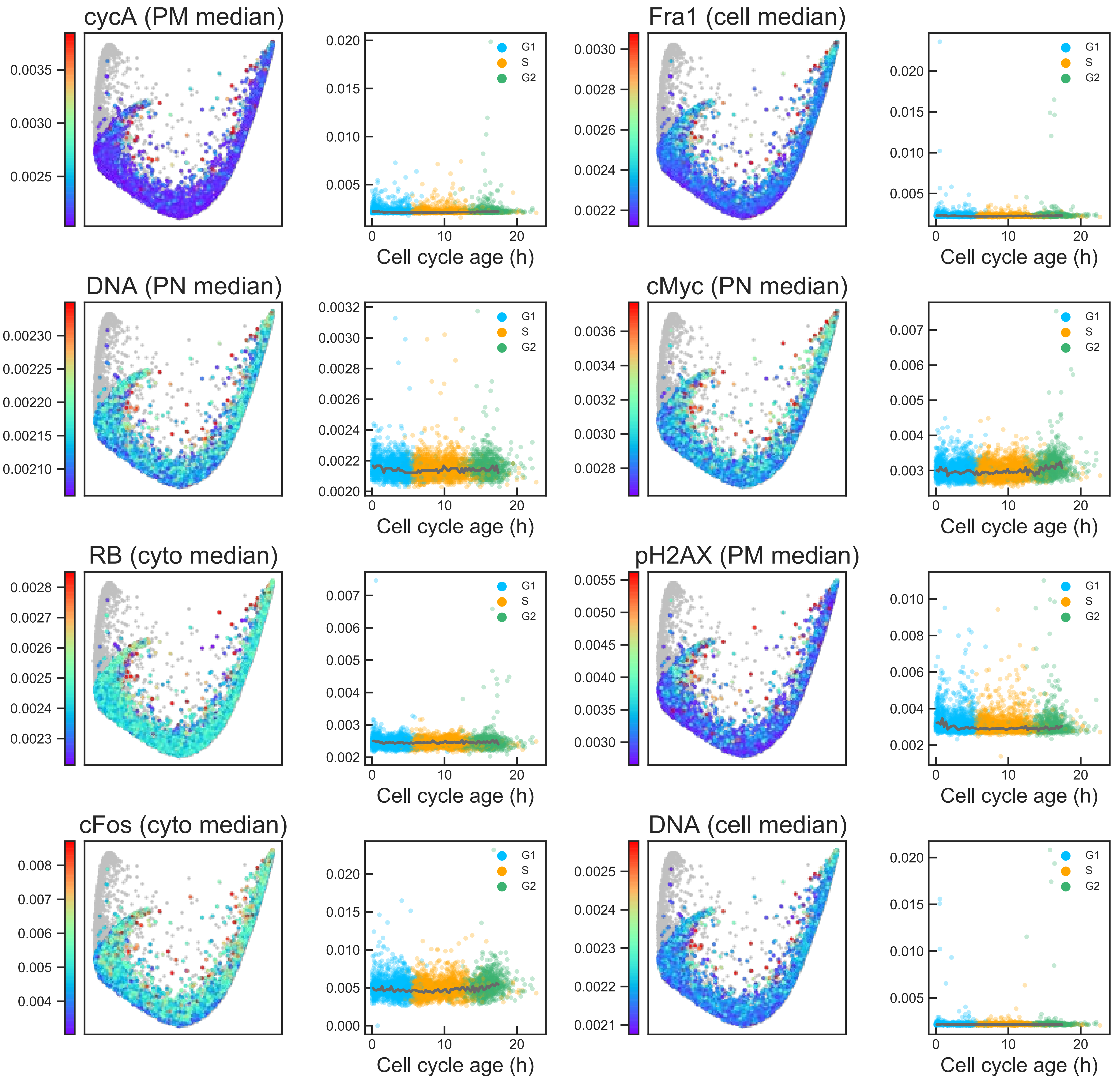

### Aux Fig. S1 - Proliferative trajectory - Page 18

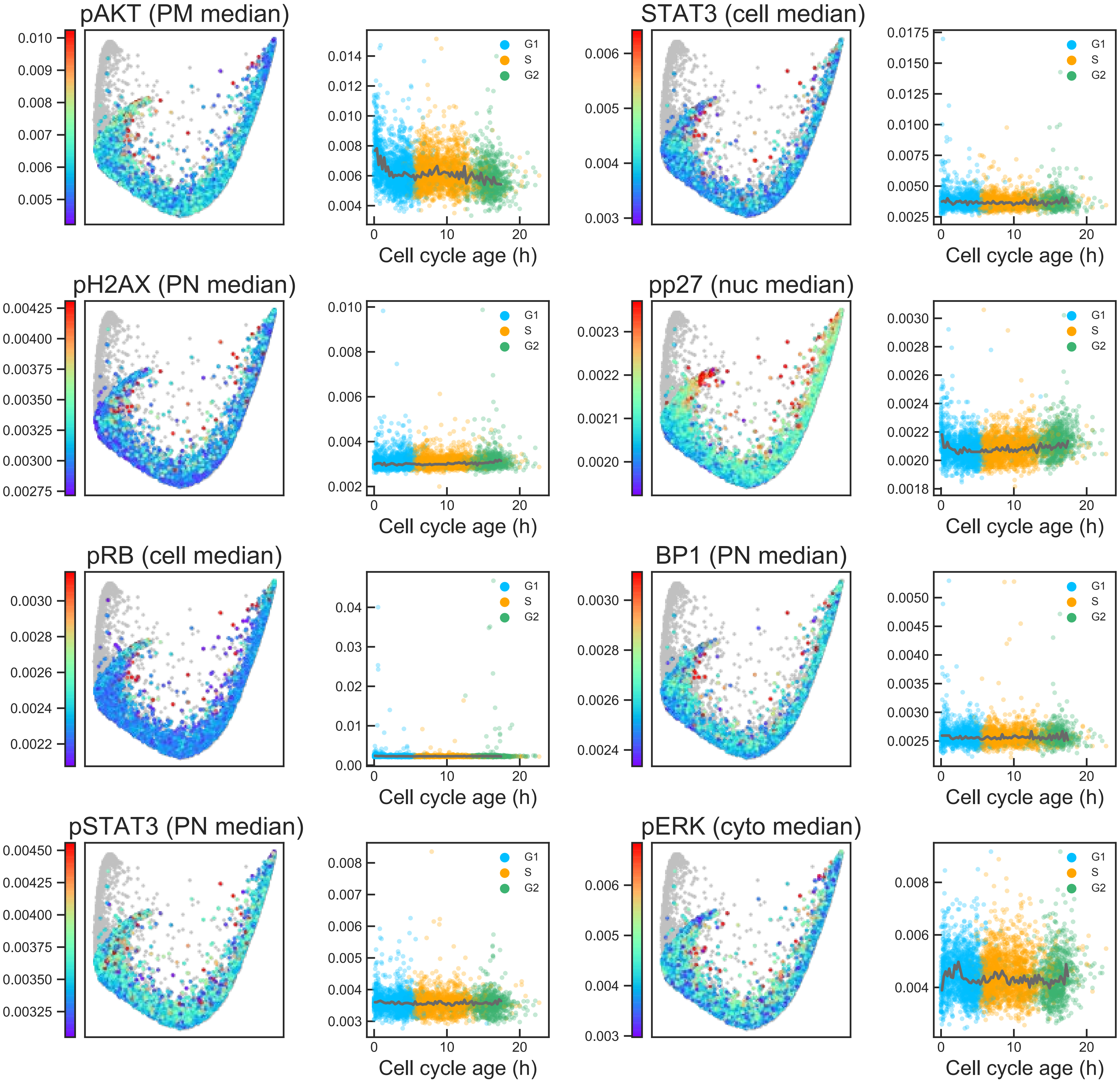

### Aux Fig. S1 - Proliferative trajectory - Page 19

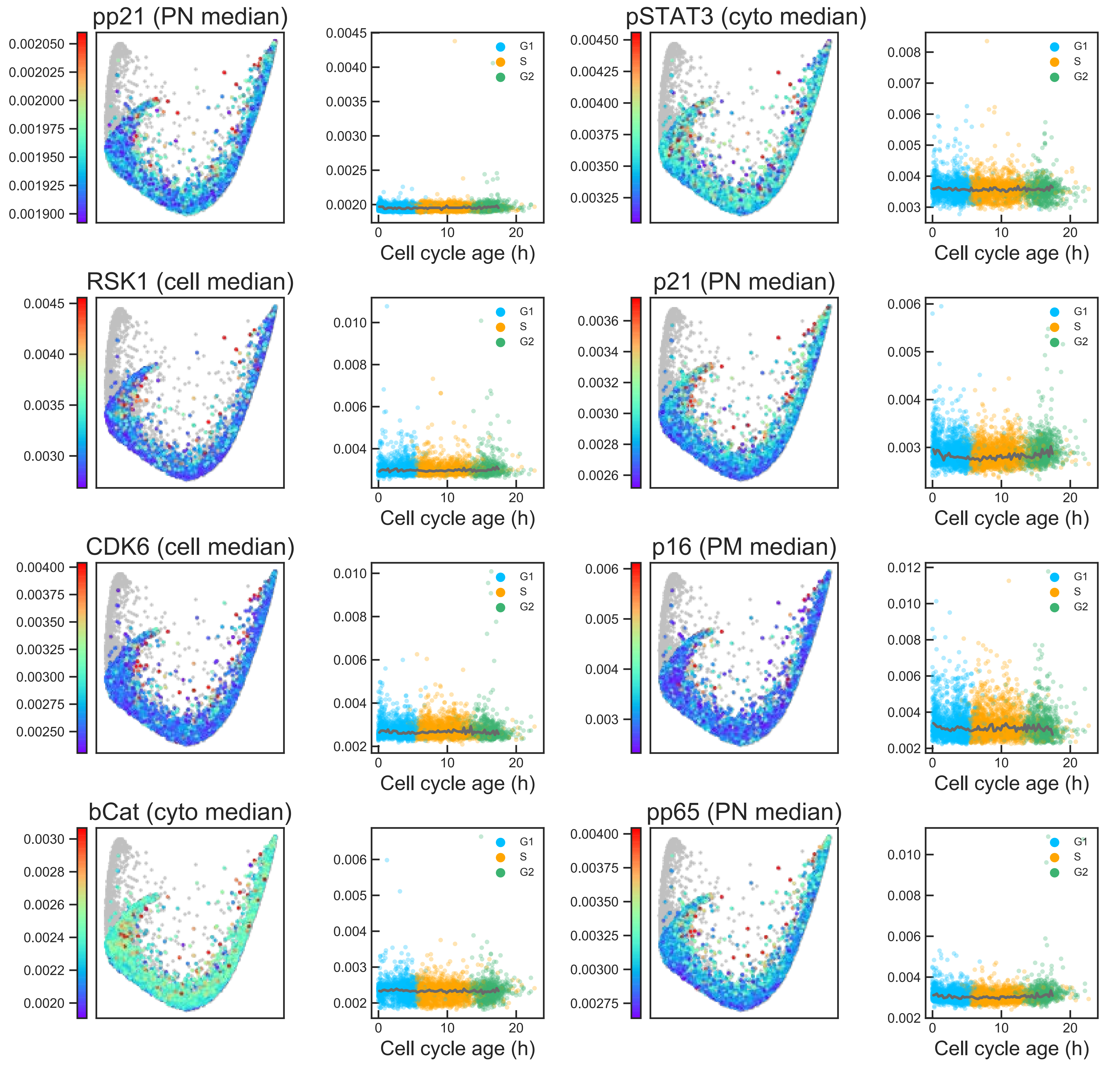

### Aux Fig. S1 - Proliferative trajectory - Page 20

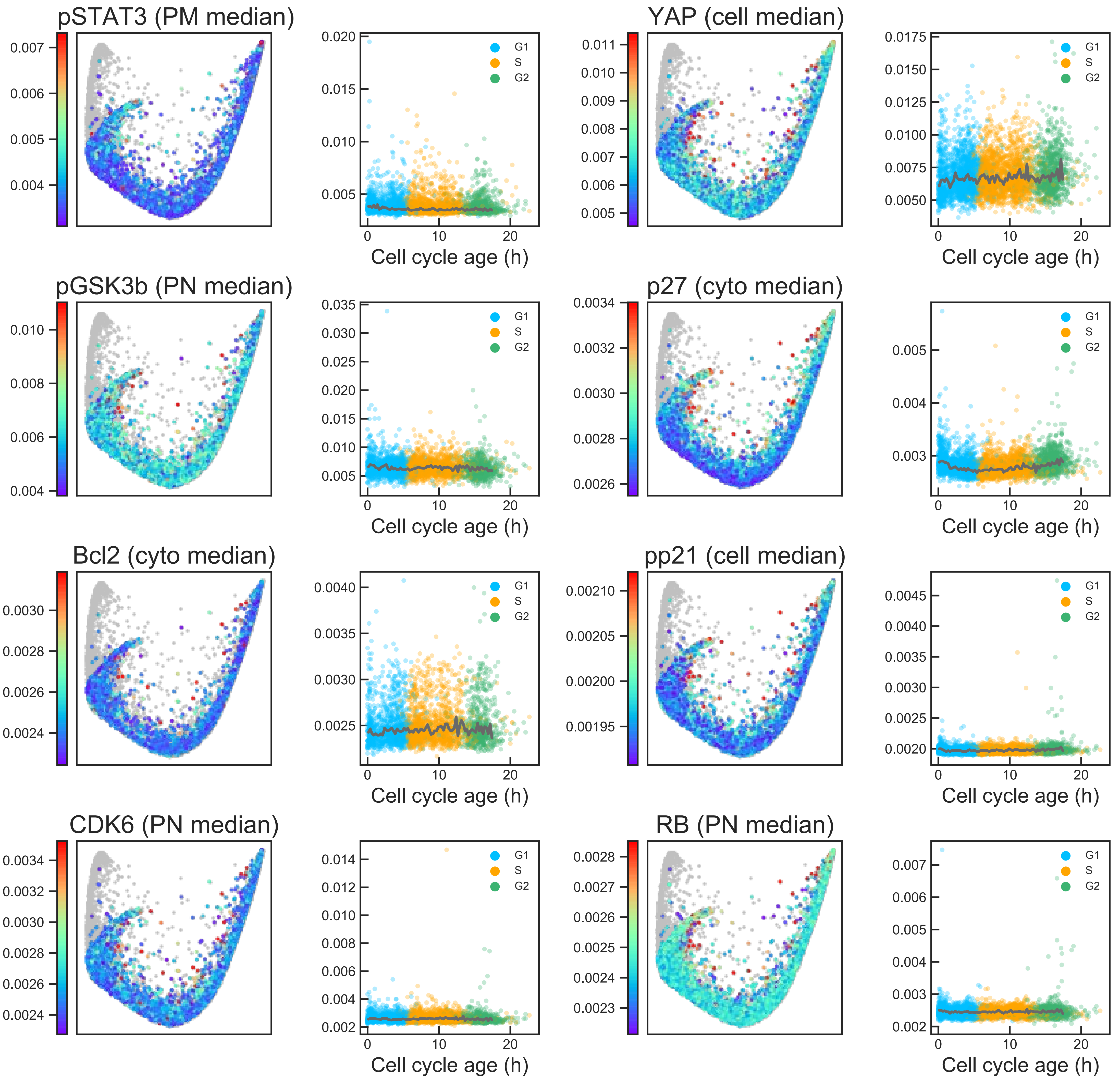

**Aux Fig. S1 - Proliferative trajectory - Page 21**

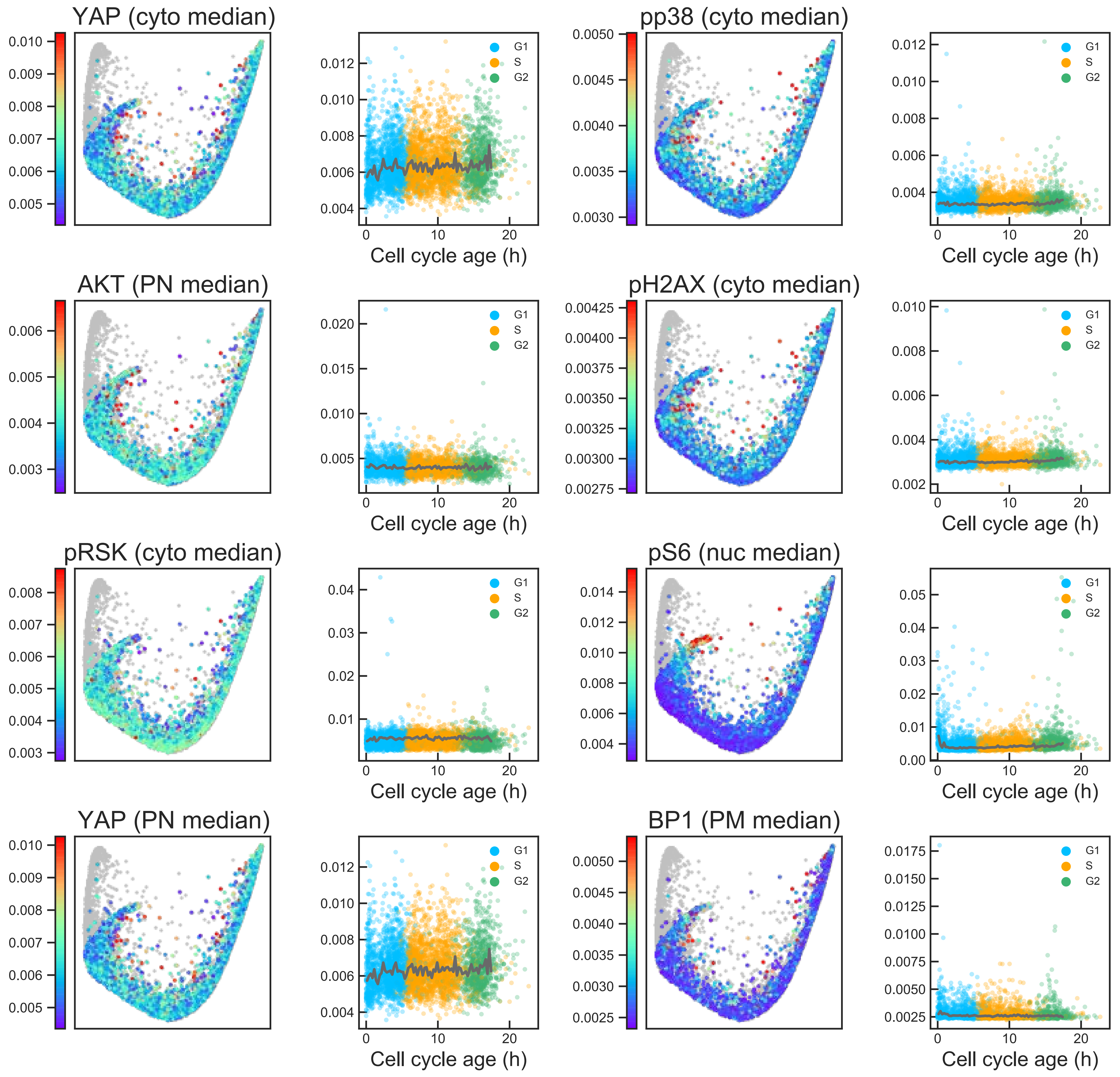

### Aux Fig. S1 - Proliferative trajectory - Page 22

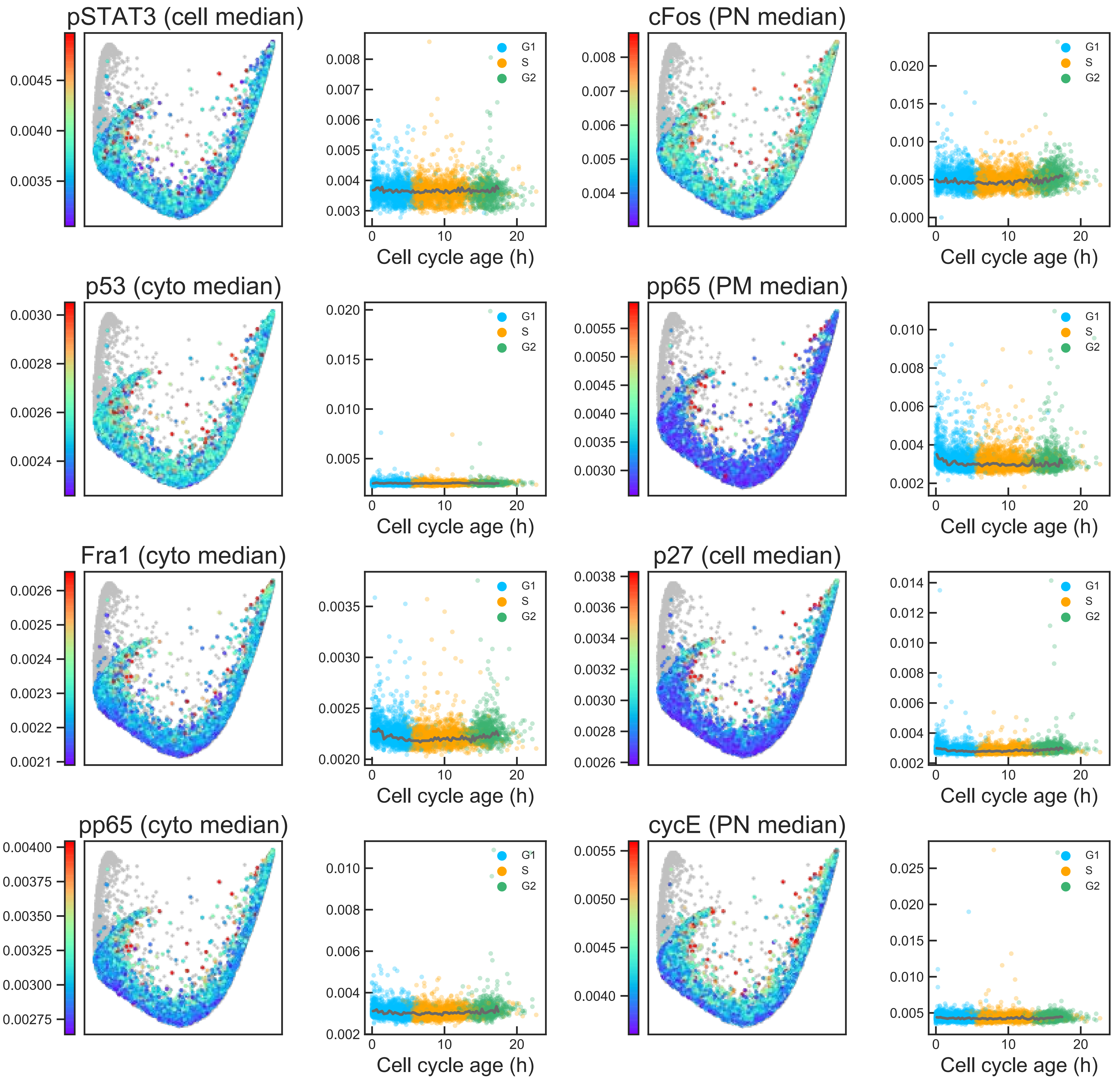

### Aux Fig. S1 - Proliferative trajectory - Page 23

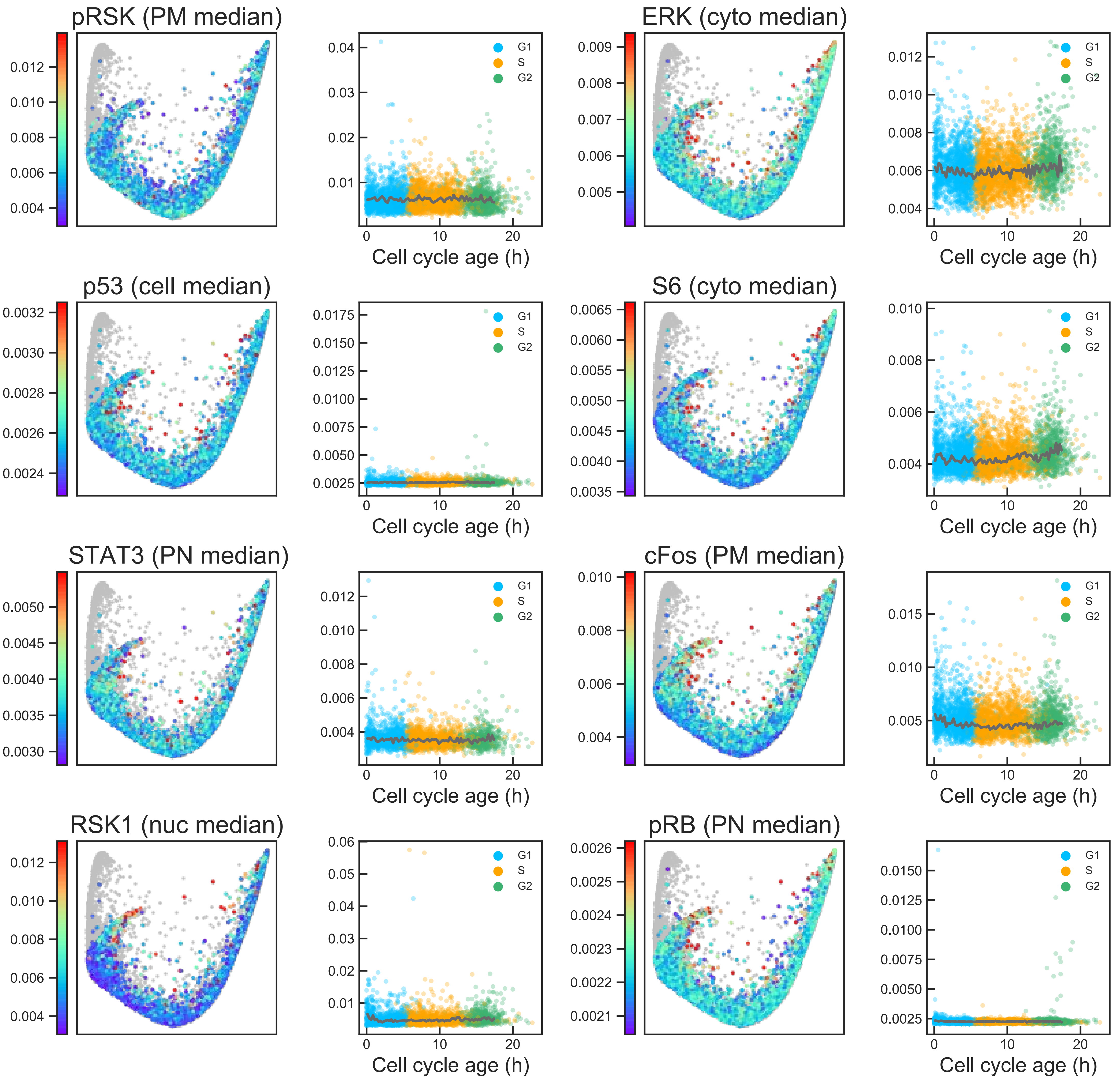

### Aux Fig. S1 - Proliferative trajectory - Page 24

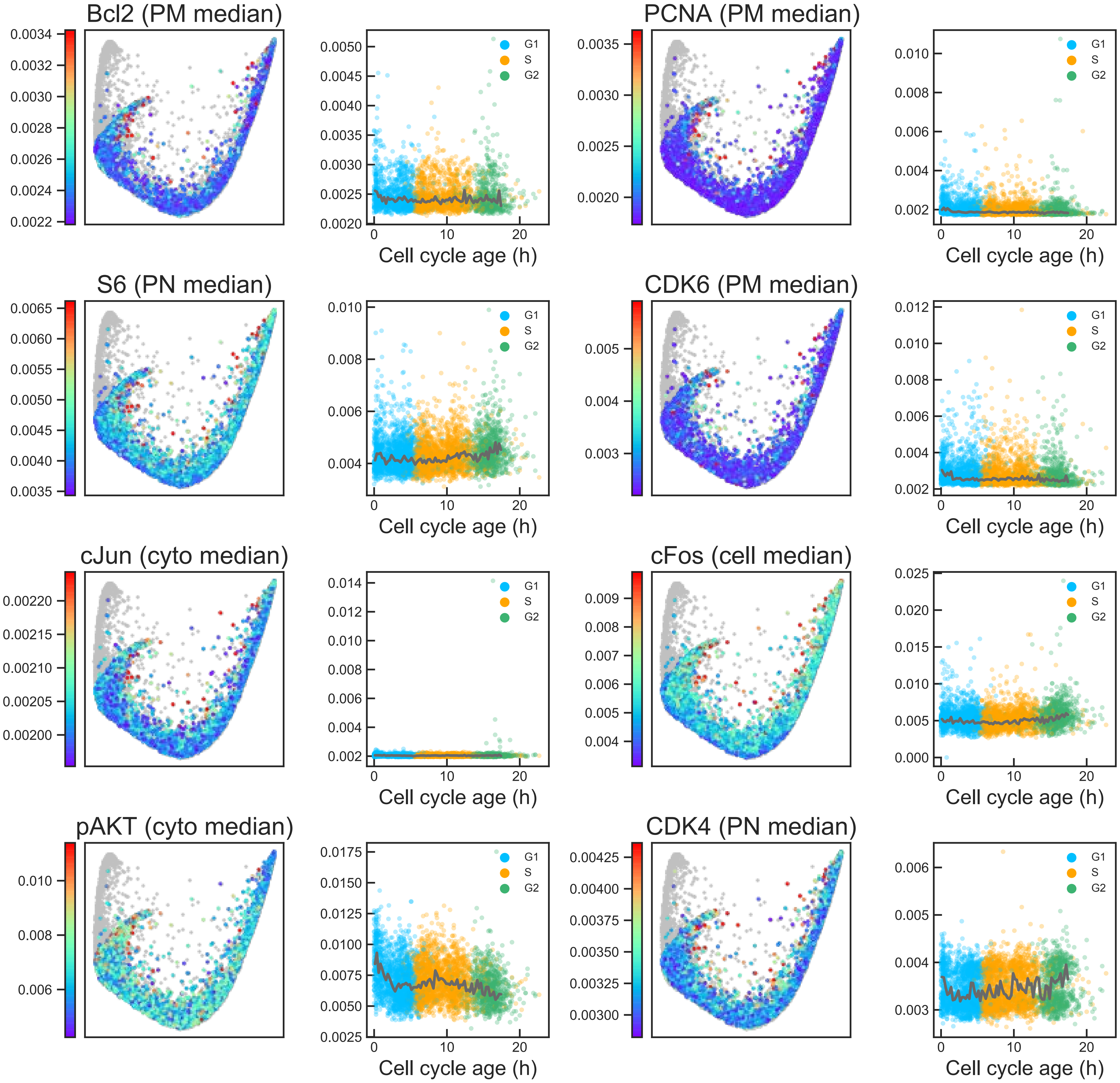

#### Aux Fig. S1 - Proliferative trajectory - Page 25

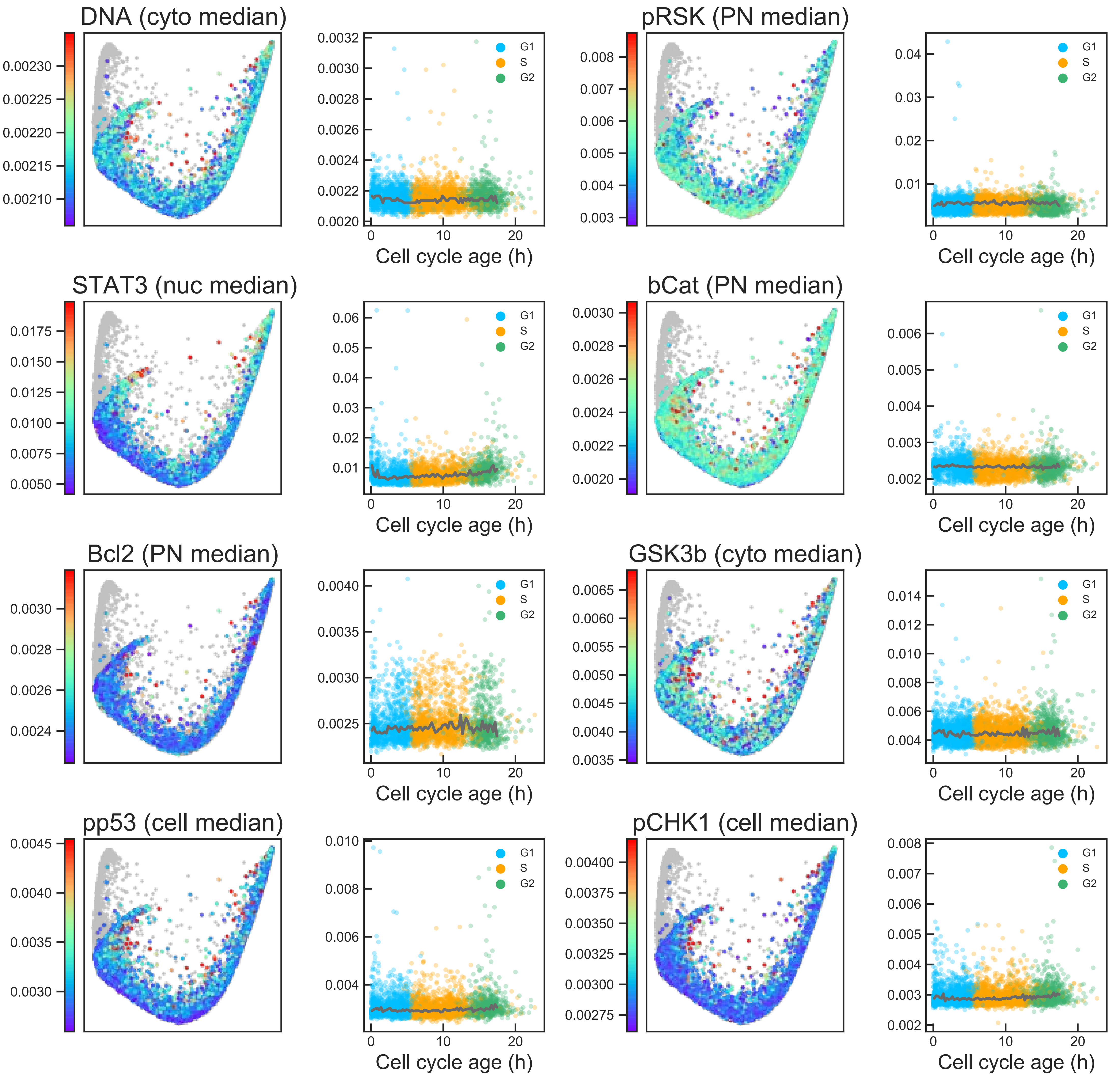

### Aux Fig. S1 - Proliferative trajectory - Page 26

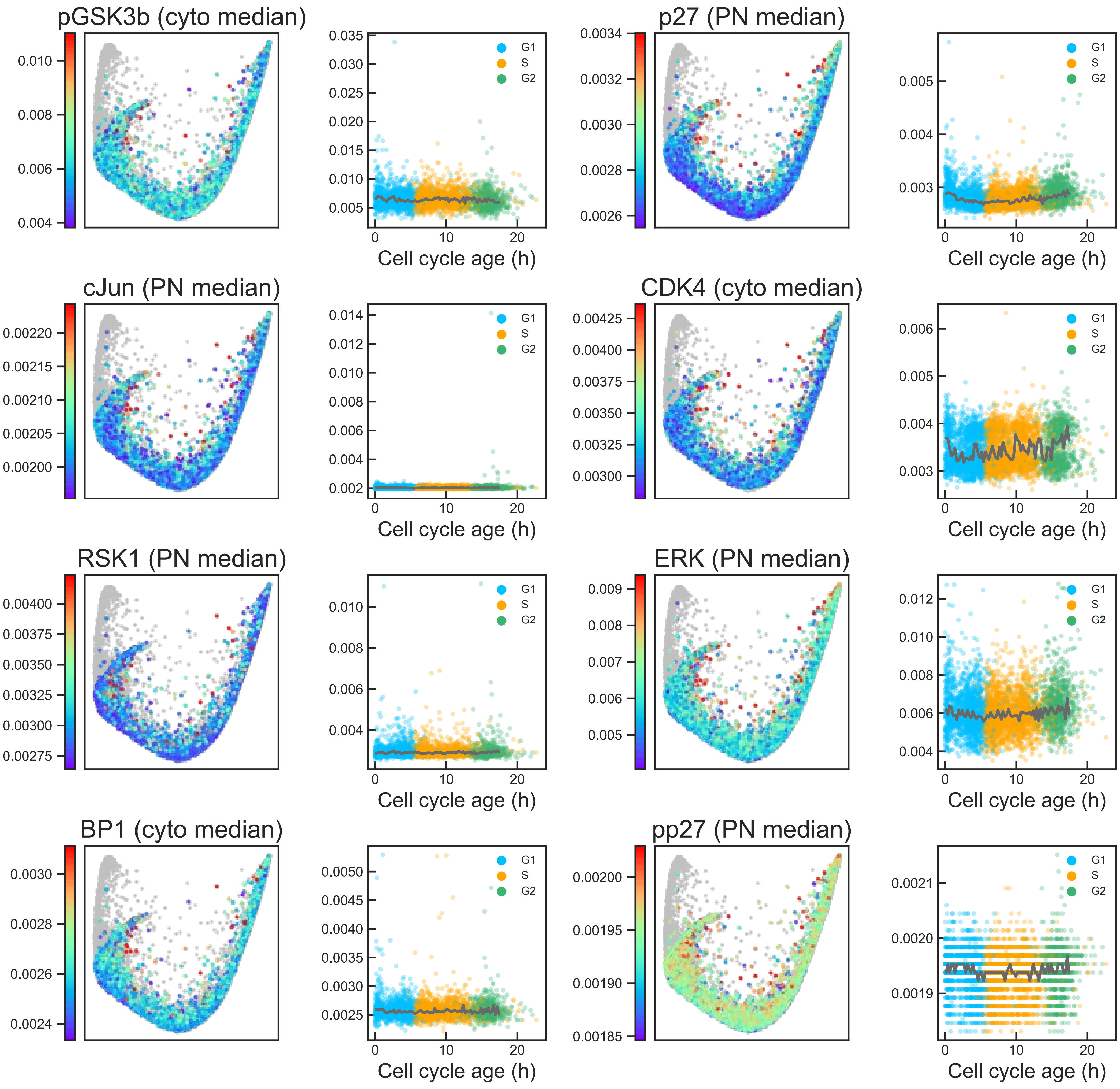

#### Aux Fig. S1 - Proliferative trajectory - Page 27

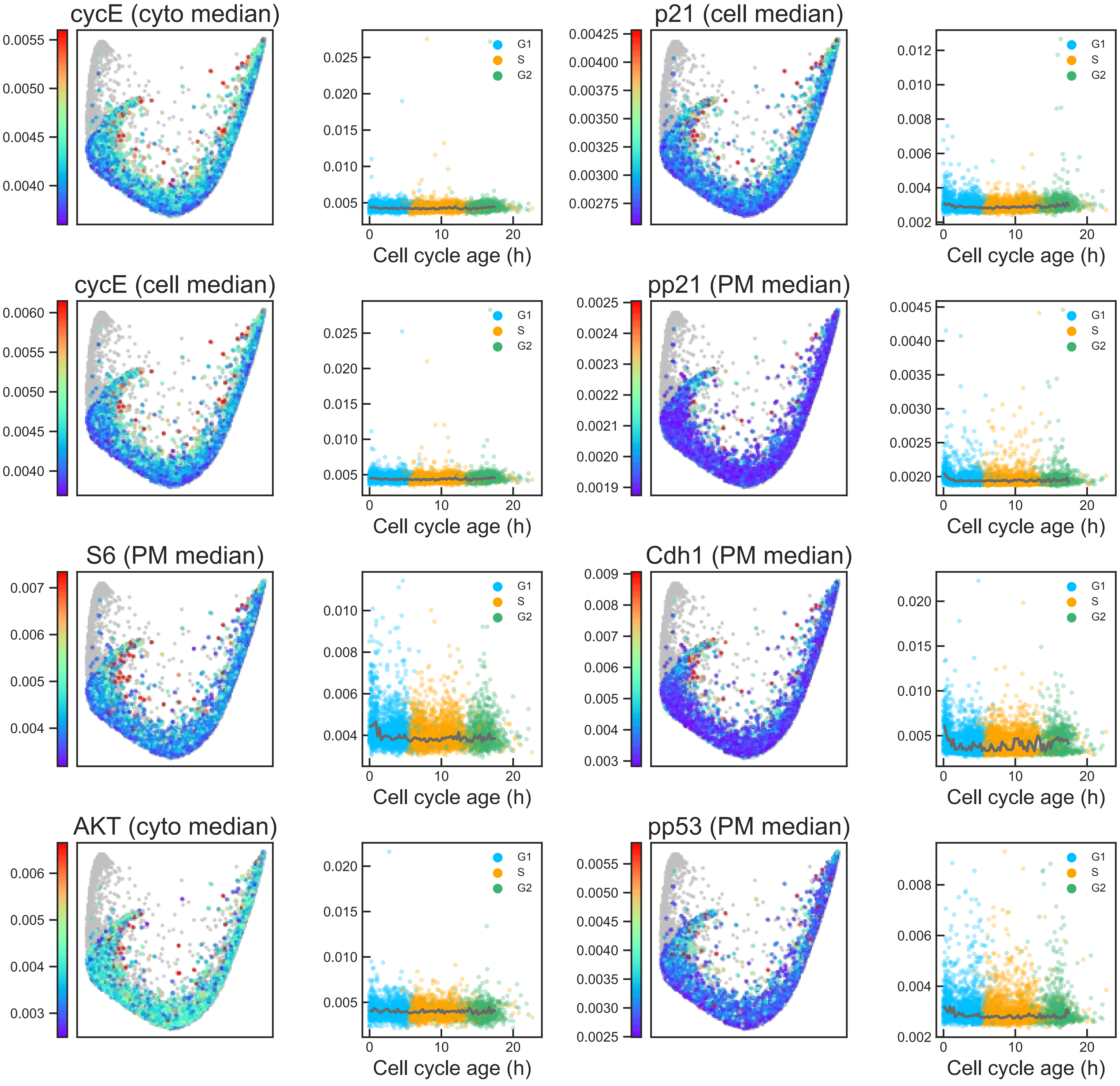

### Aux Fig. S1 - Proliferative trajectory - Page 28

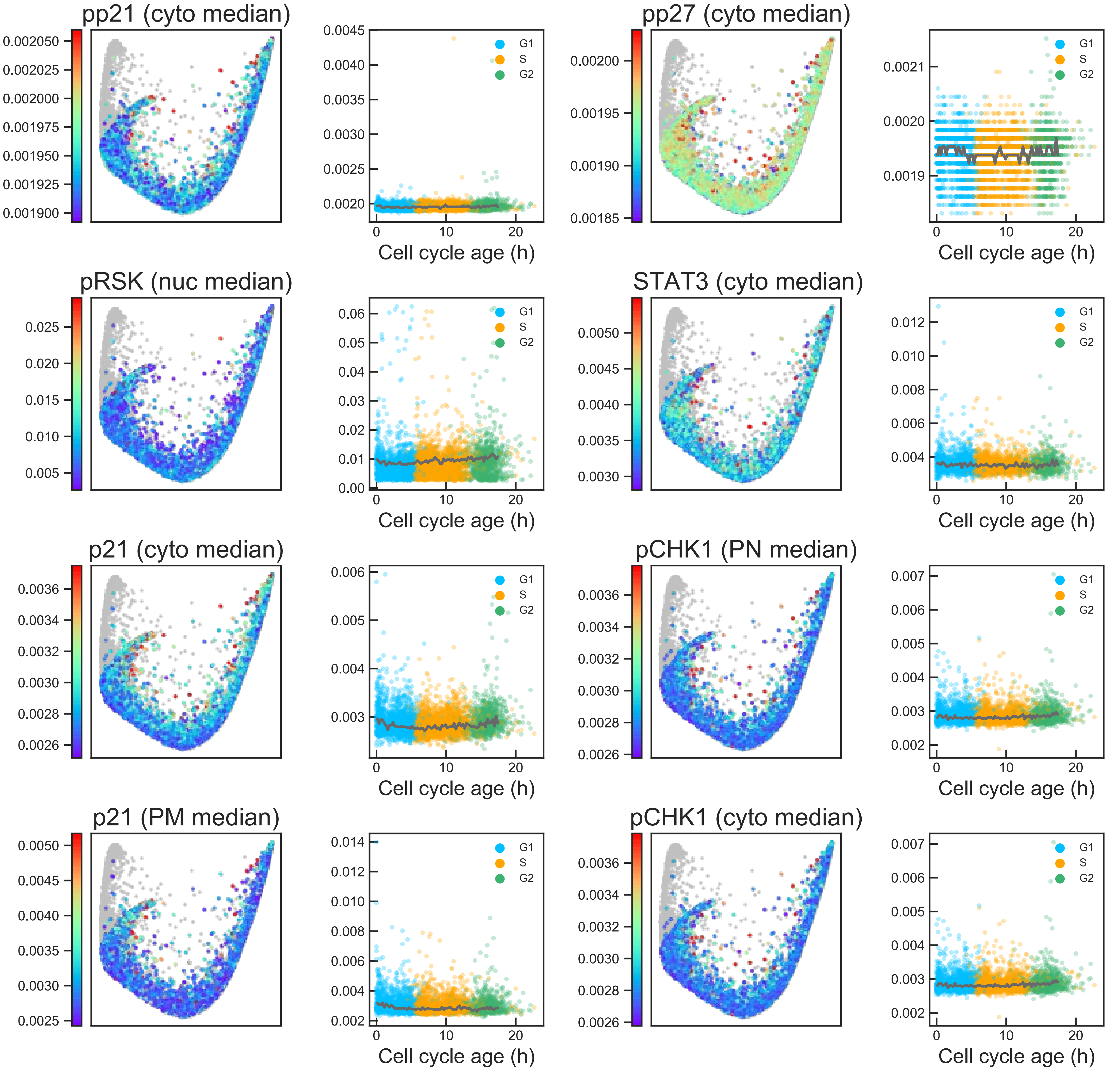

### Aux Fig. S1 - Proliferative trajectory - Page 29

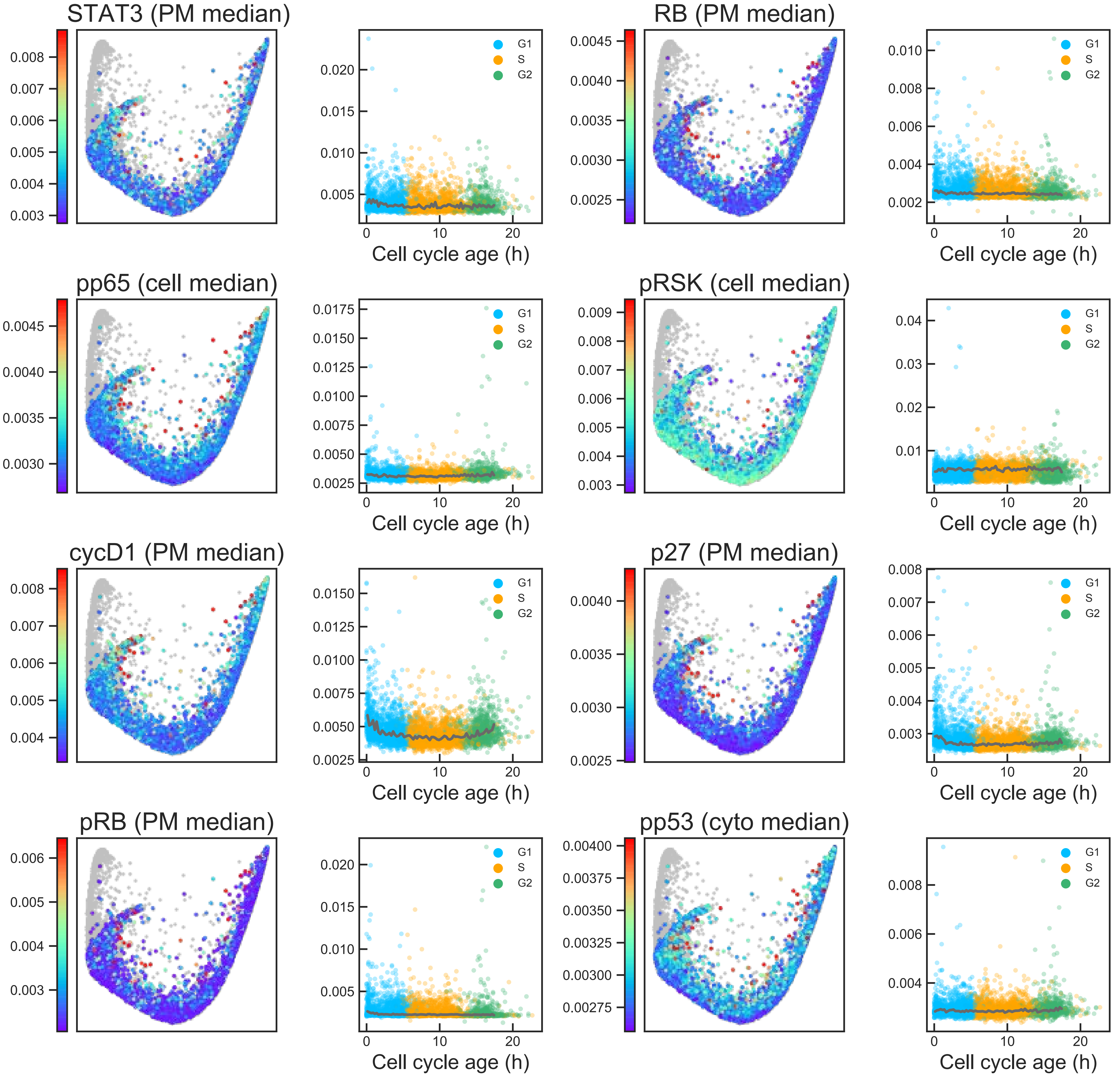

### Aux Fig. S1 - Proliferative trajectory - Page 30

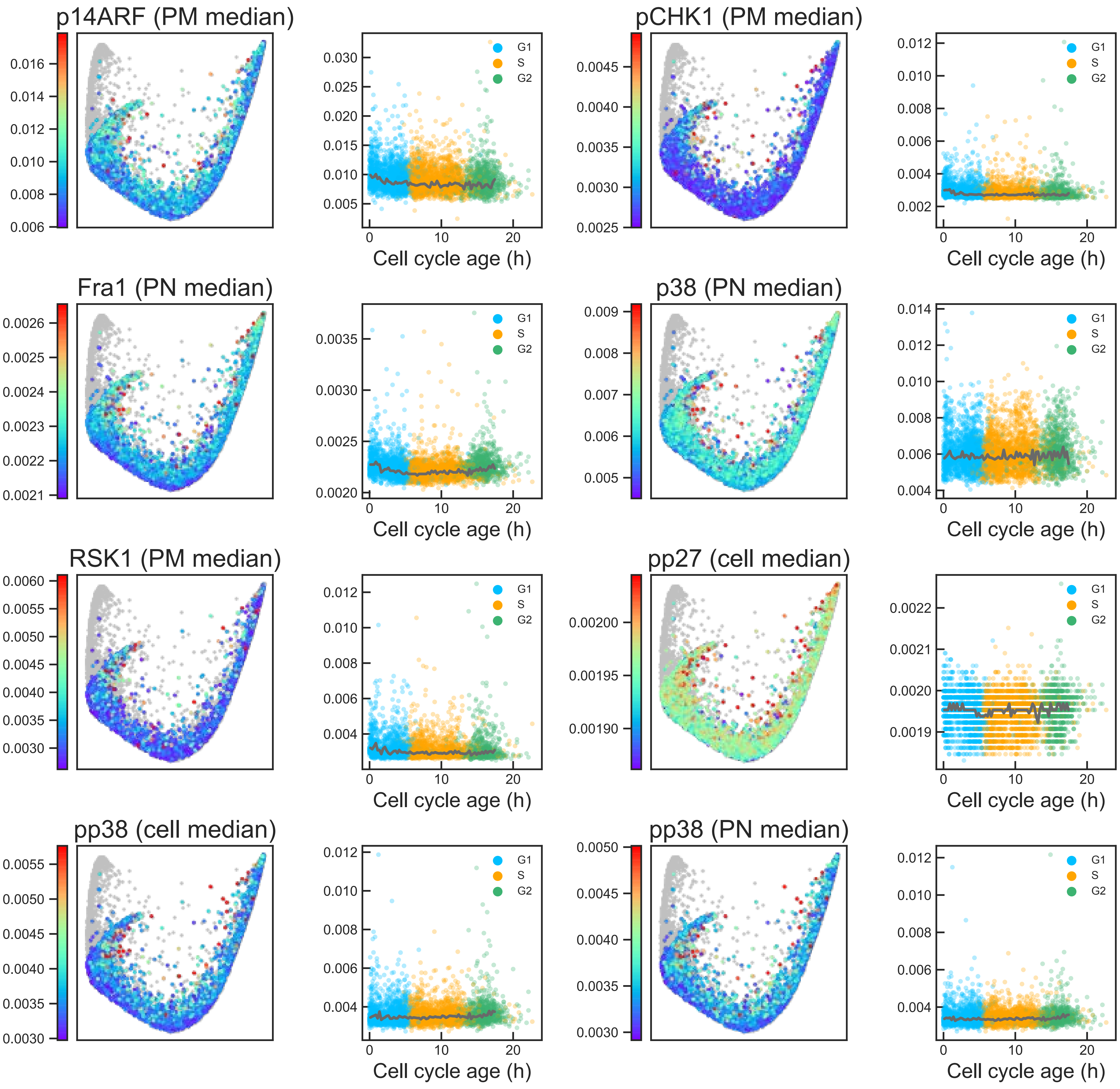

#### Aux Fig. S1 - Proliferative trajectory - Page 31

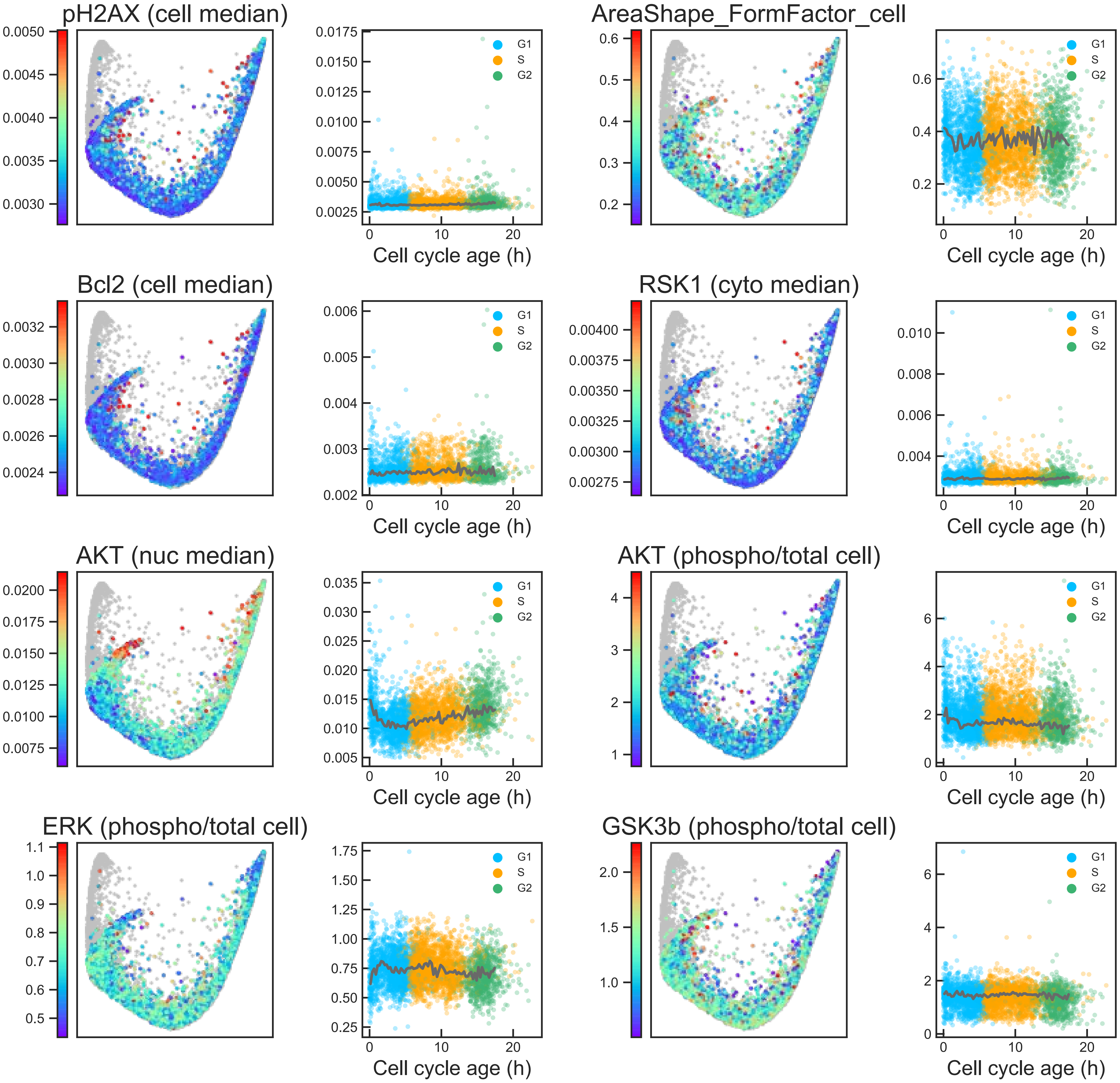

### Aux Fig. S1 - Proliferative trajectory - Page 32

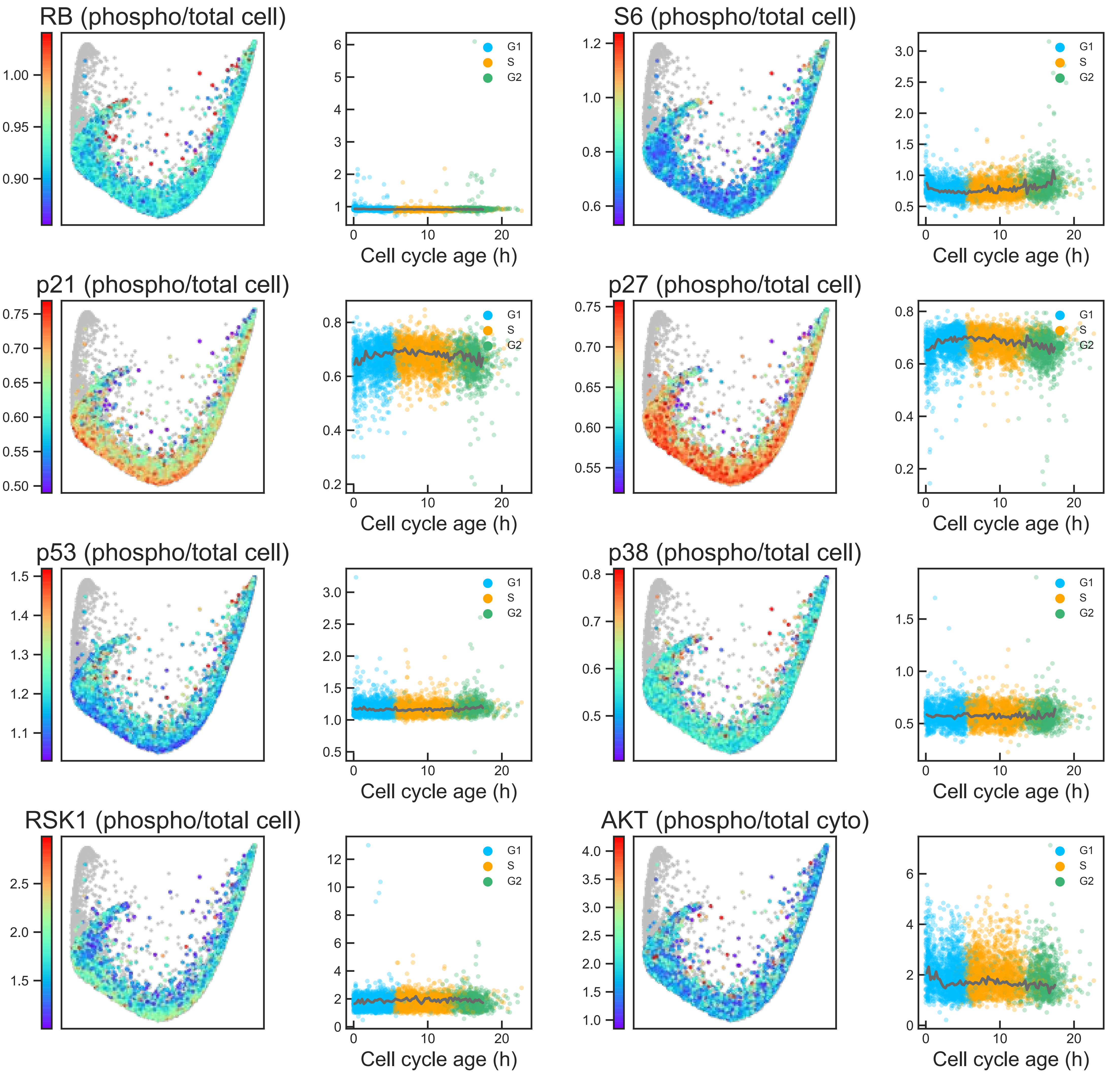

### Aux Fig. S1 - Proliferative trajectory - Page 33

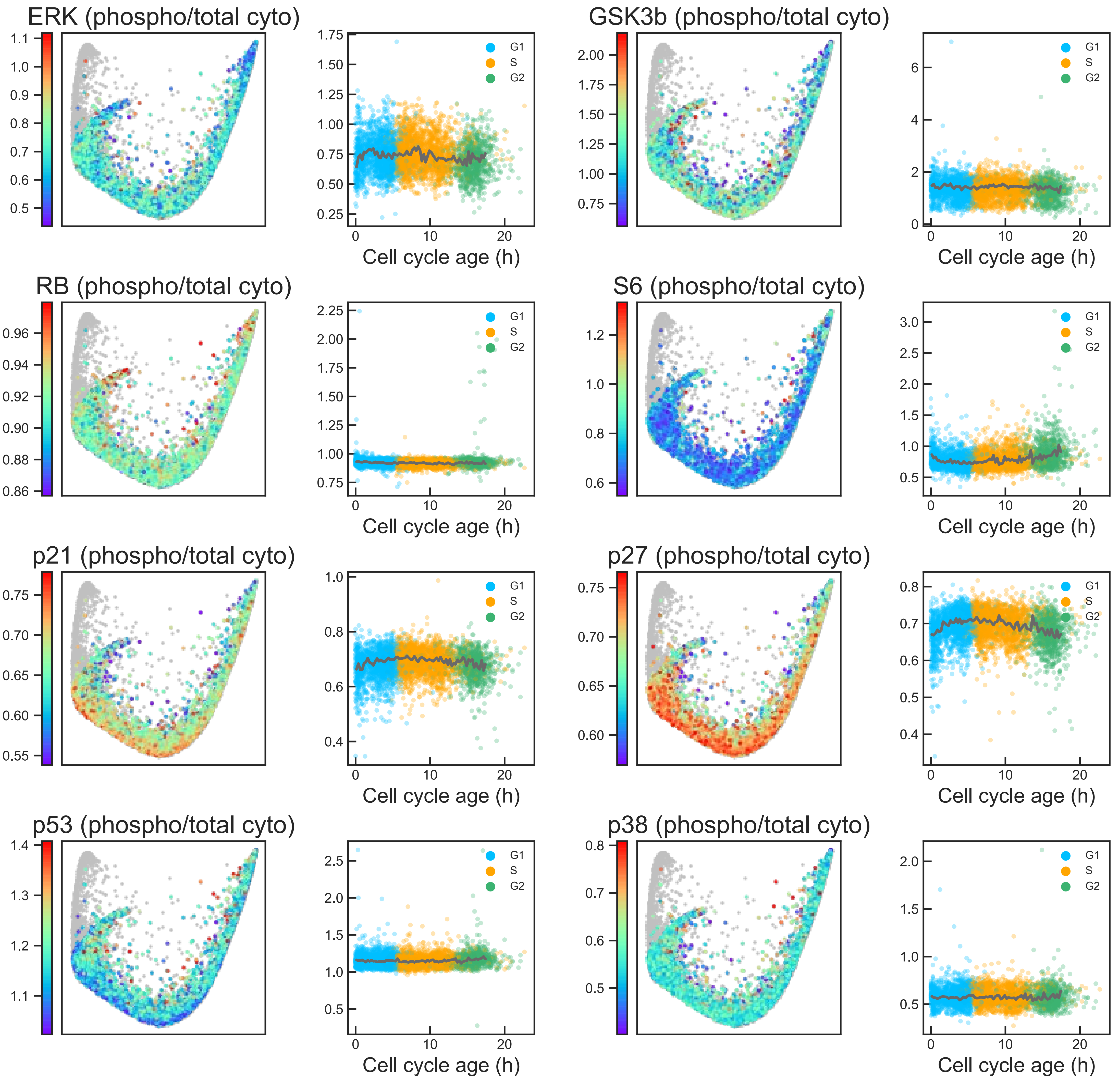

### Aux Fig. S1 - Proliferative trajectory - Page 34

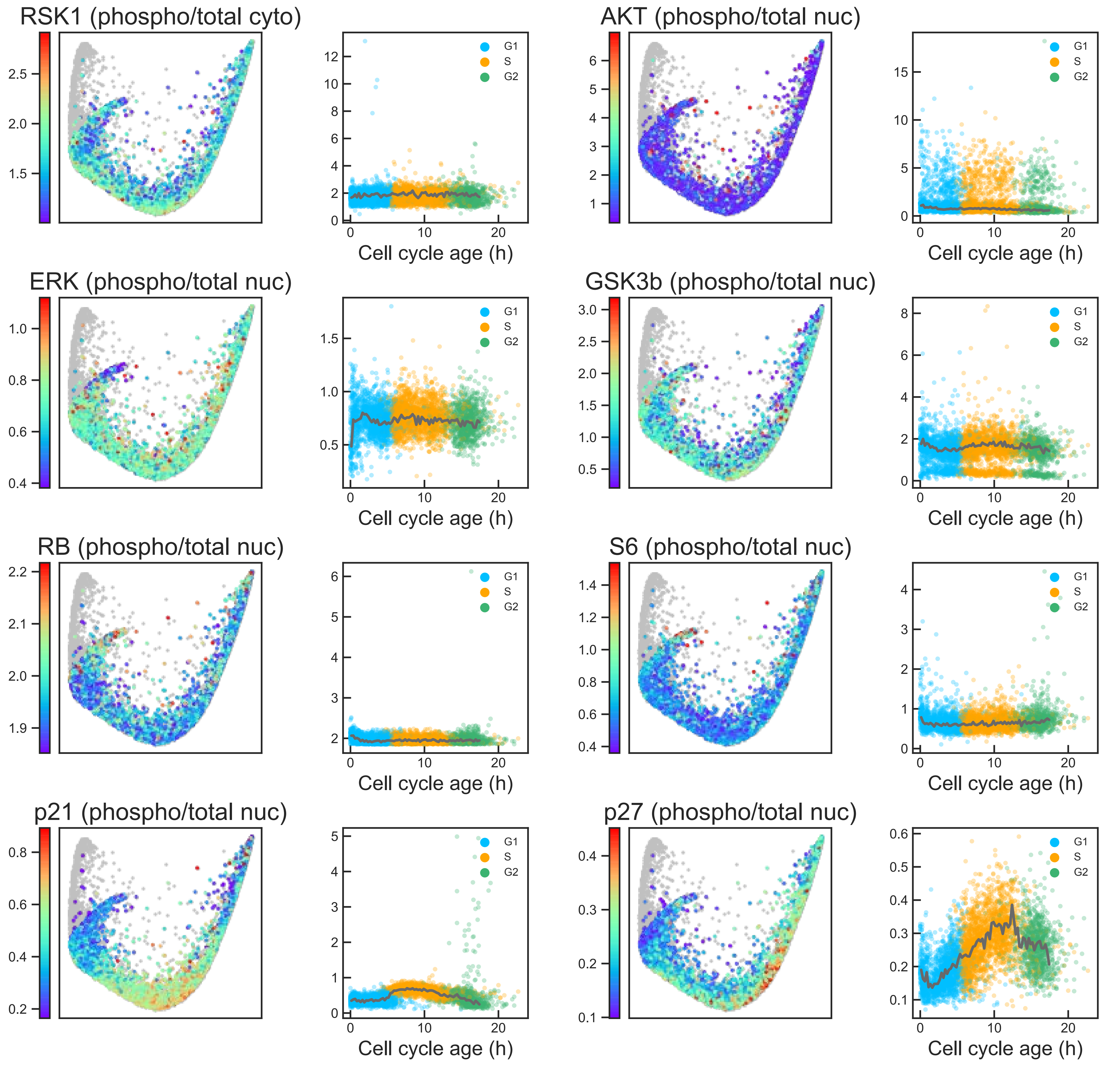

#### Aux Fig. S1 - Proliferative trajectory - Page 35

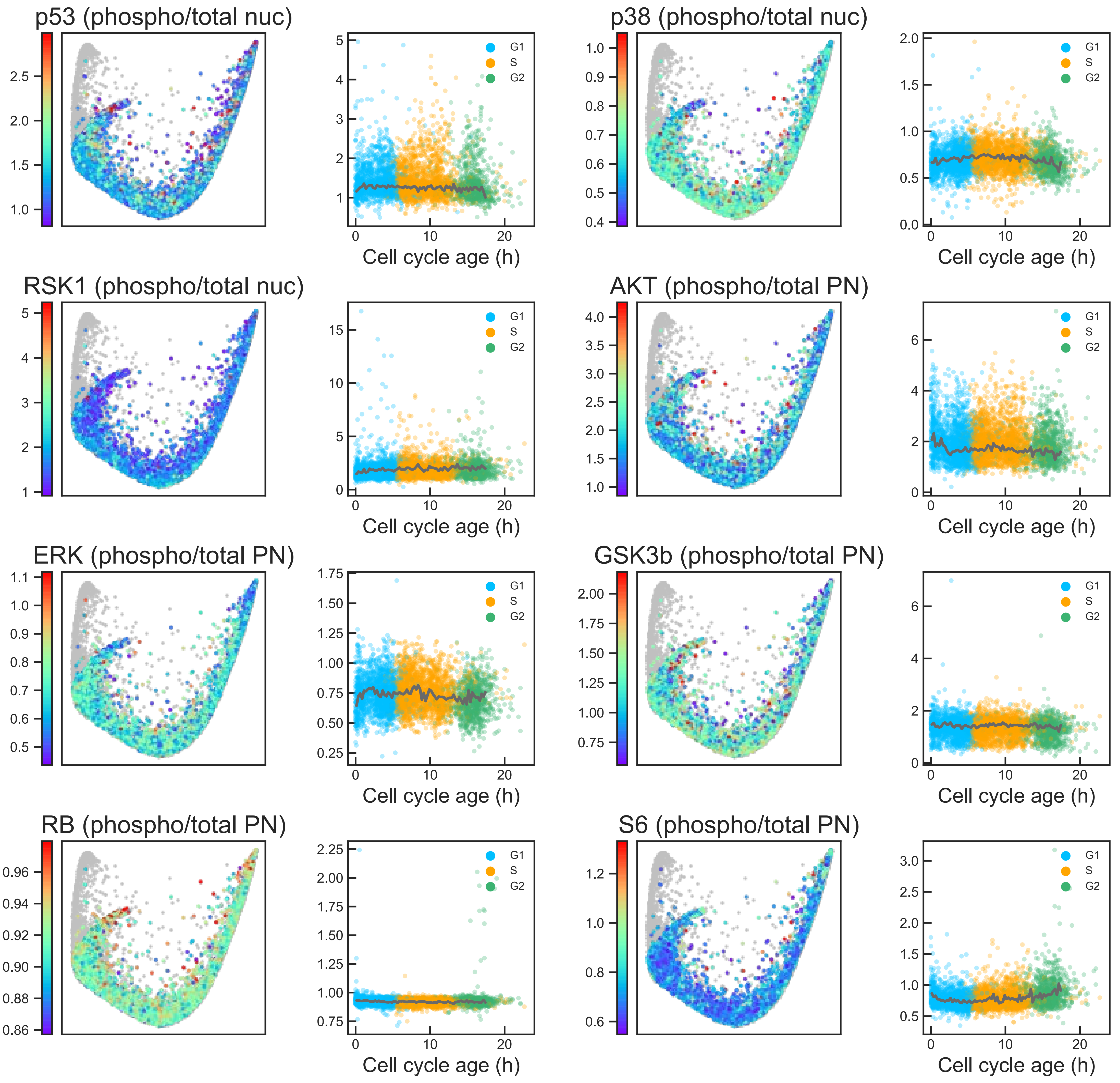

### Aux Fig. S1 - Proliferative trajectory - Page 36

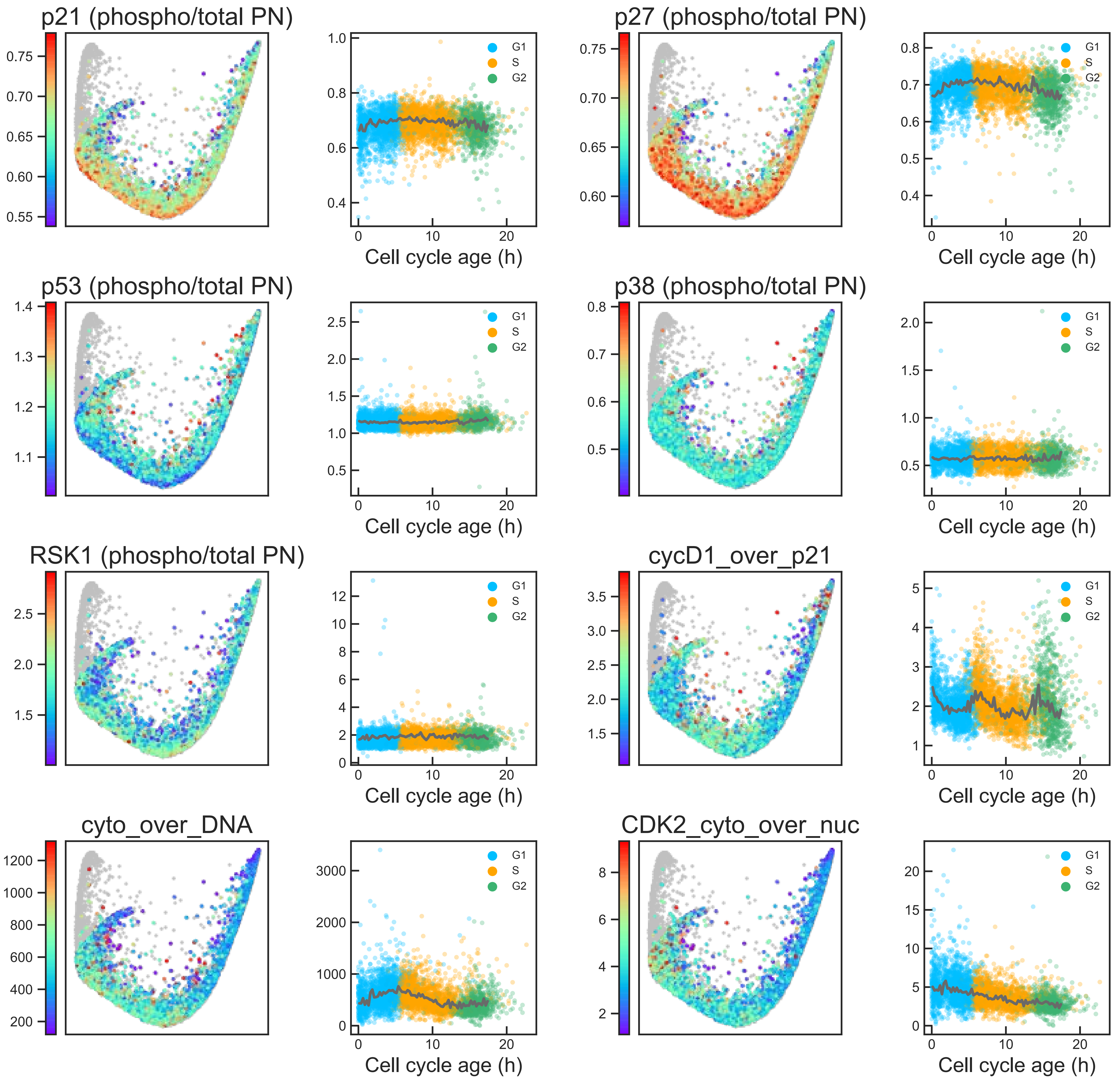
