## Supplementary material for "The structure of the human cell cycle": Auxiliary fig S2. Effector dynamics along the arrest trajectory

**Auxiliary Figure S2. Effector dynamics along the arrest trajectory.** Features are mapped onto the arrest trajectory of the cell cycle structure (left panels) and plotted against the duration of arrest (right panels). Population medians in time courses indicated by solid grey lines. Non-arrested cells (phospho/total RB $> 1.6$) are shown in grey on the structure and are excluded from time courses. Plasma membrane, PM; perinuclear, PN.

### Aux Fig. S2 - Arrest trajectory - Page 1

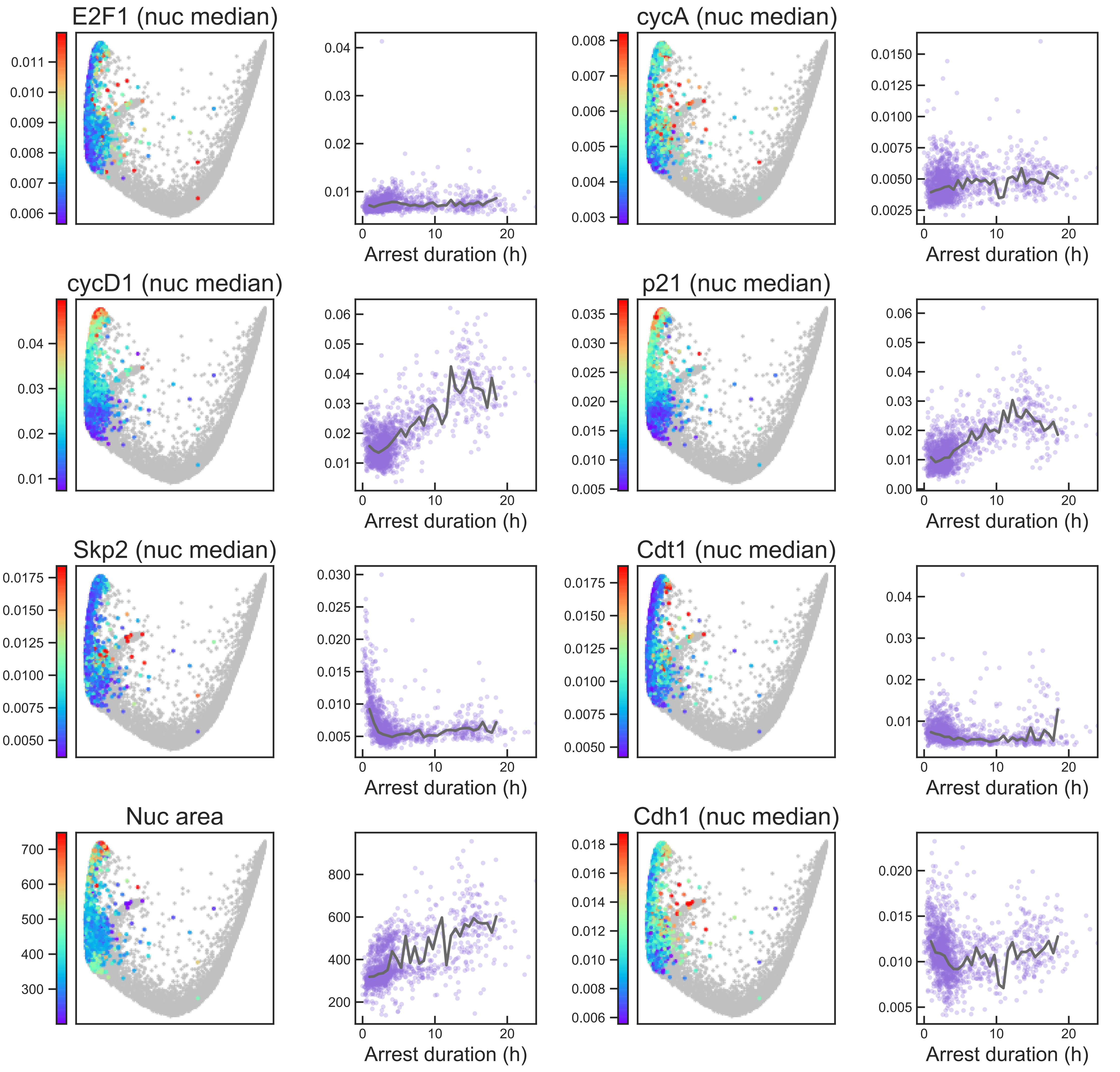

### Aux Fig. S2 - Arrest trajectory - Page 2

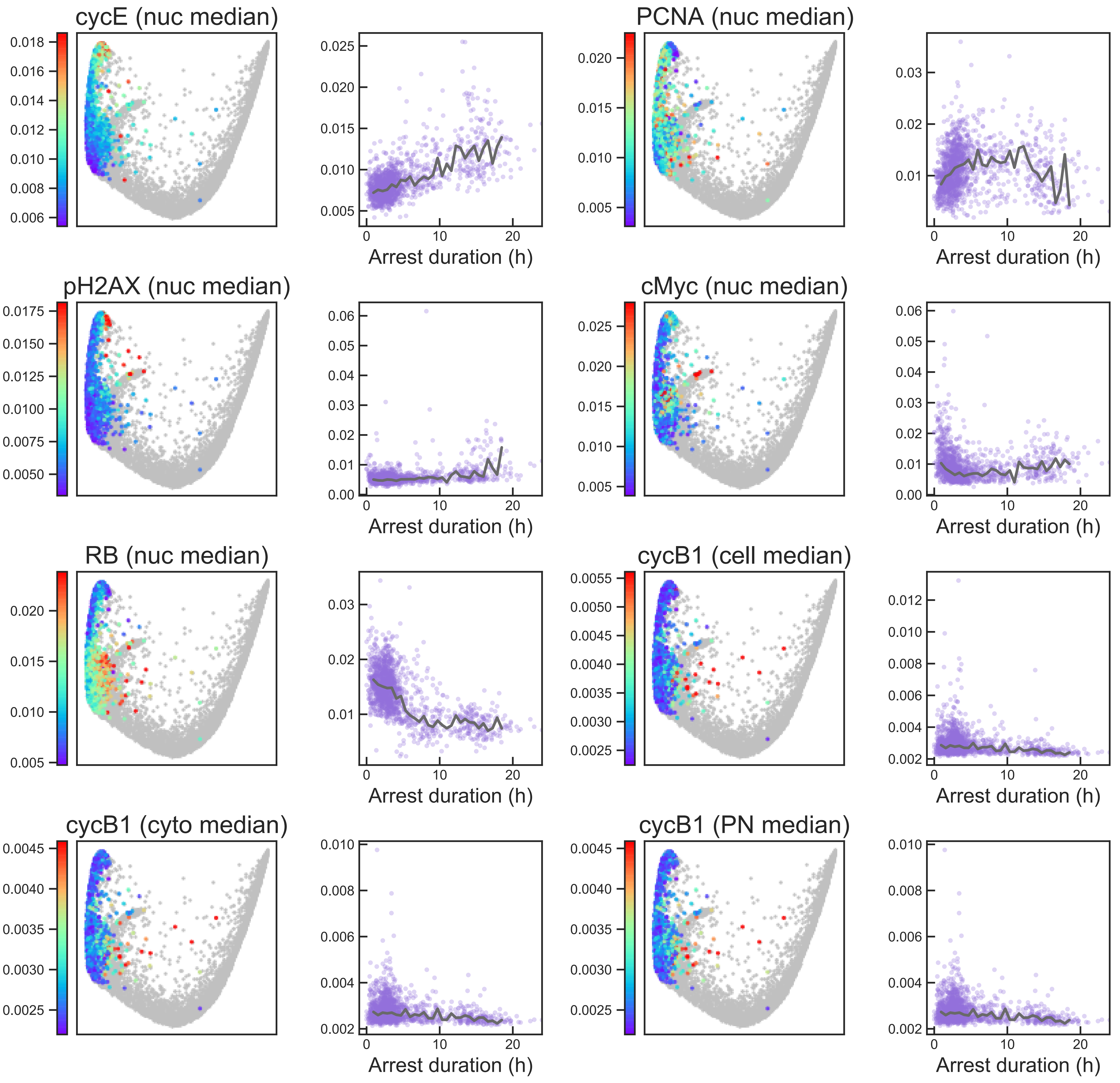

### Aux Fig. S2 - Arrest trajectory - Page 3

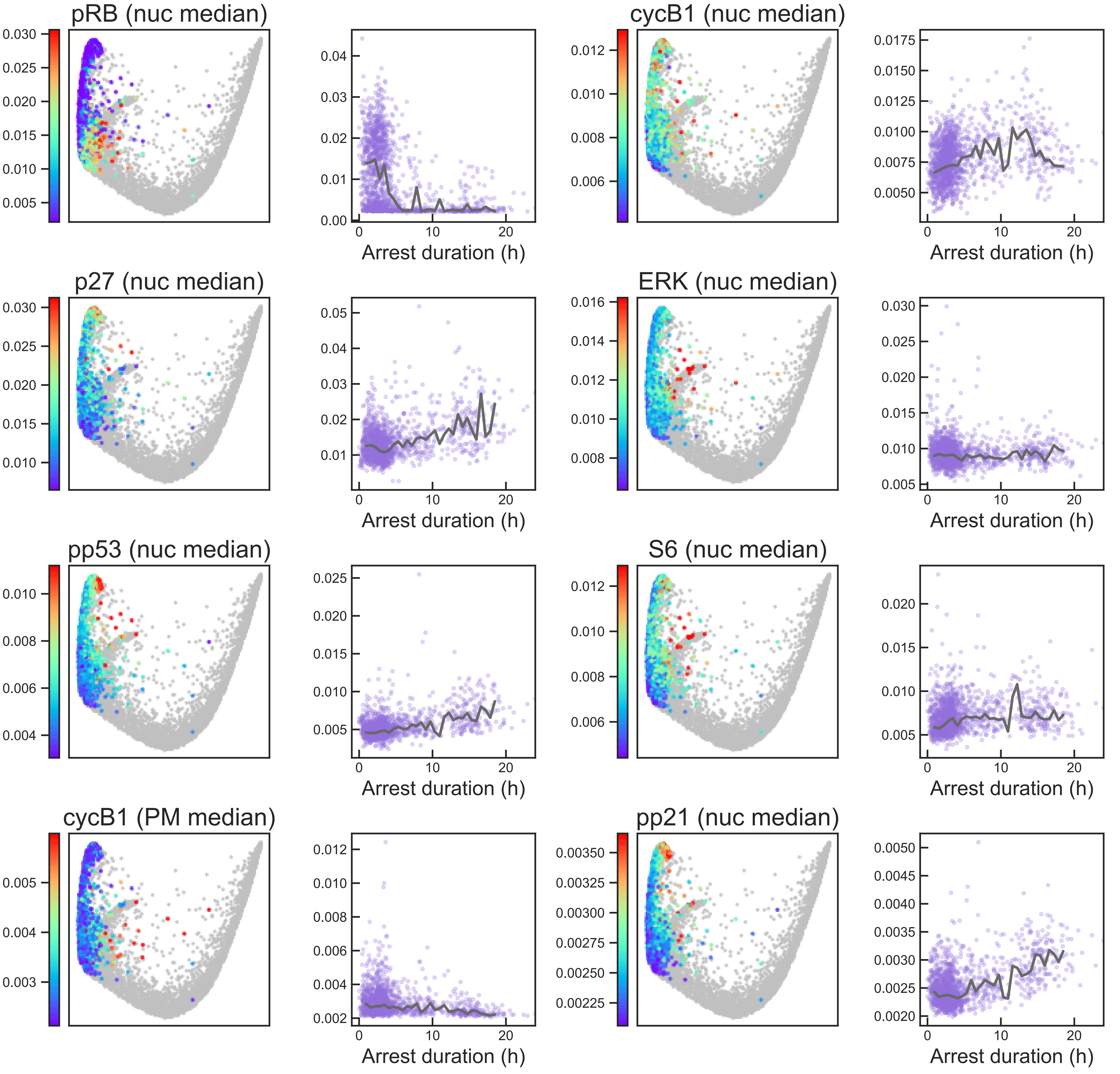

#### Aux Fig. S2 - Arrest trajectory - Page 4

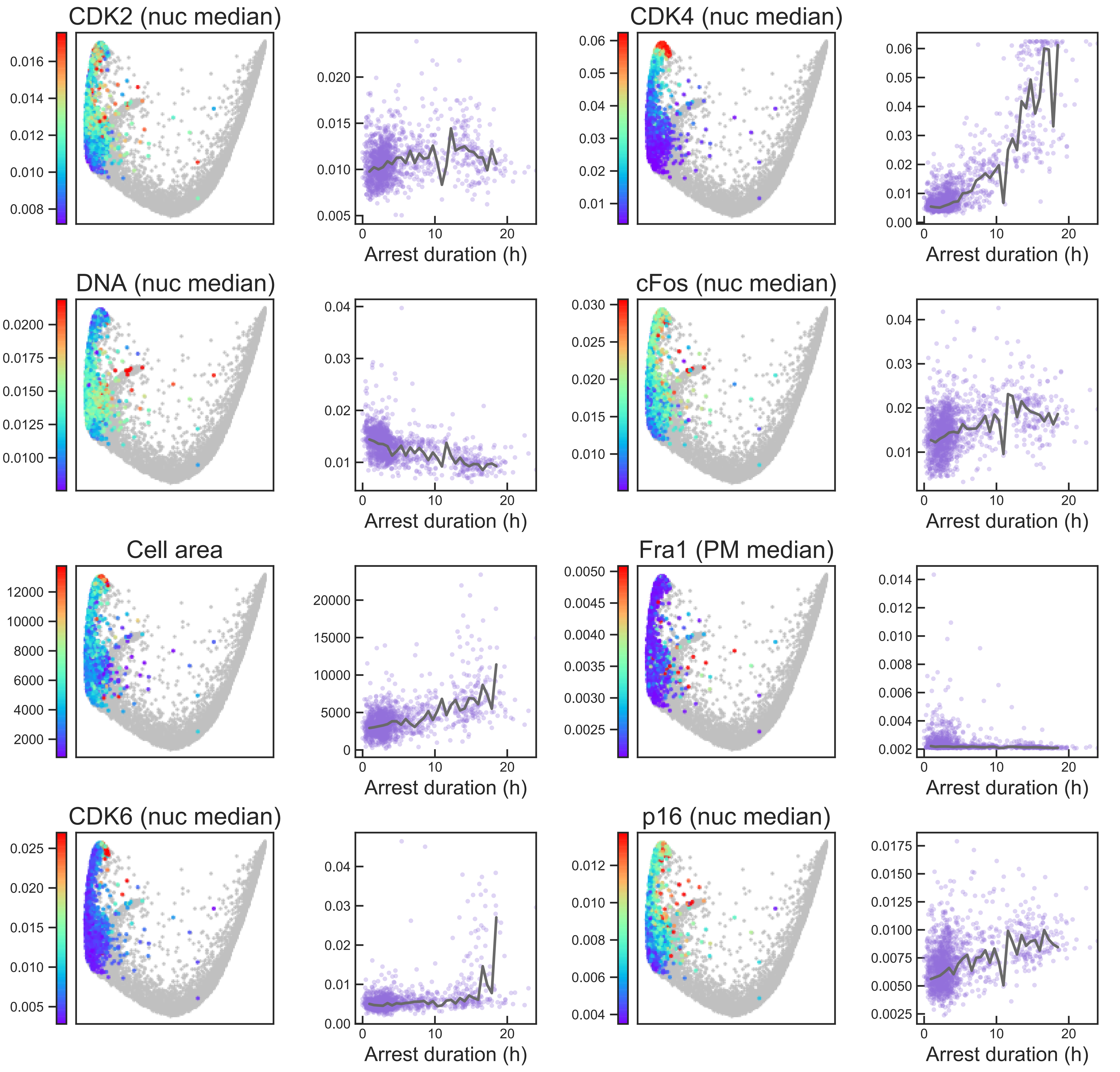

#### Aux Fig. S2 - Arrest trajectory - Page 5

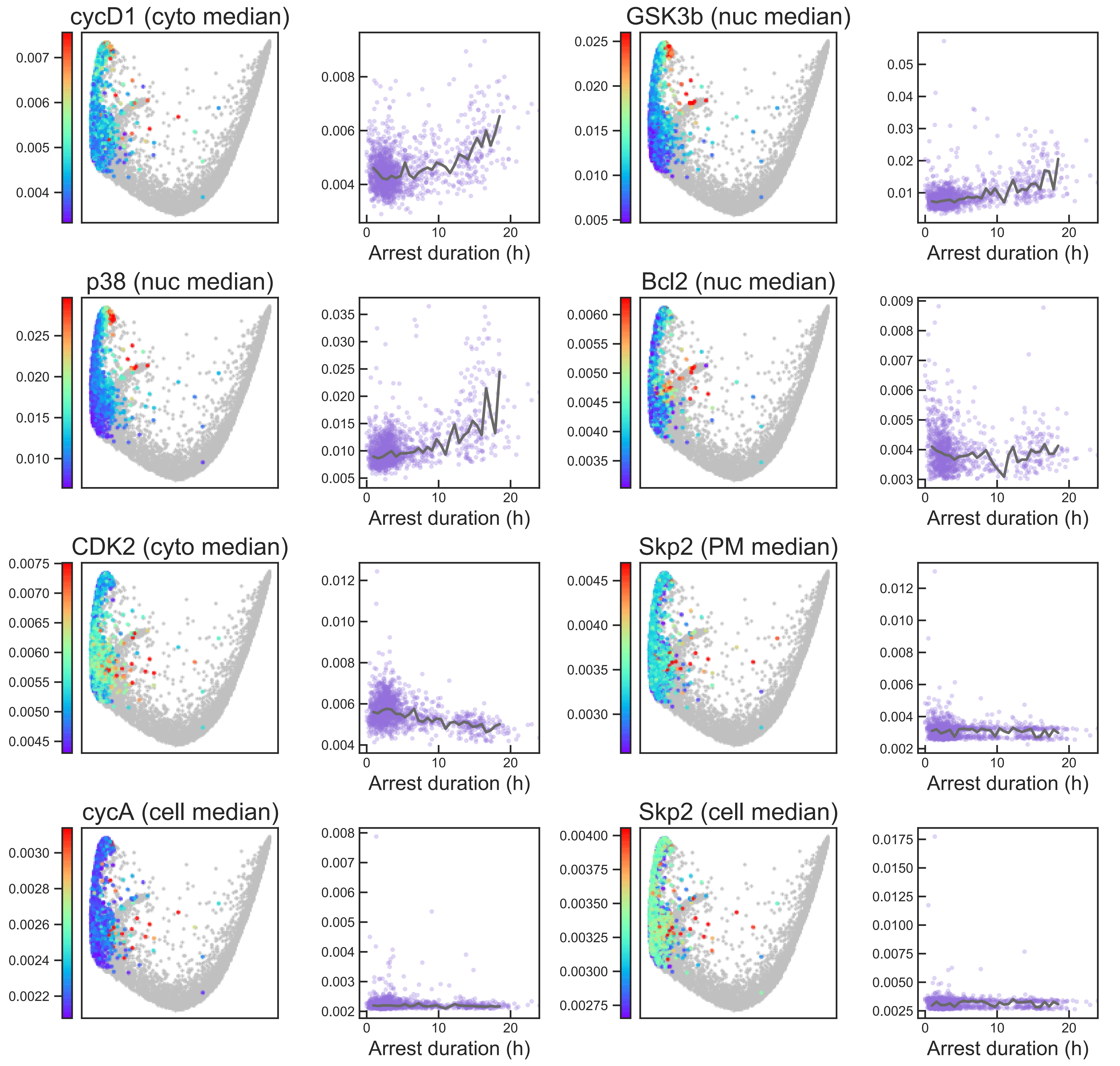

#### Aux Fig. S2 - Arrest trajectory - Page 6

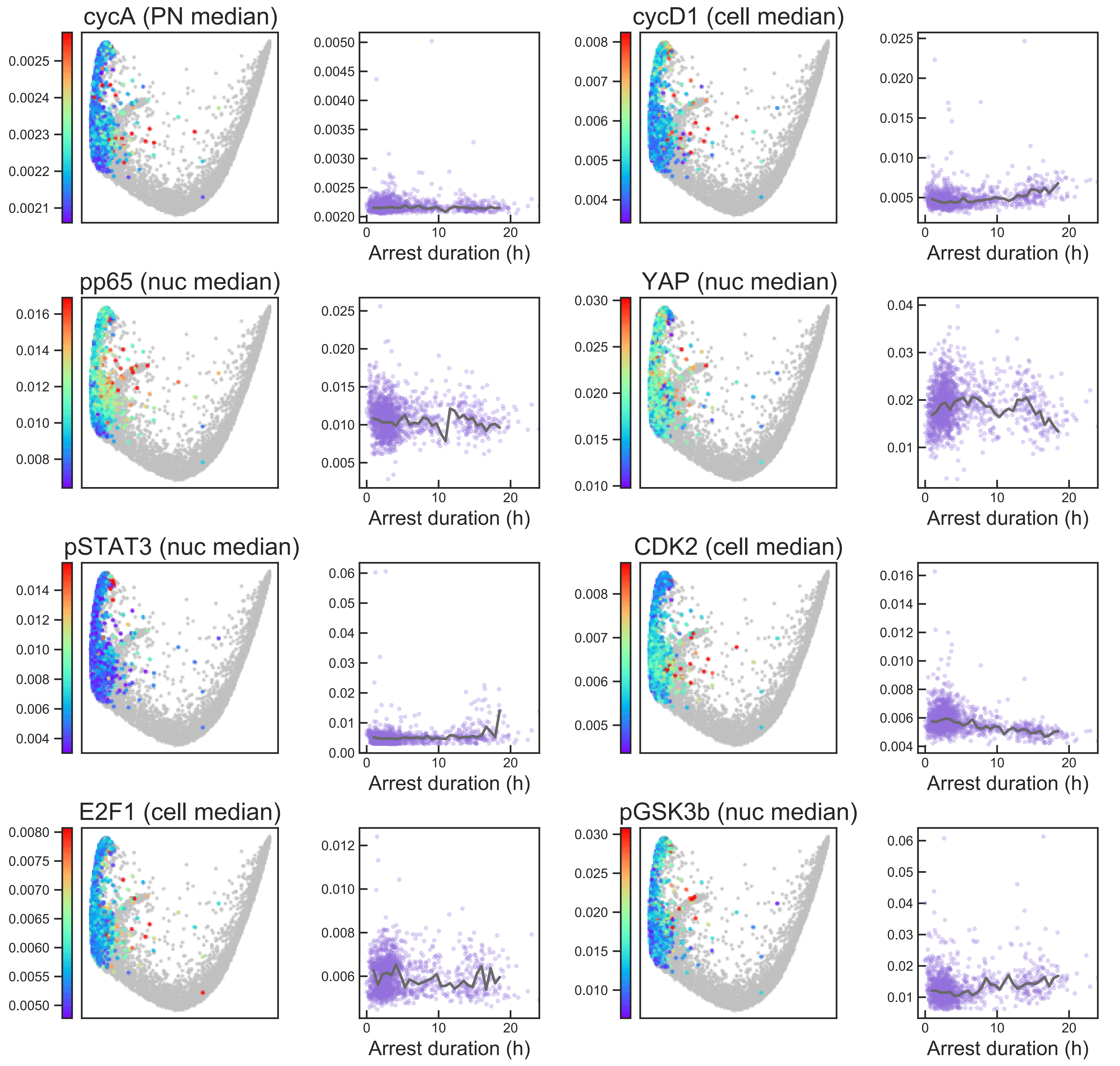

### Aux Fig. S2 - Arrest trajectory - Page 7

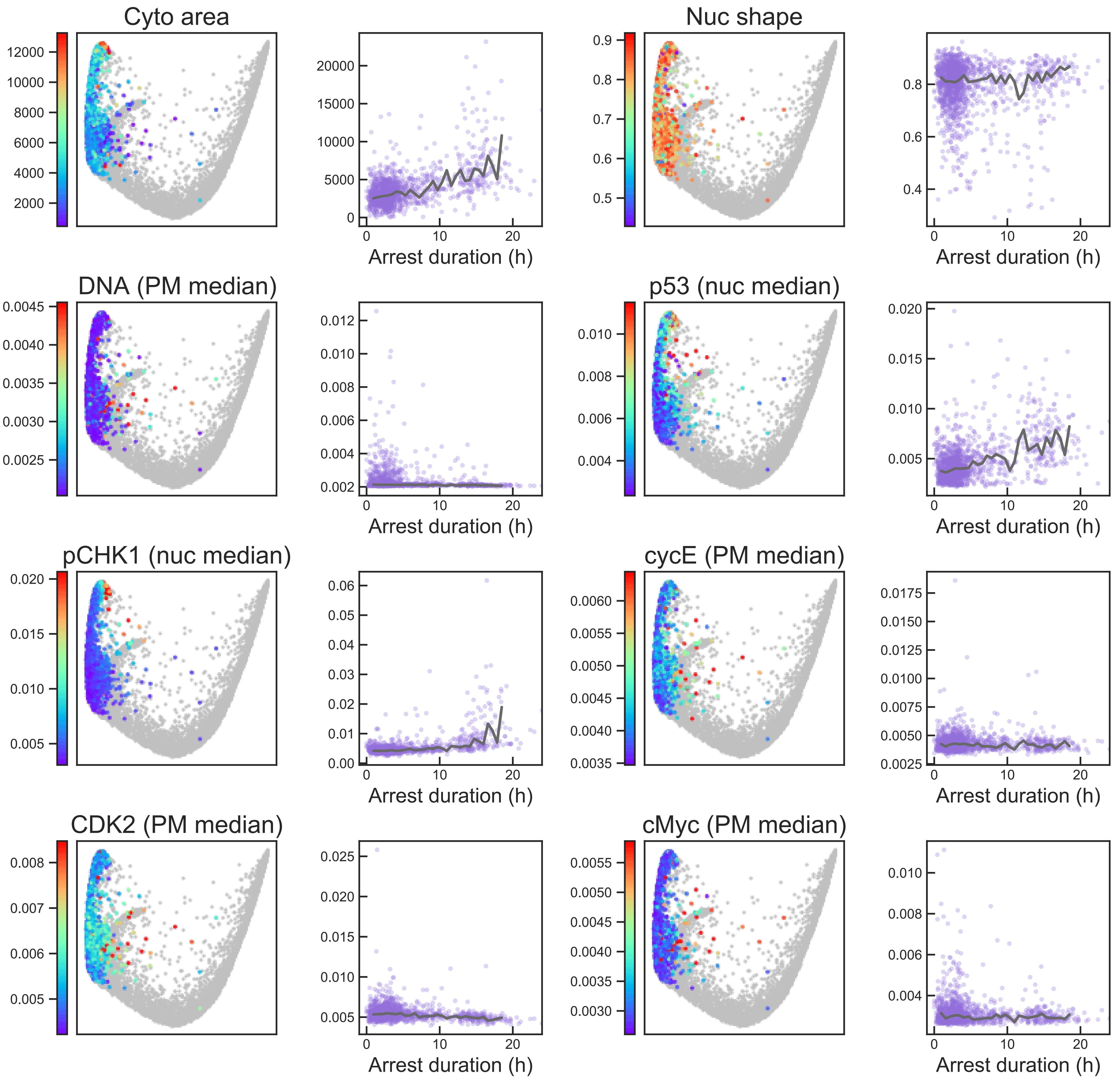

### Aux Fig. S2 - Arrest trajectory - Page 8

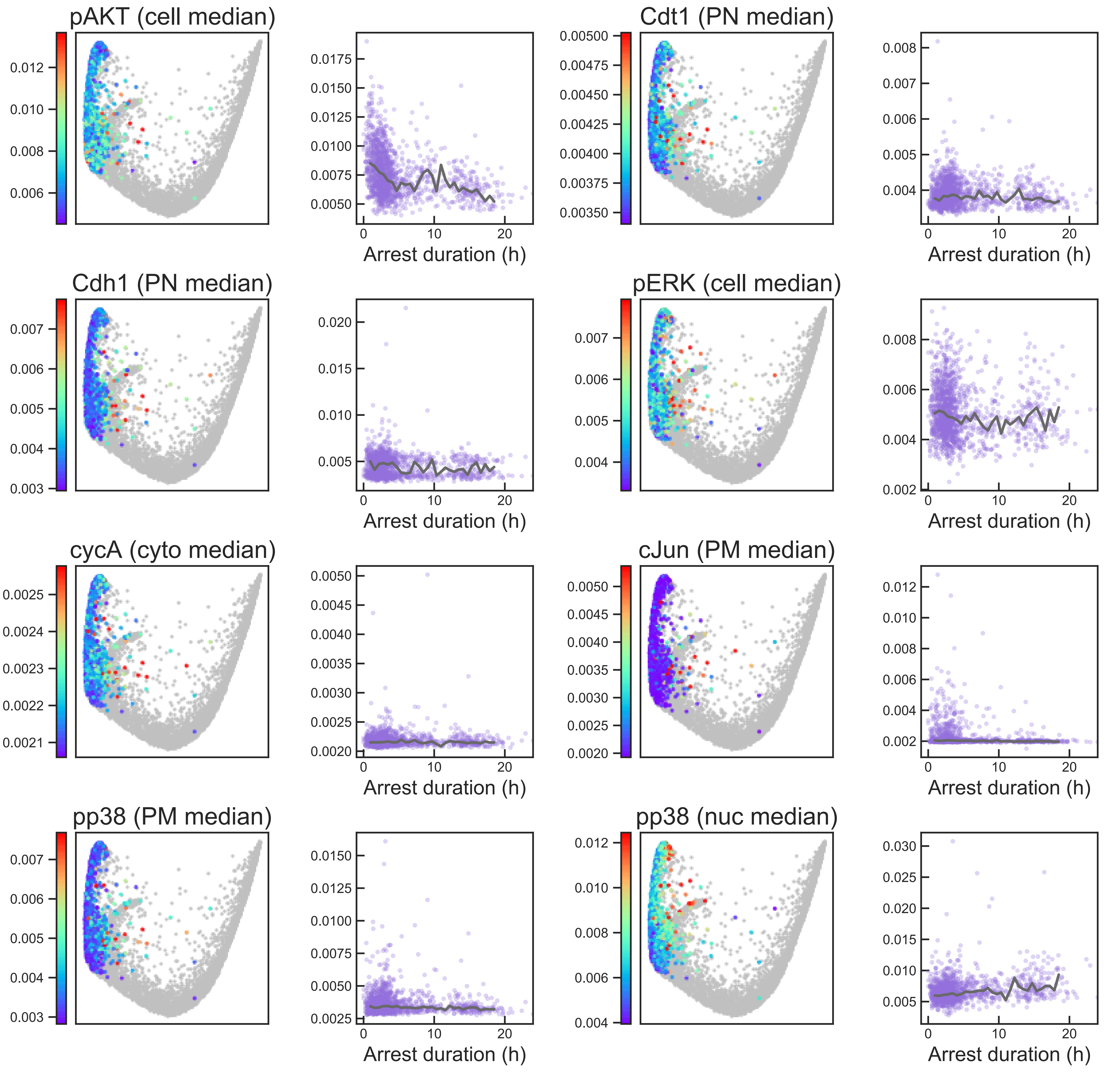

### Aux Fig. S2 - Arrest trajectory - Page 9

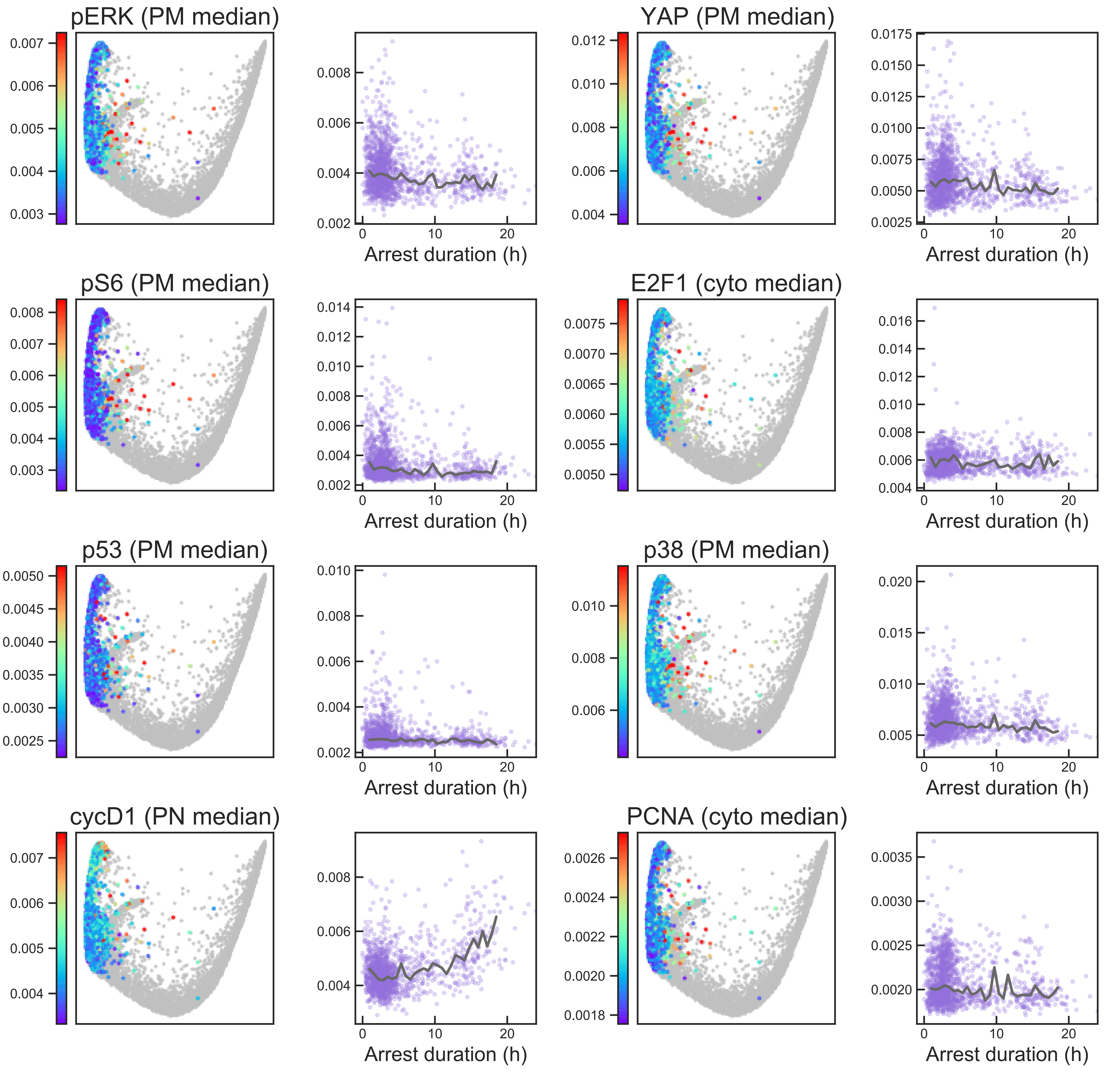

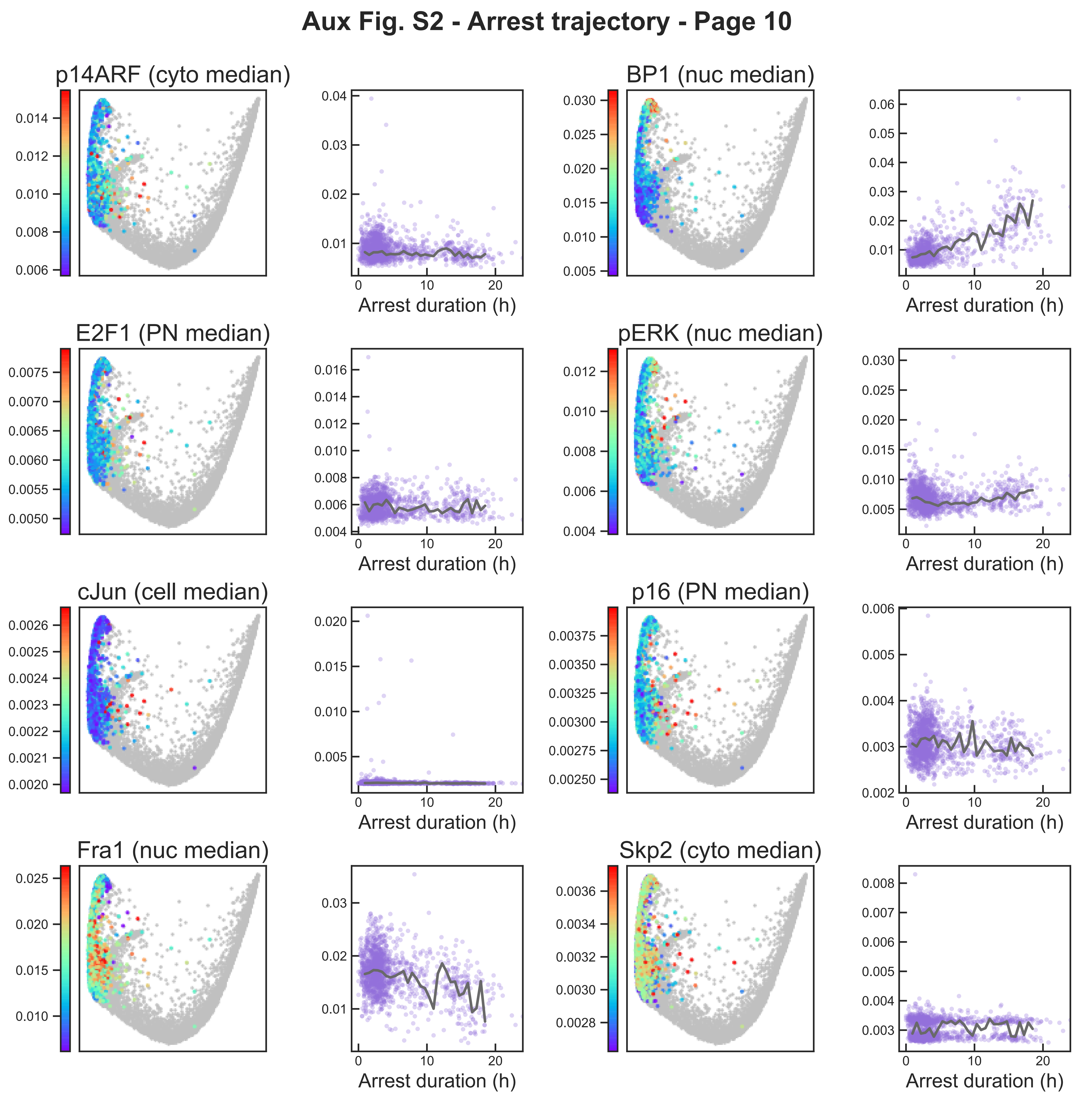
Aux Fig. S2 - Arrest trajectory - Page 10
p14ARF (cyto median)
BP1 (nuc median)
E2F1 (PN median)
pERK (nuc median)
cJun (cell median)
p16 (PN median)
Fra1 (nuc median)
Skp2 (cyto median)
Arrest duration (h)

**Aux Fig. S2 - Arrest trajectory - Page 11**

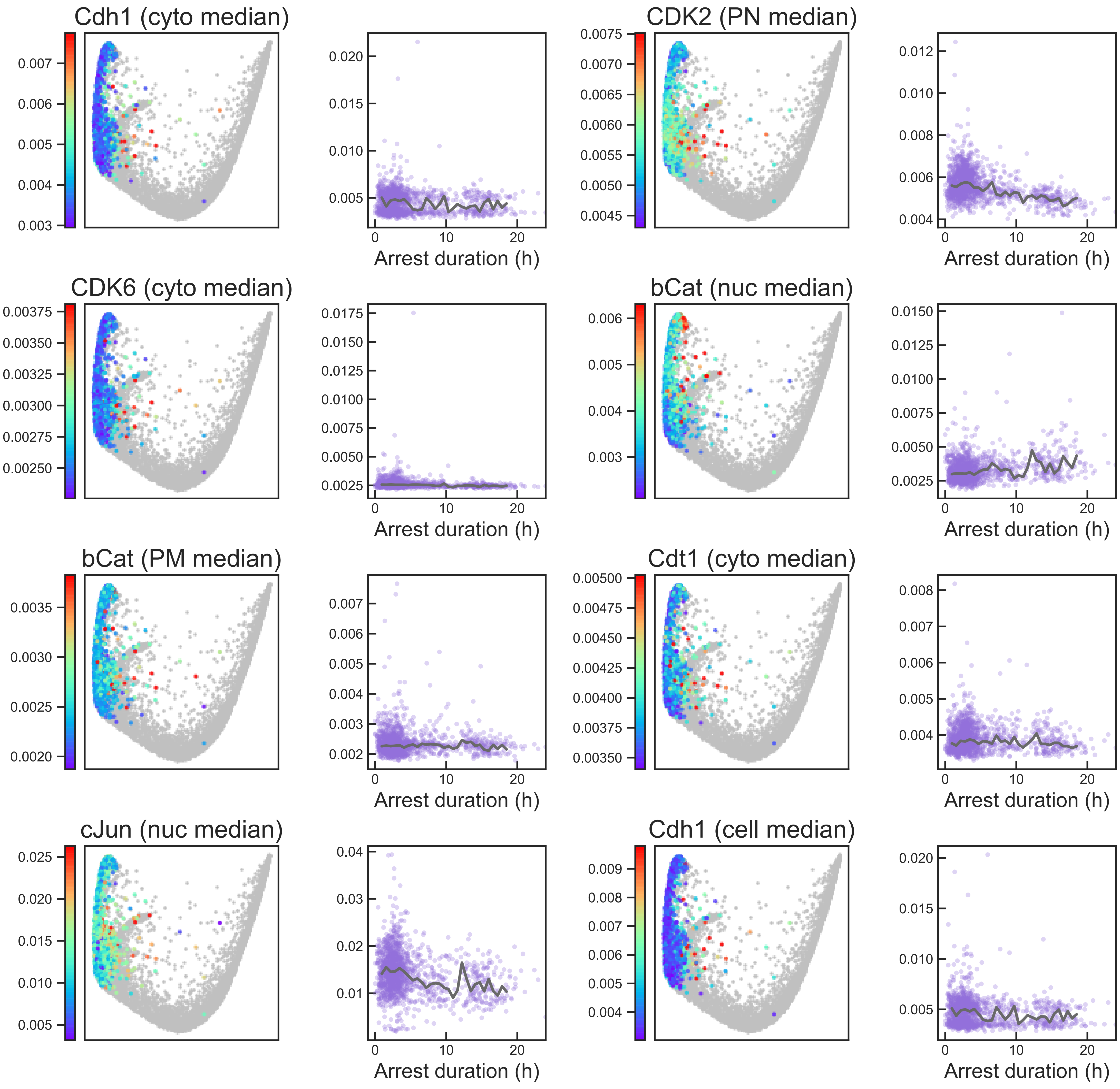

#### Aux Fig. S2 - Arrest trajectory - Page 12

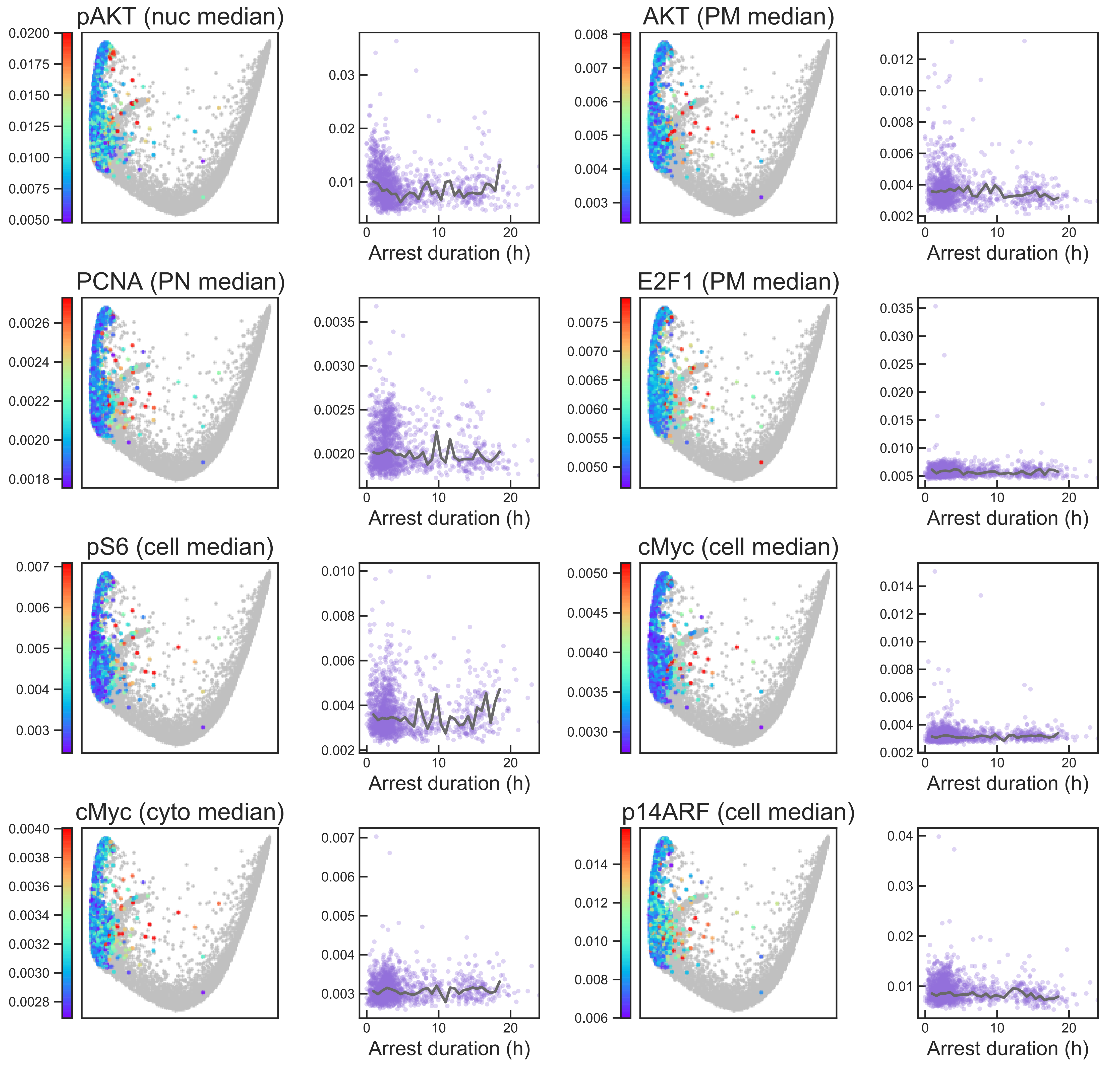

### Aux Fig. S2 - Arrest trajectory - Page 13

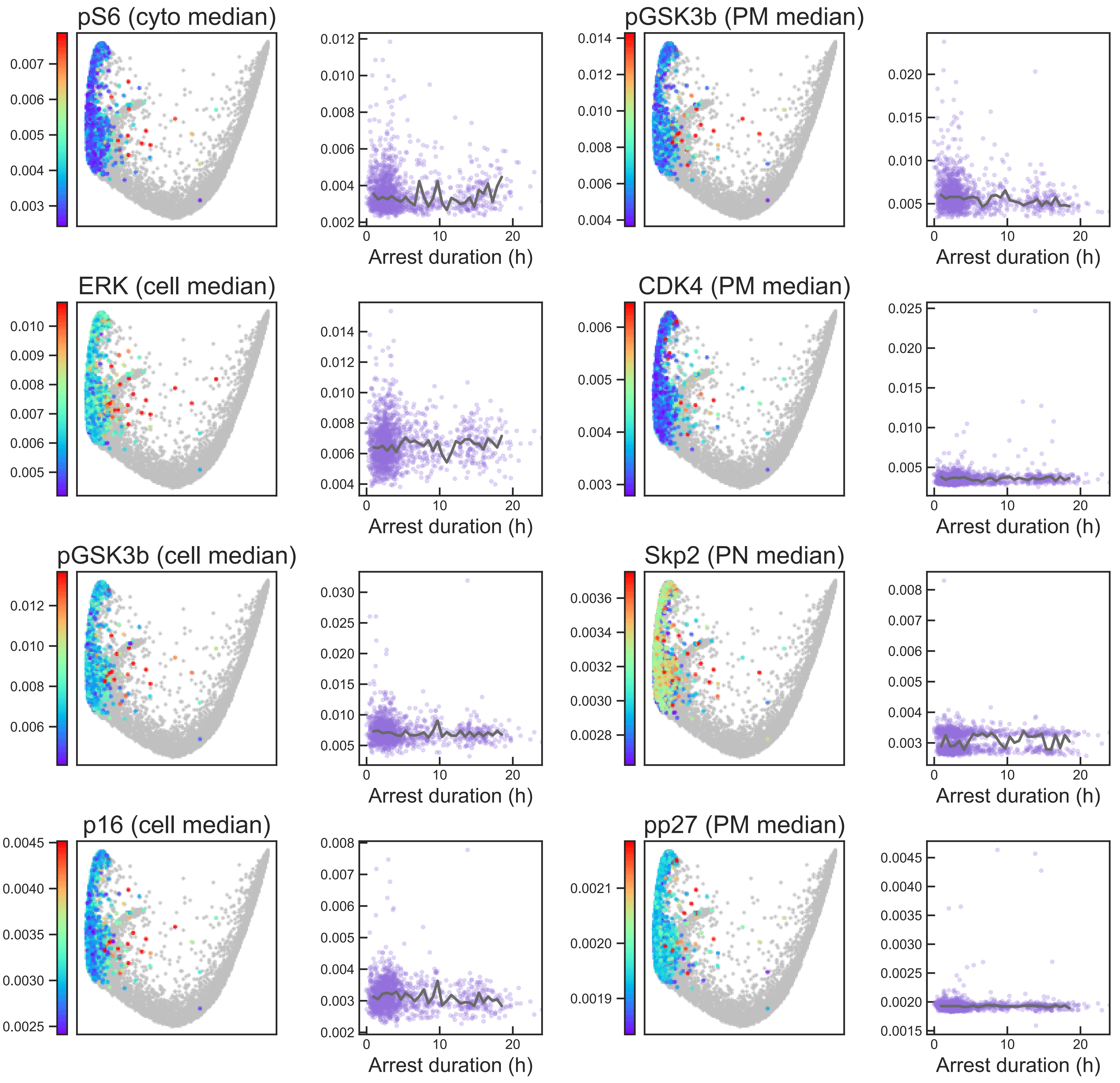

#### Aux Fig. S2 - Arrest trajectory - Page 14

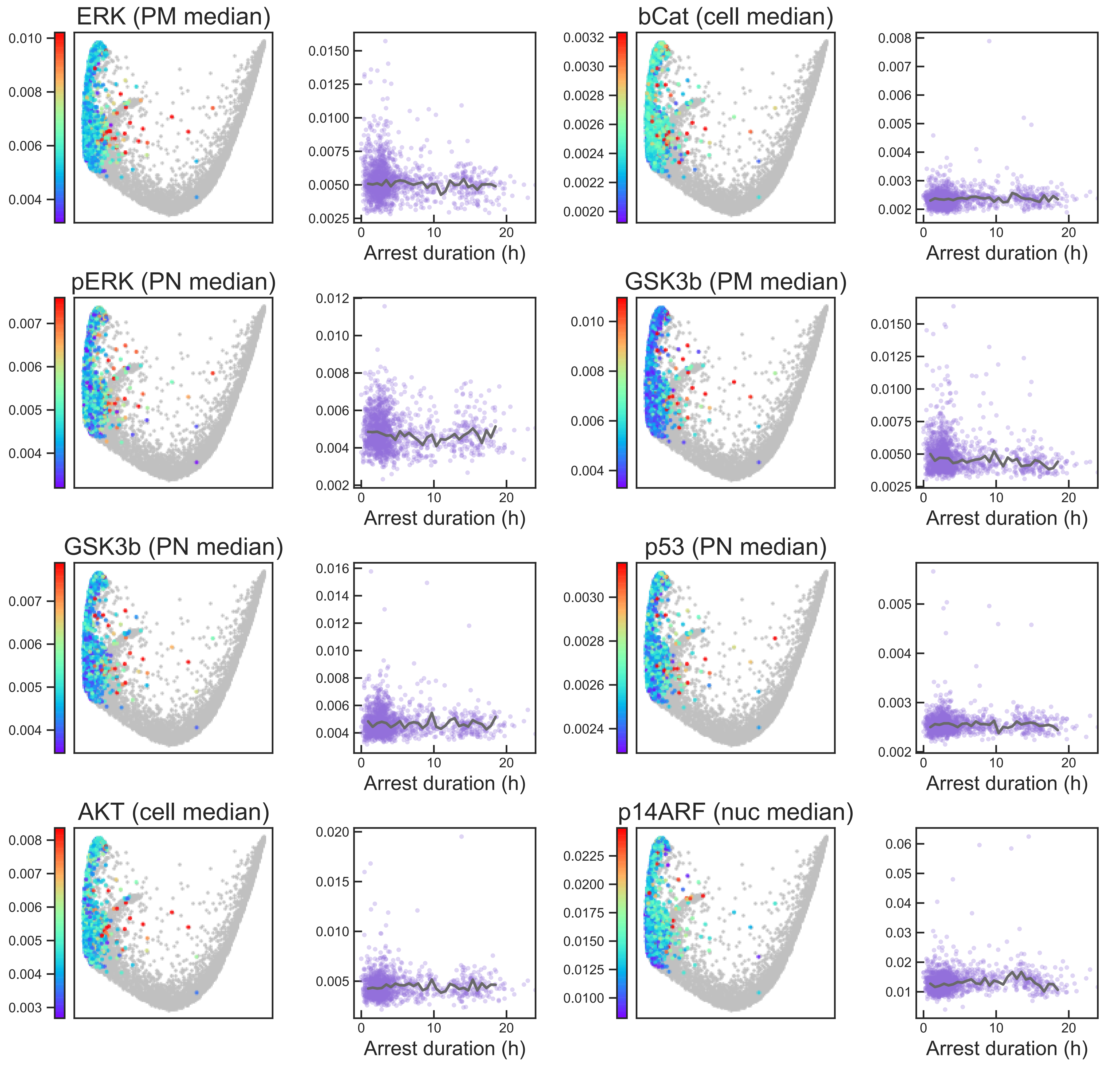

#### Aux Fig. S2 - Arrest trajectory - Page 15

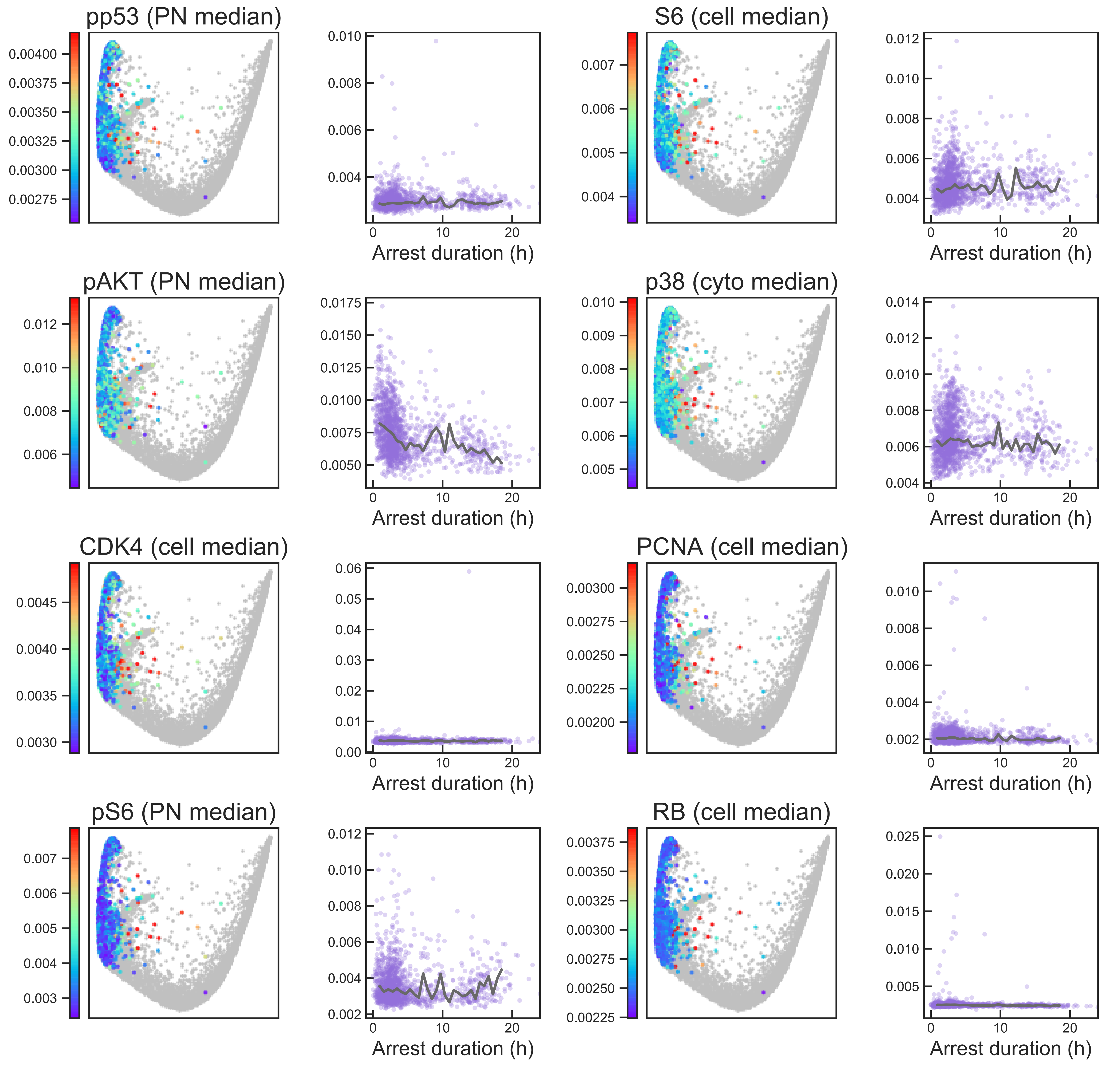

#### Aux Fig. S2 - Arrest trajectory - Page 16

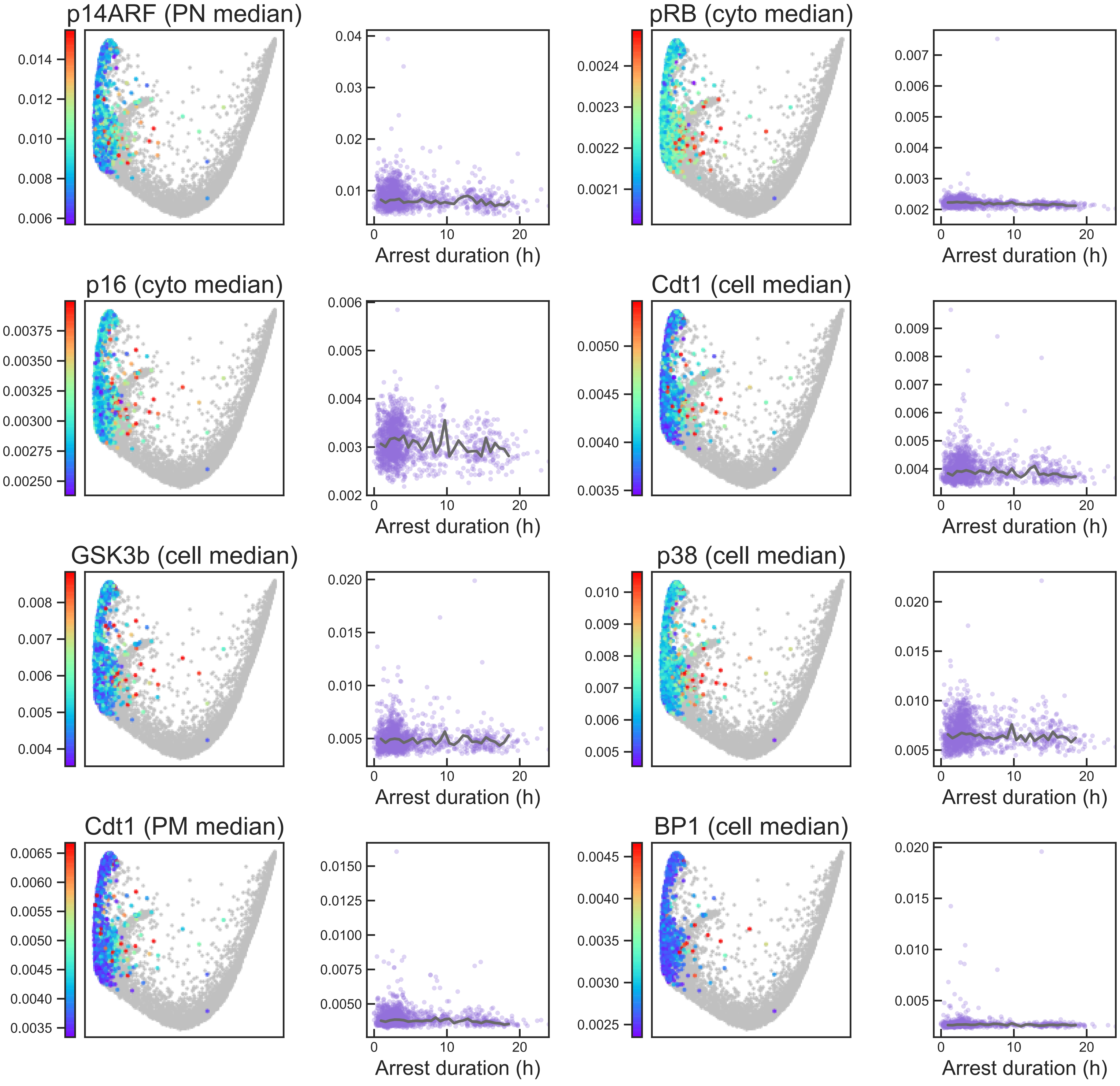

#### Aux Fig. S2 - Arrest trajectory - Page 17

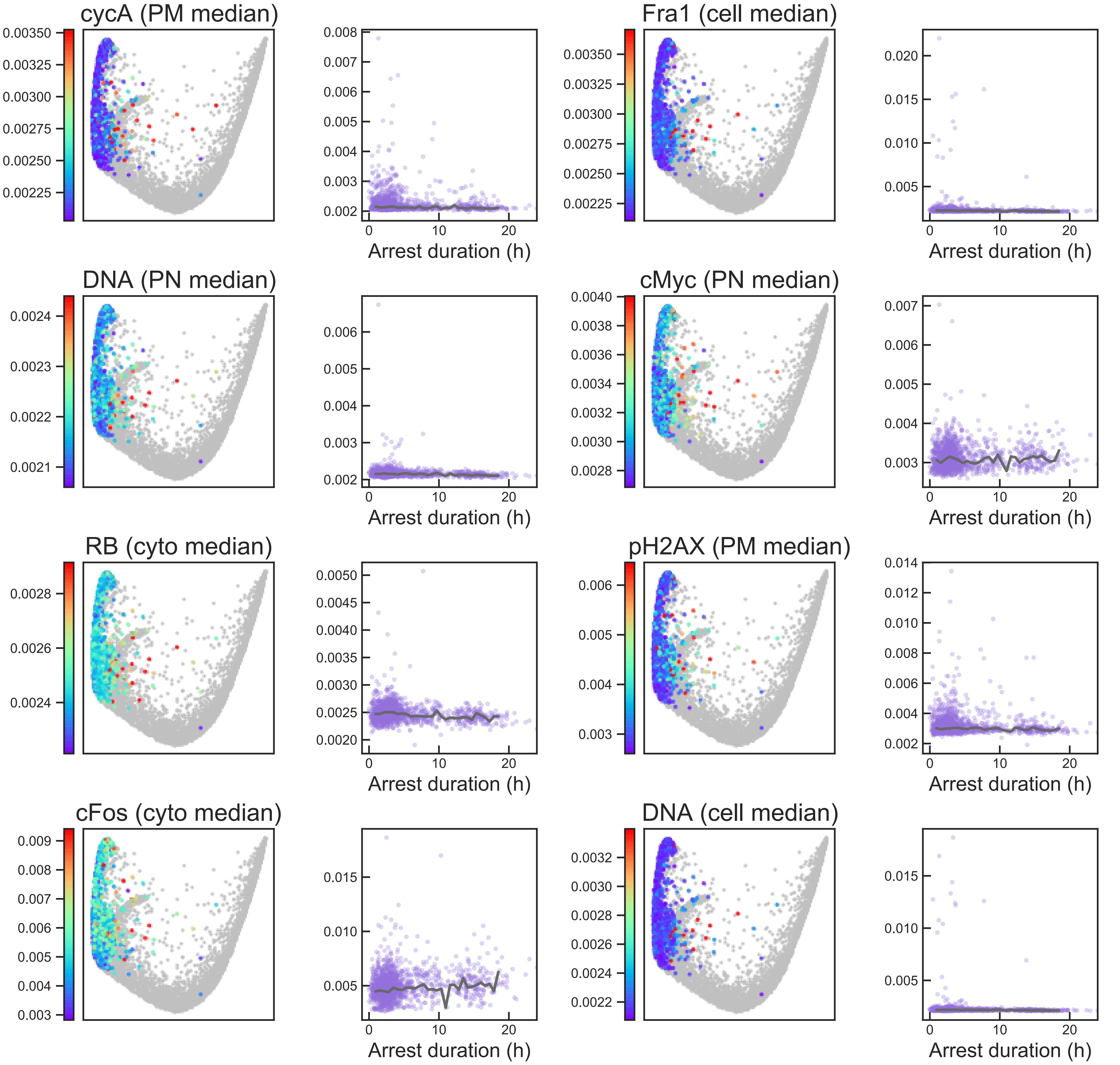

### Aux Fig. S2 - Arrest trajectory - Page 18

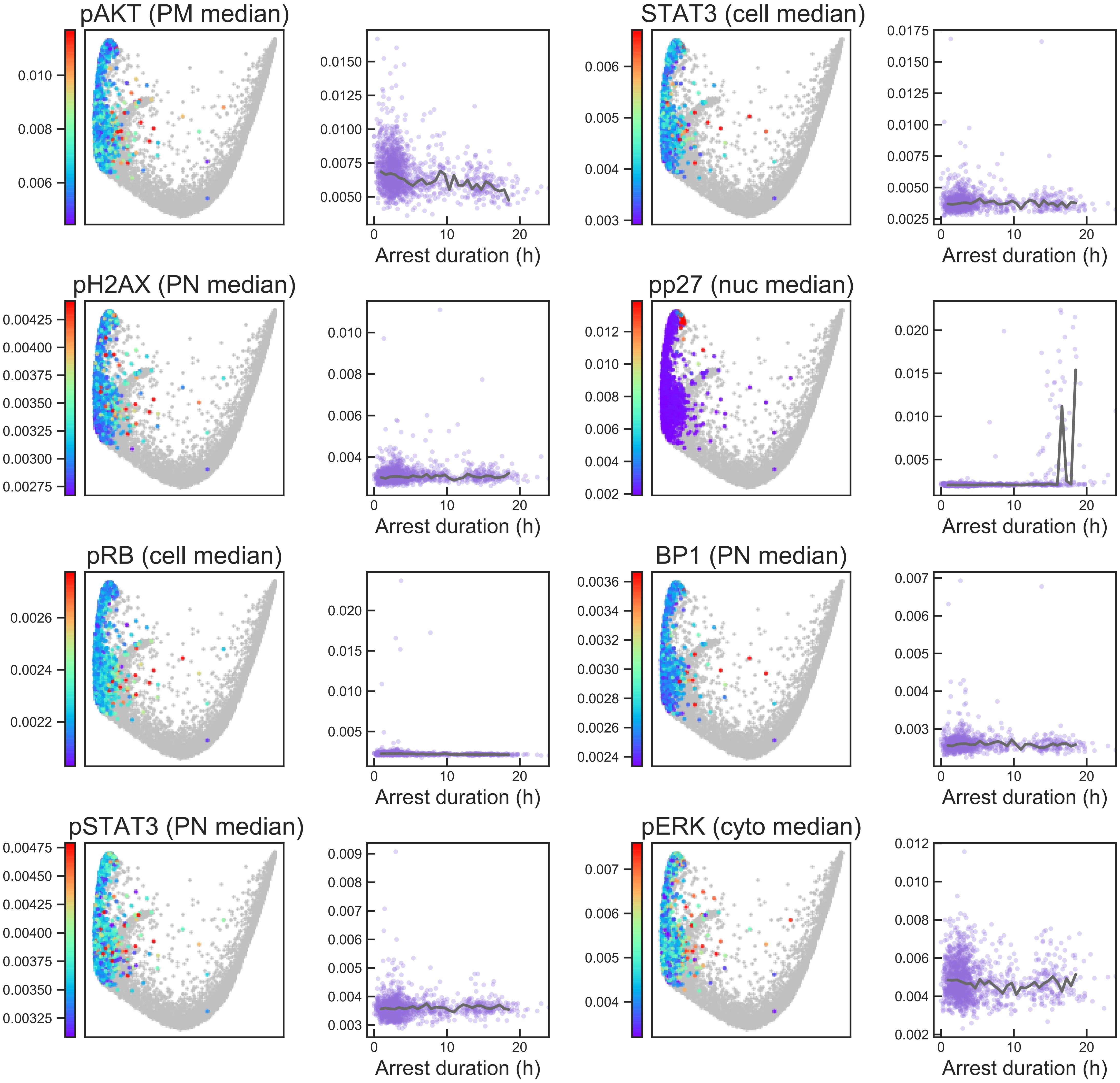

#### Aux Fig. S2 - Arrest trajectory - Page 19

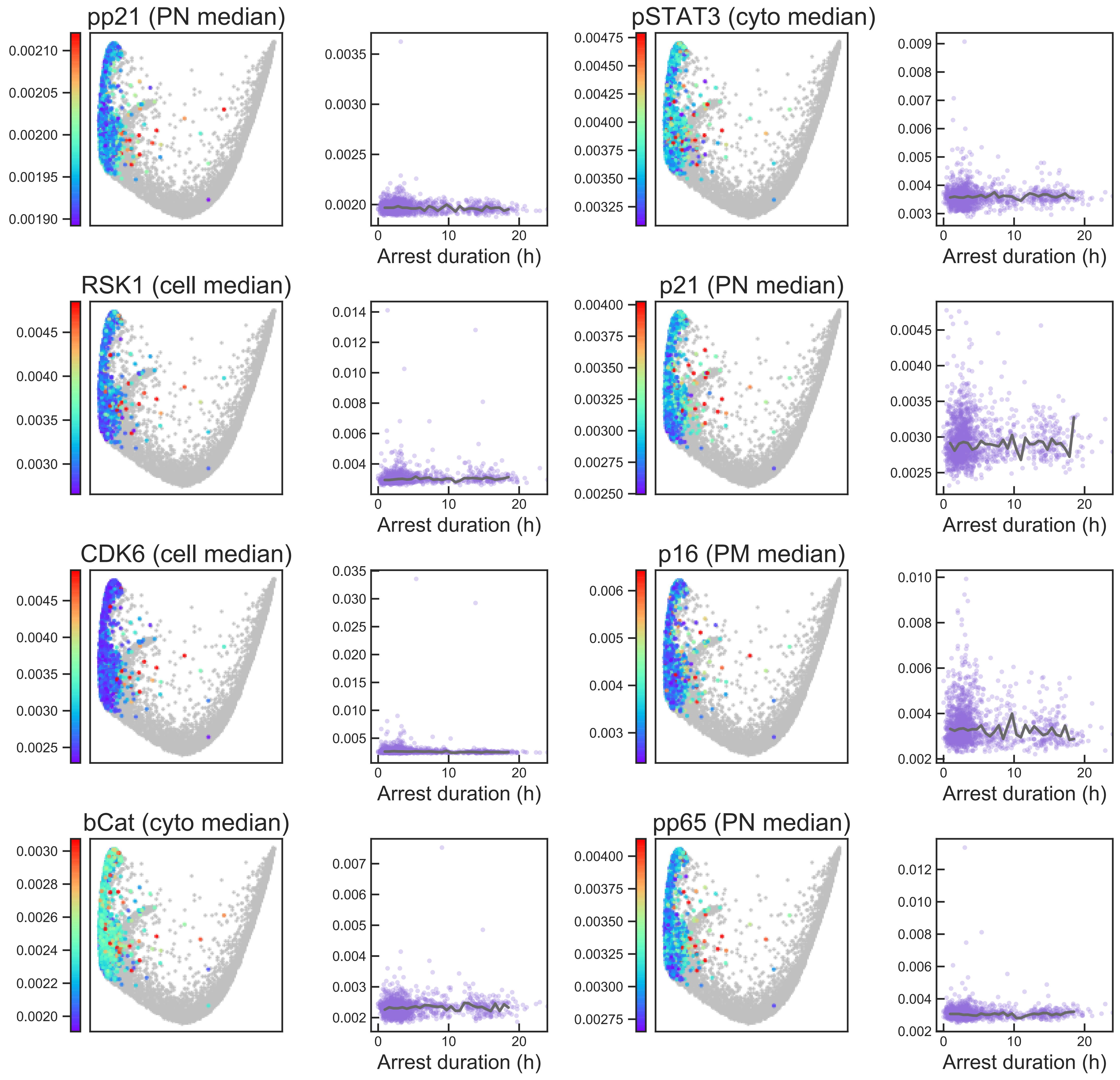

### Aux Fig. S2 - Arrest trajectory - Page 20

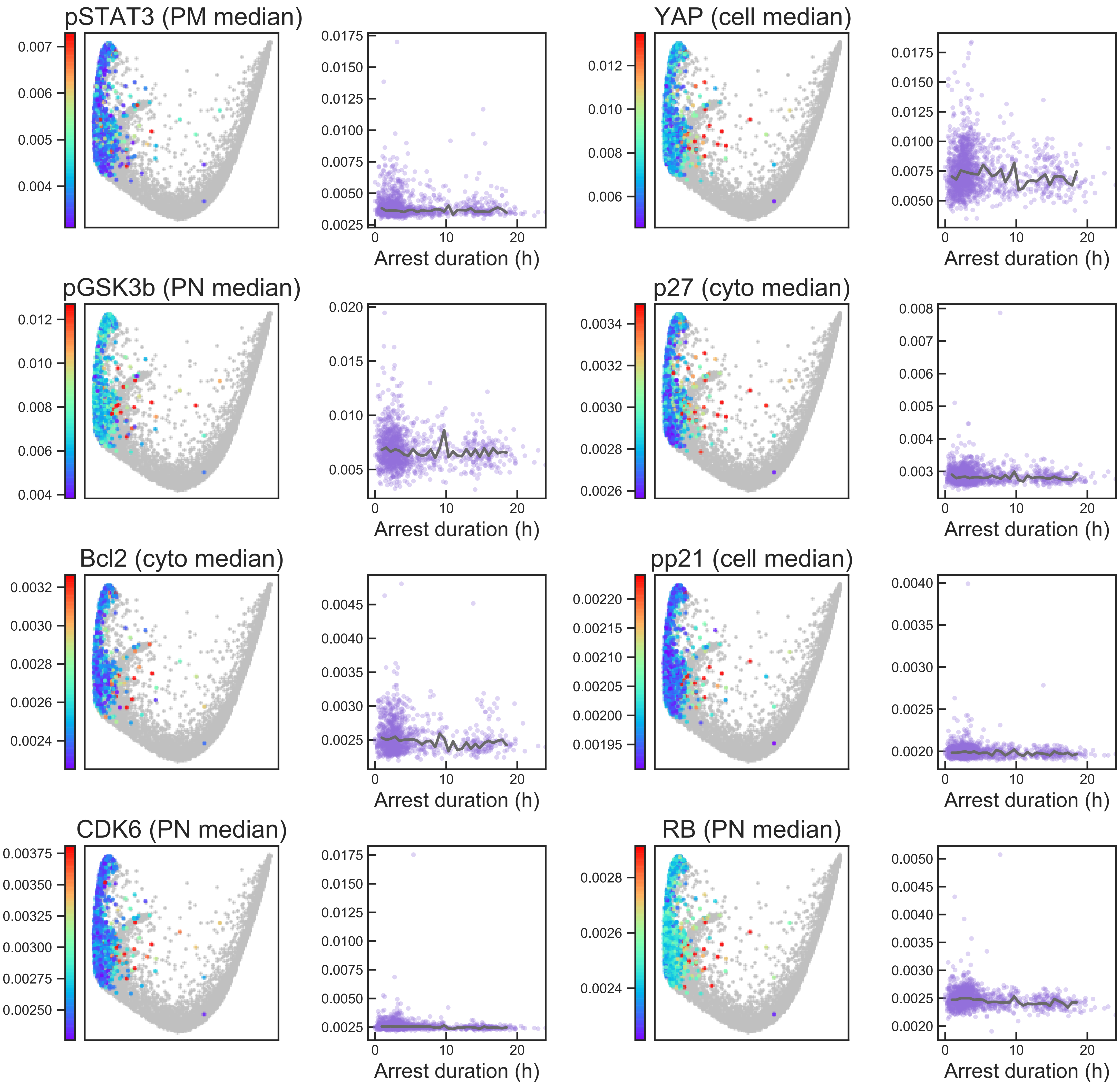

### Aux Fig. S2 - Arrest trajectory - Page 21

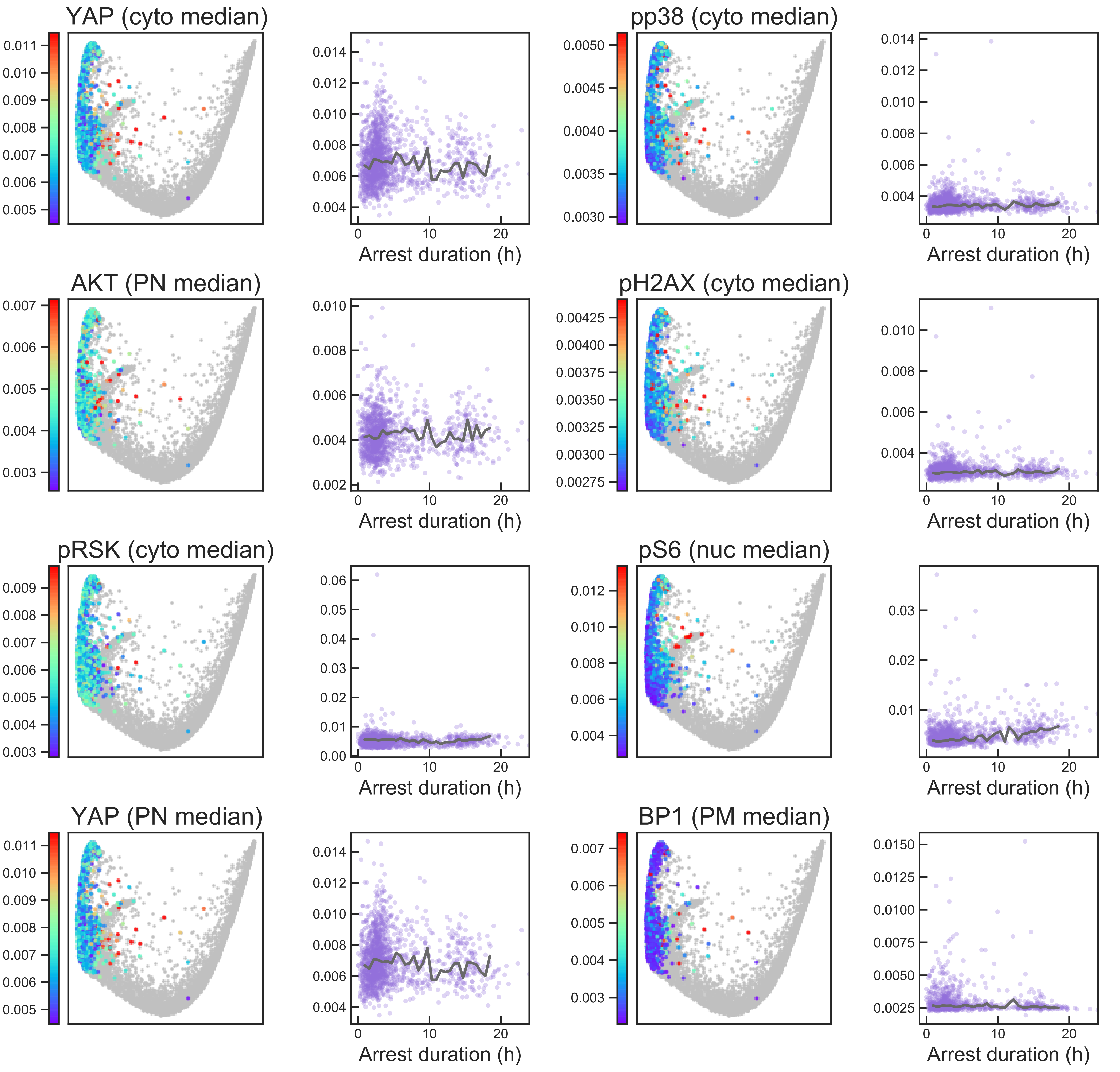

### Aux Fig. S2 - Arrest trajectory - Page 22

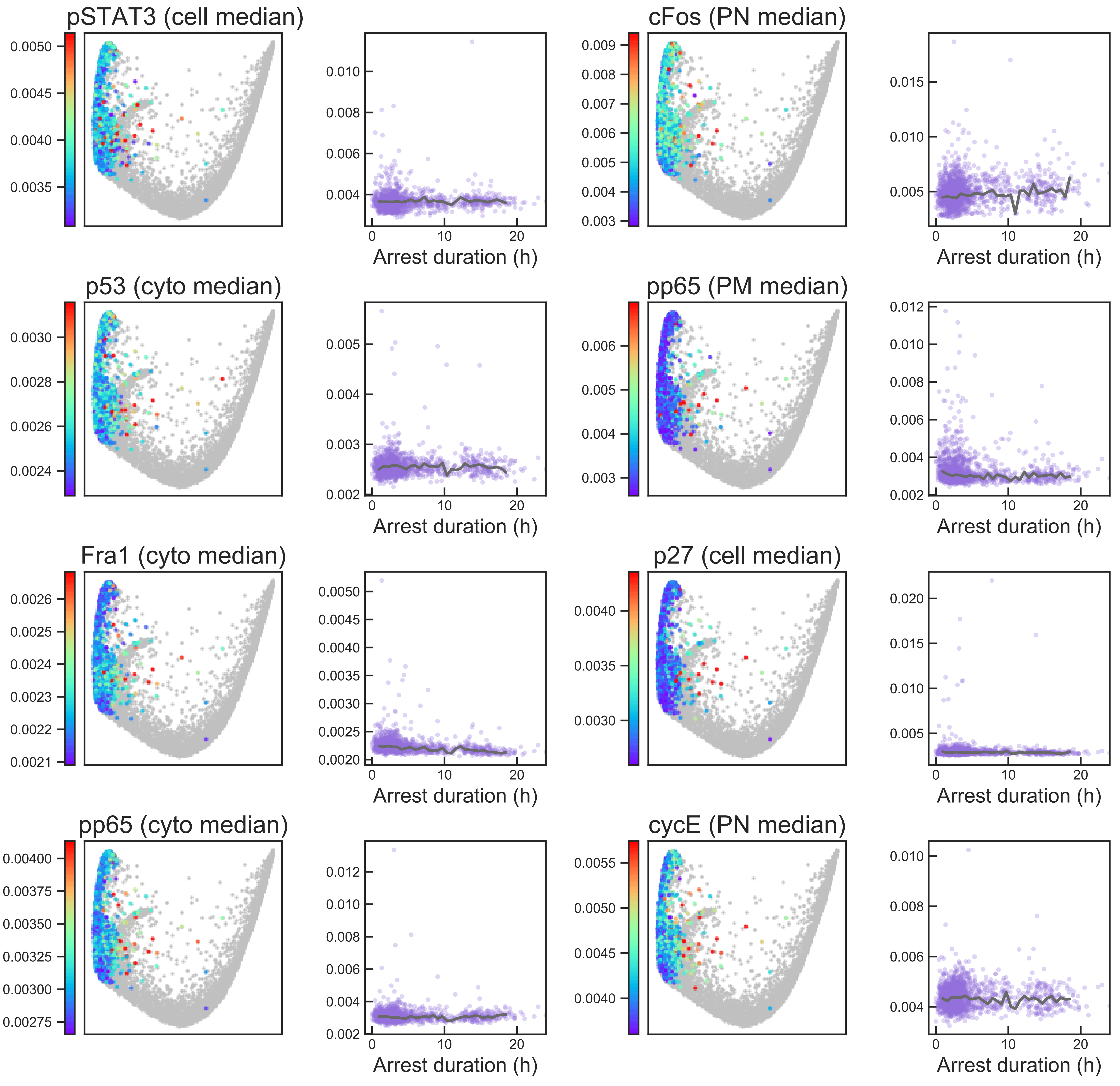

#### Aux Fig. S2 - Arrest trajectory - Page 23

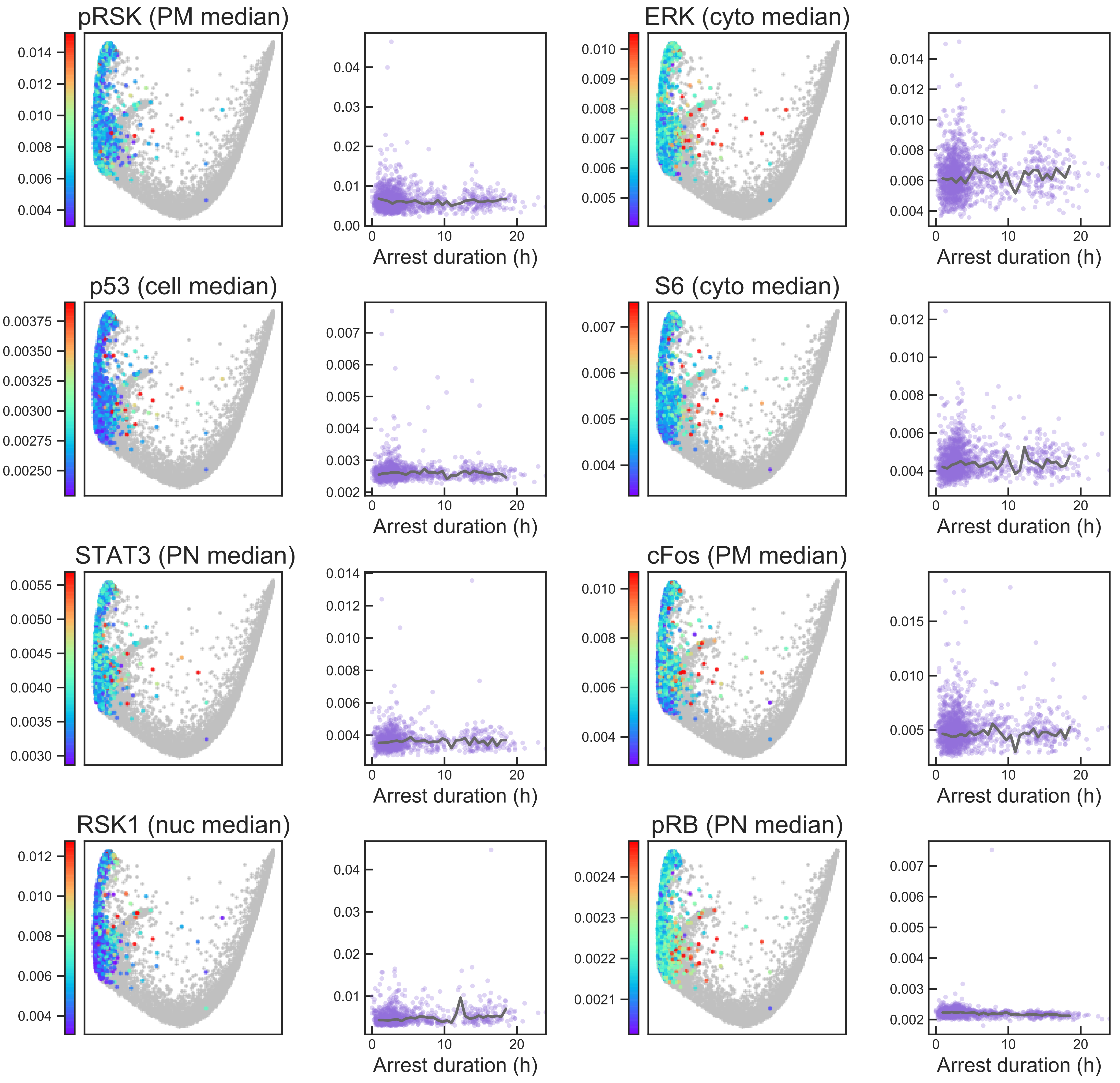

### Aux Fig. S2 - Arrest trajectory - Page 24

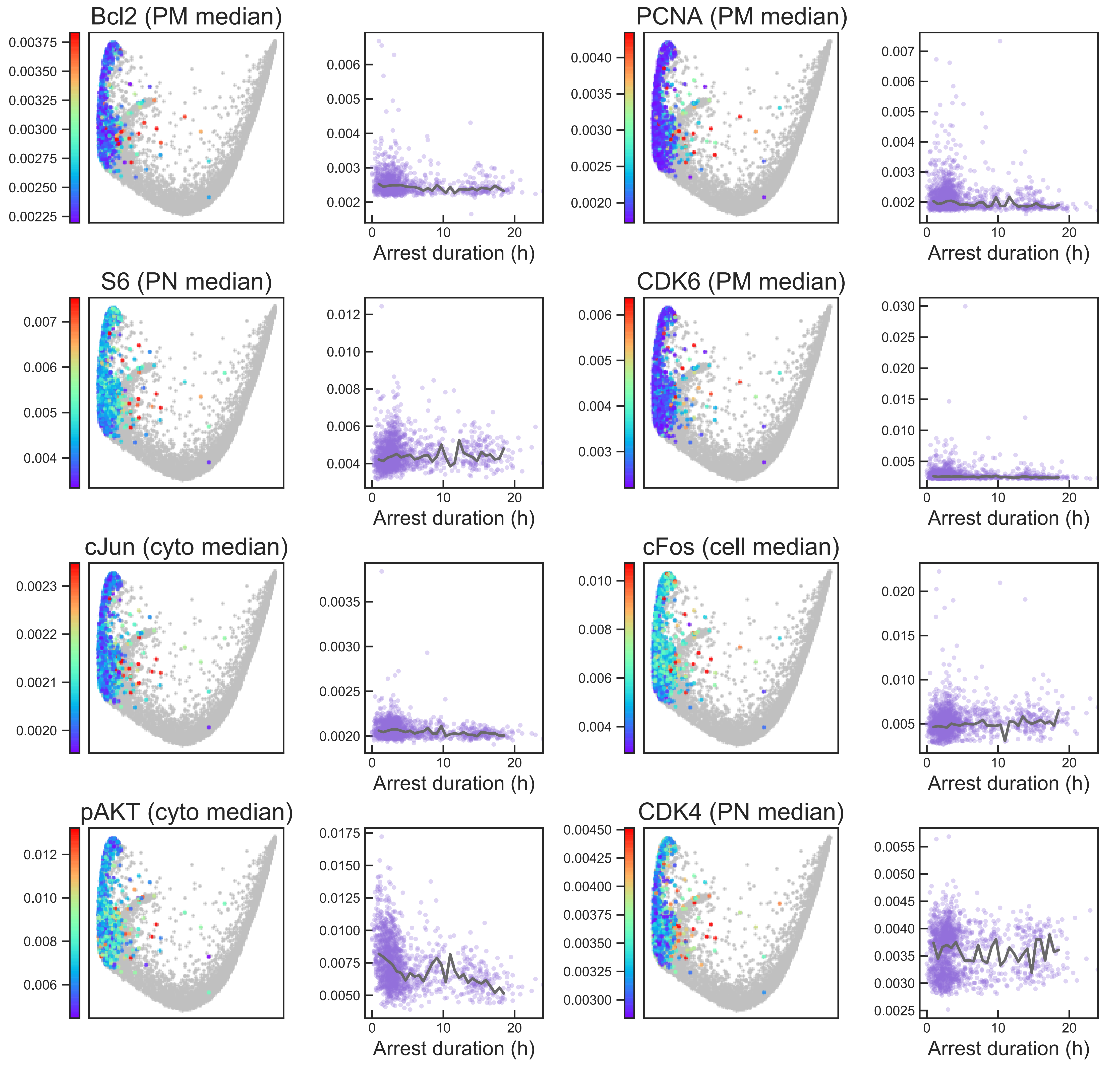

#### Aux Fig. S2 - Arrest trajectory - Page 25

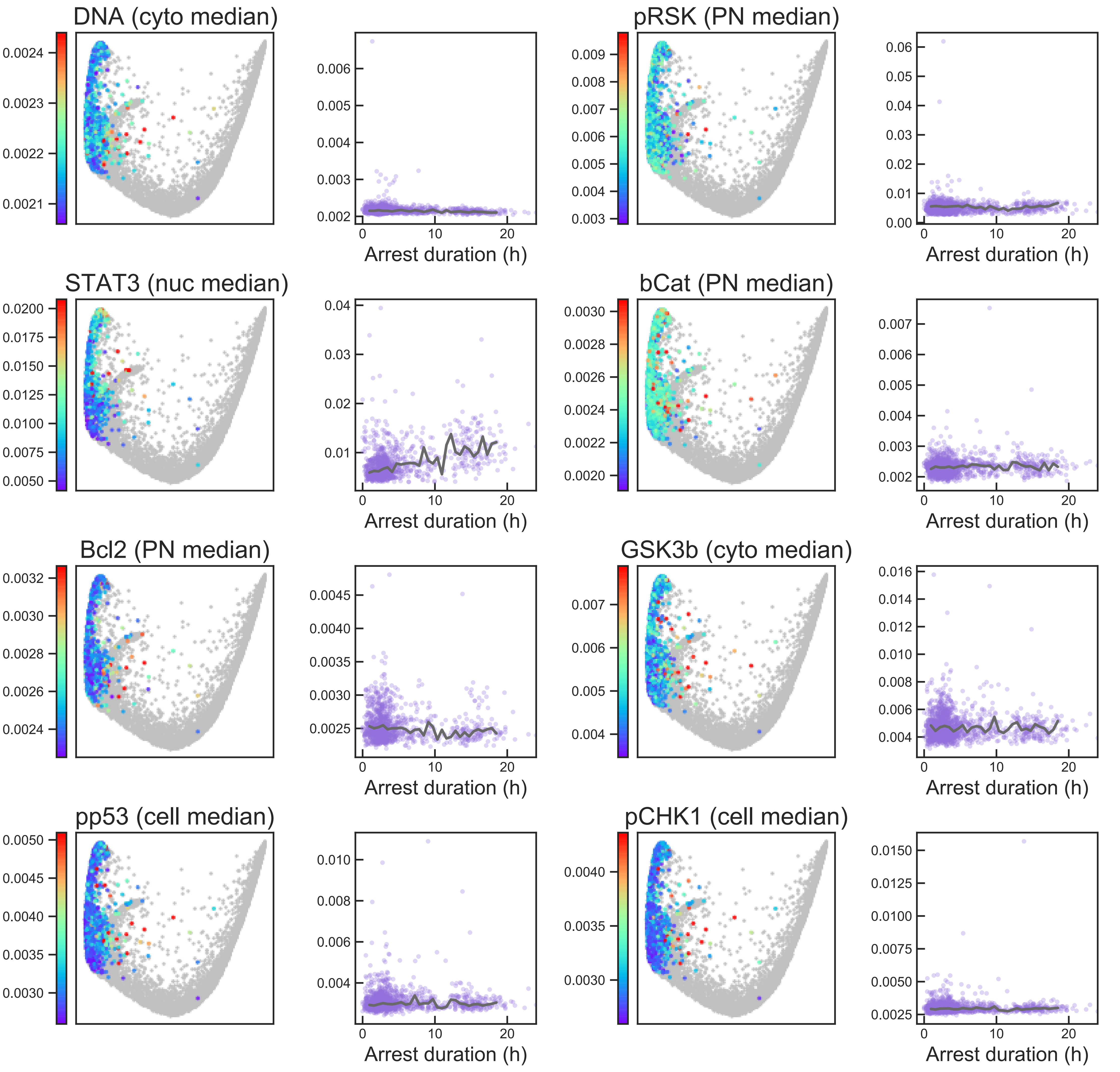

#### Aux Fig. S2 - Arrest trajectory - Page 26

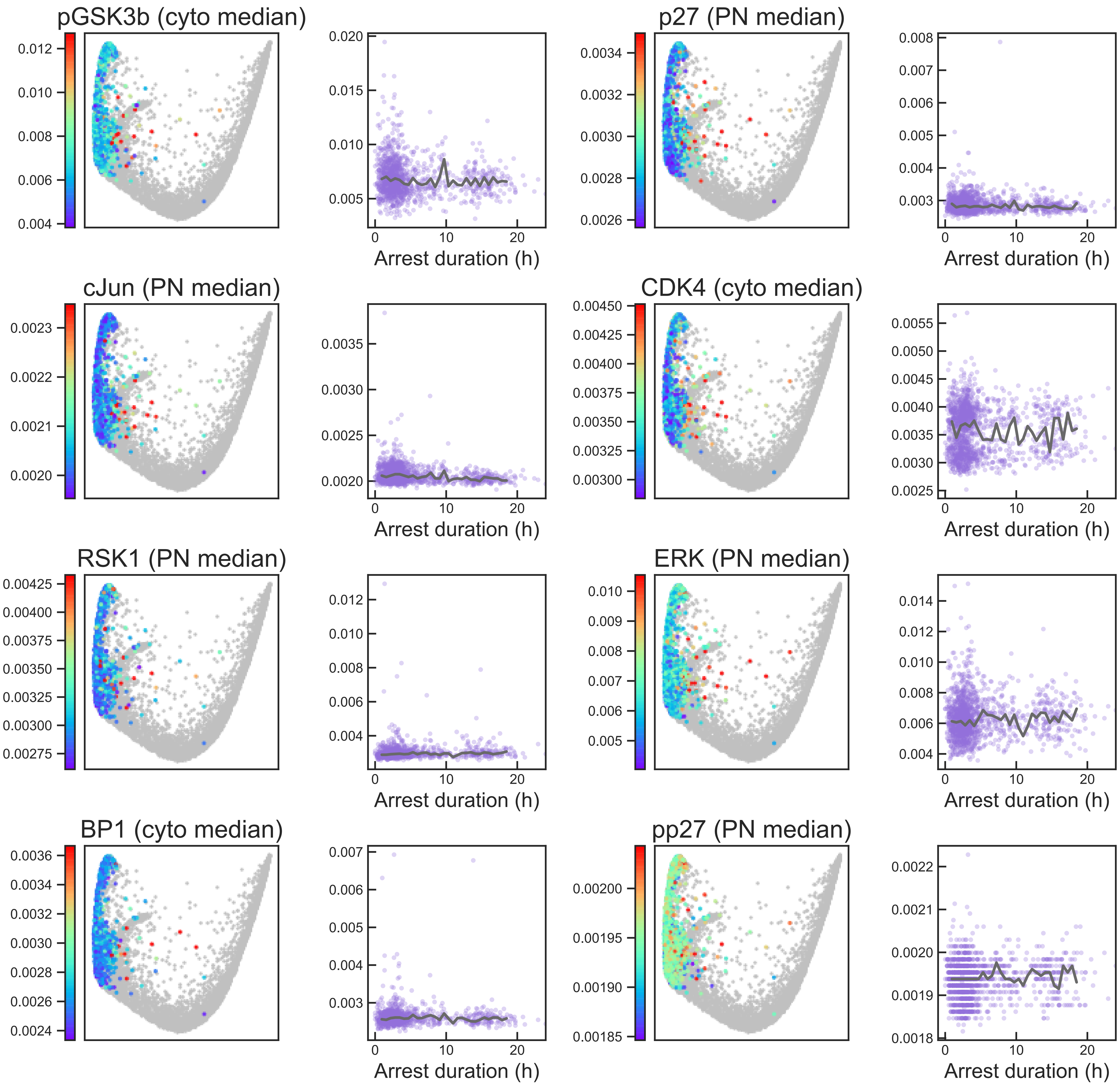

### Aux Fig. S2 - Arrest trajectory - Page 27

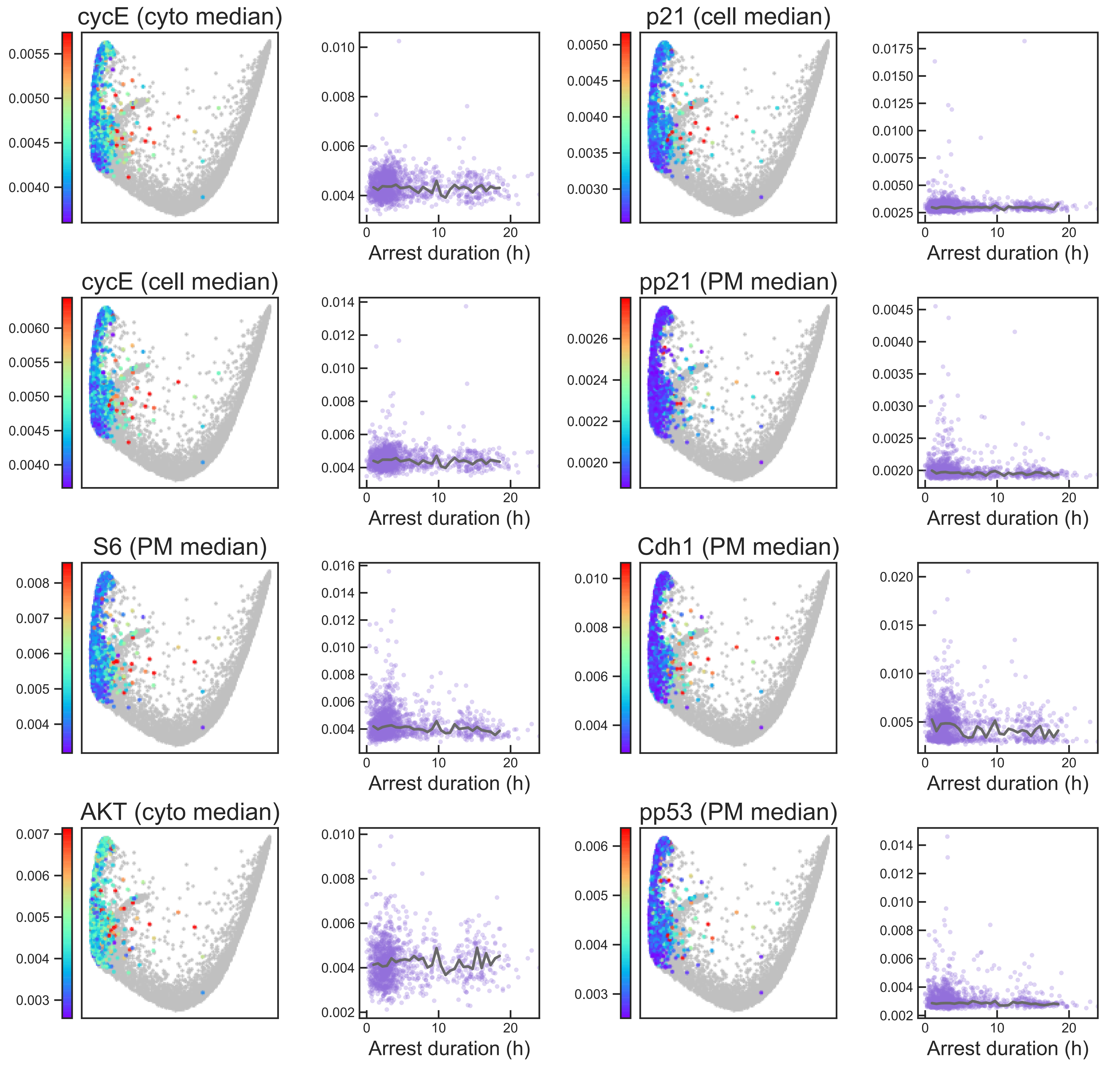

### Aux Fig. S2 - Arrest trajectory - Page 28

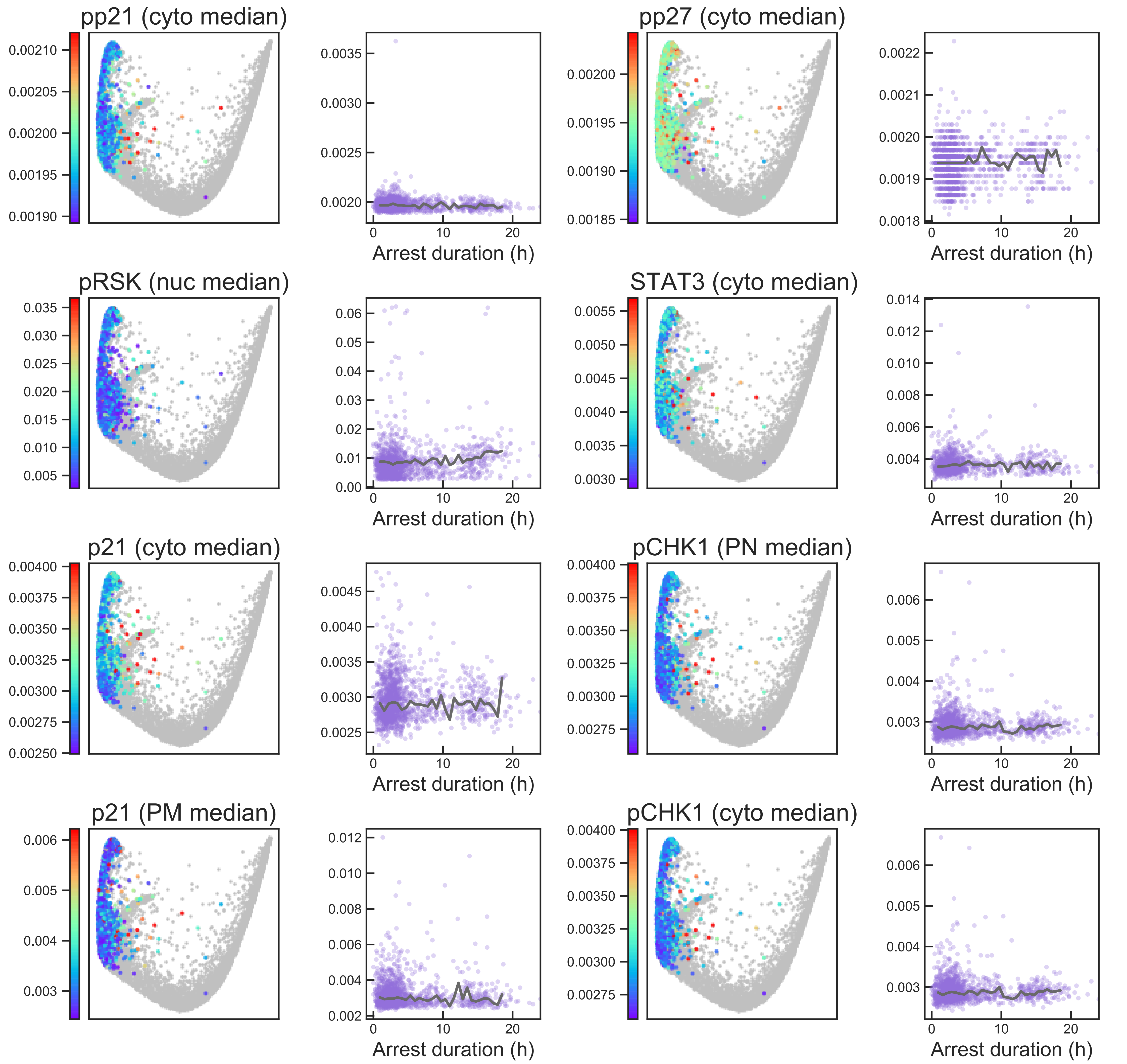

### Aux Fig. S2 - Arrest trajectory - Page 29

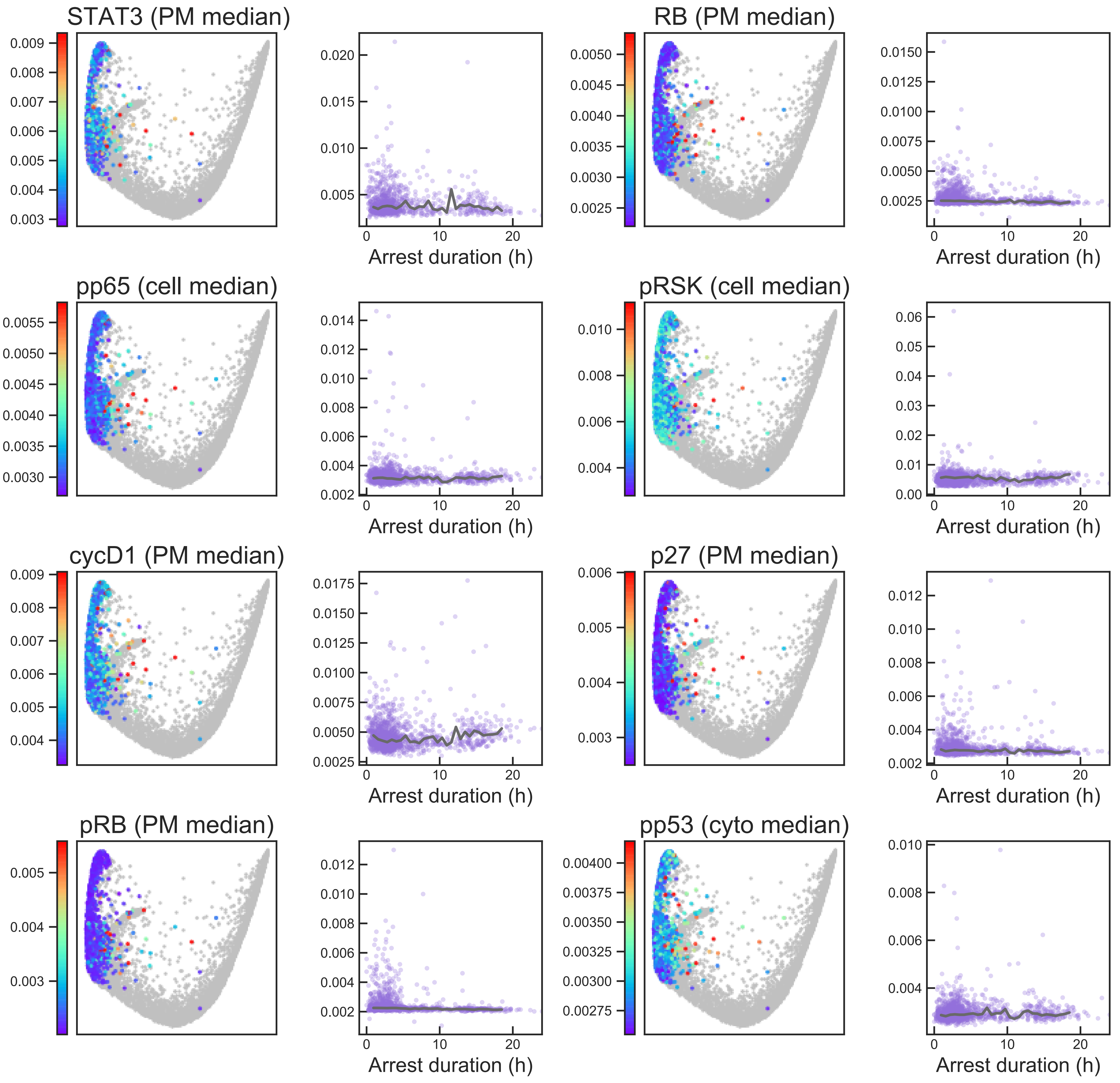

### Aux Fig. S2 - Arrest trajectory - Page 30

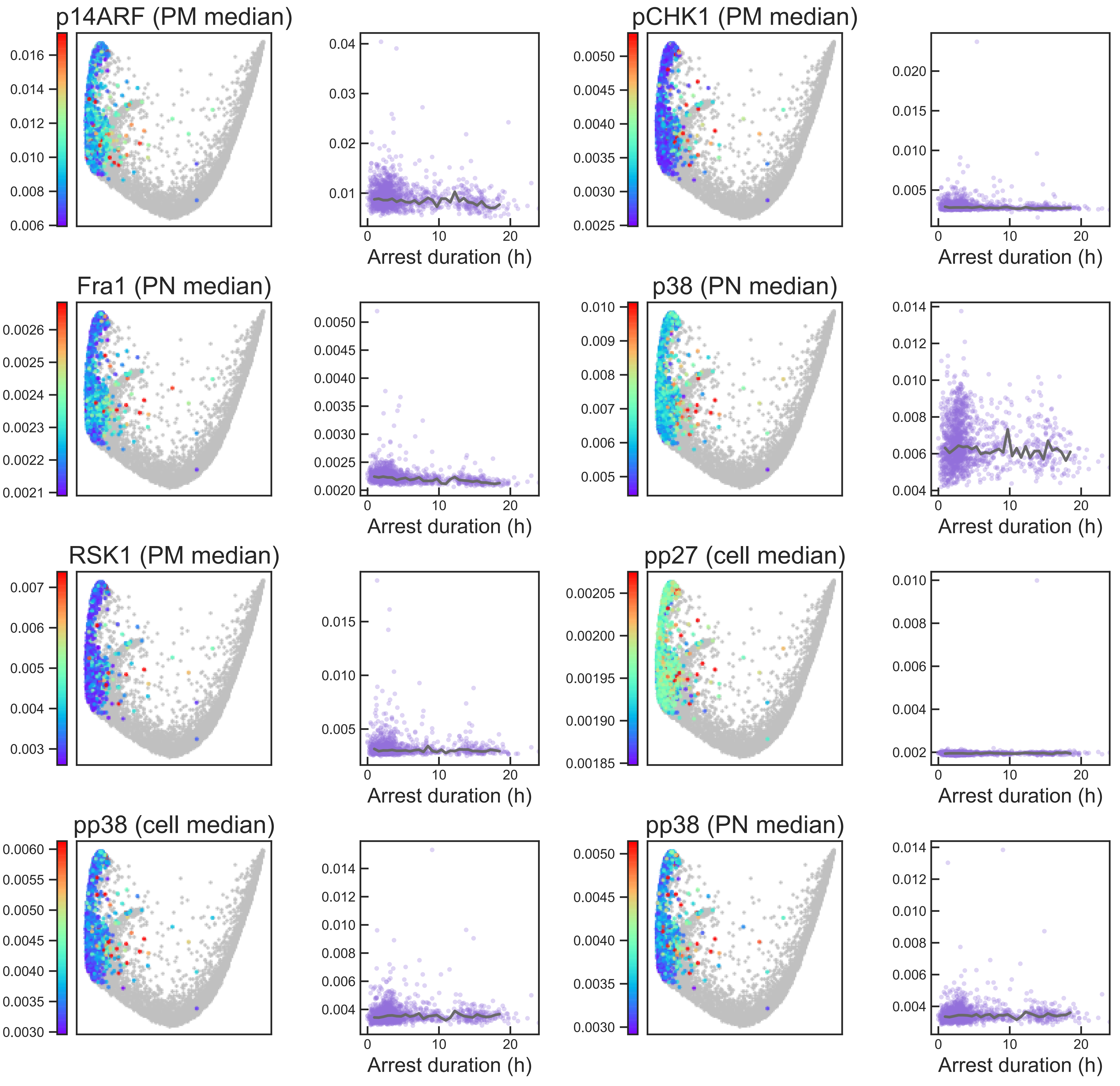

### Aux Fig. S2 - Arrest trajectory - Page 31

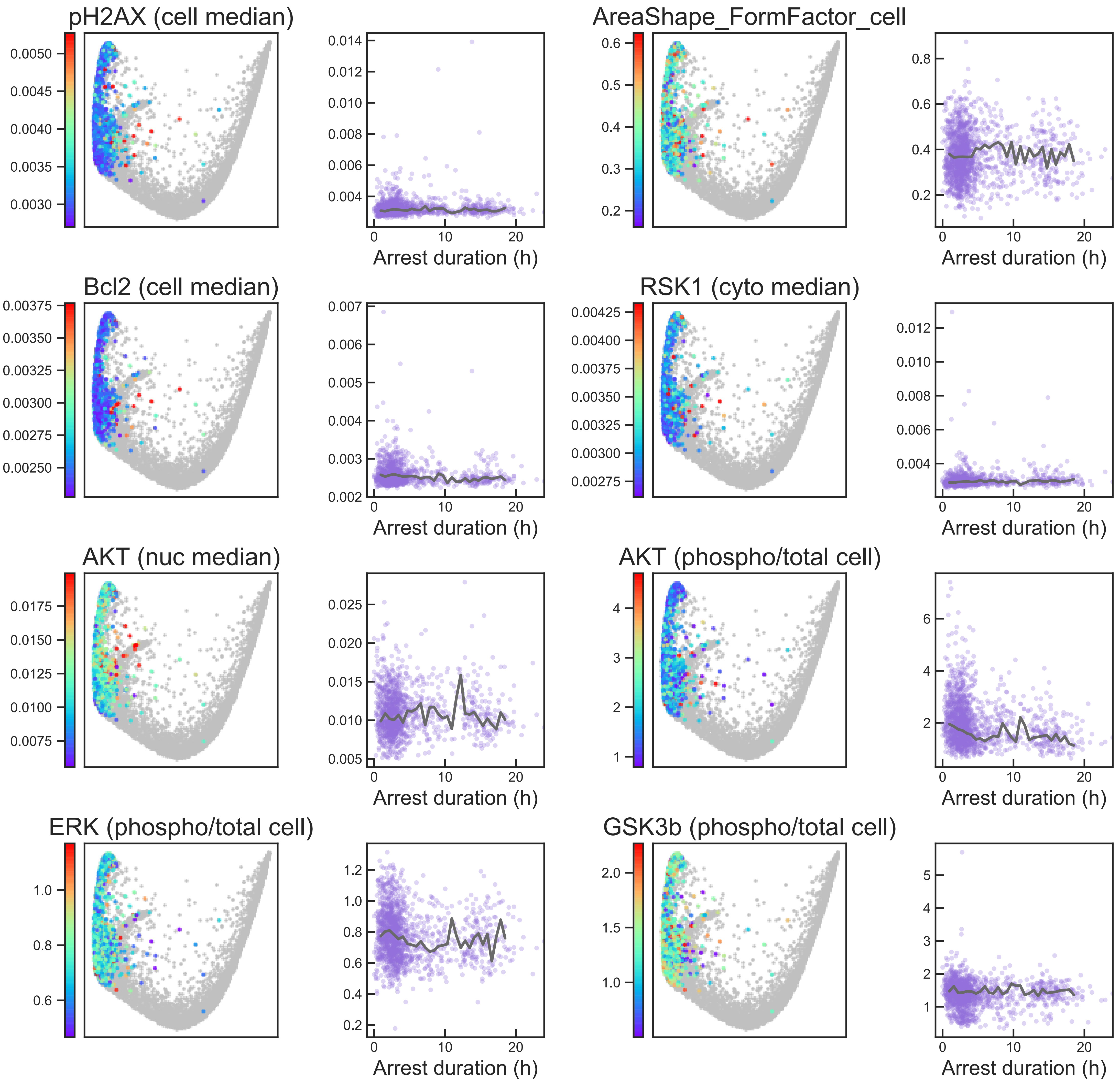

### Aux Fig. S2 - Arrest trajectory - Page 32

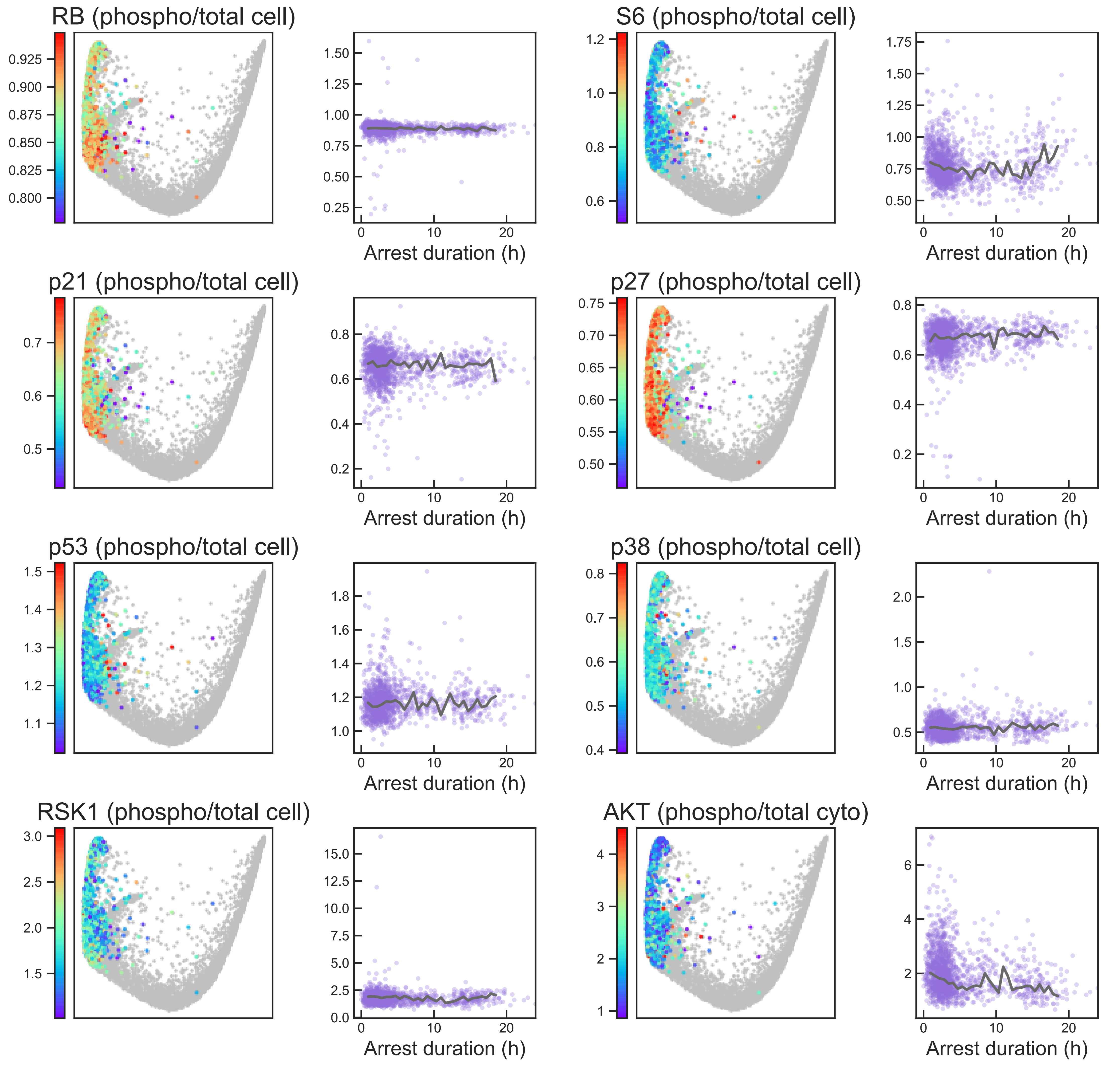

#### Aux Fig. S2 - Arrest trajectory - Page 33

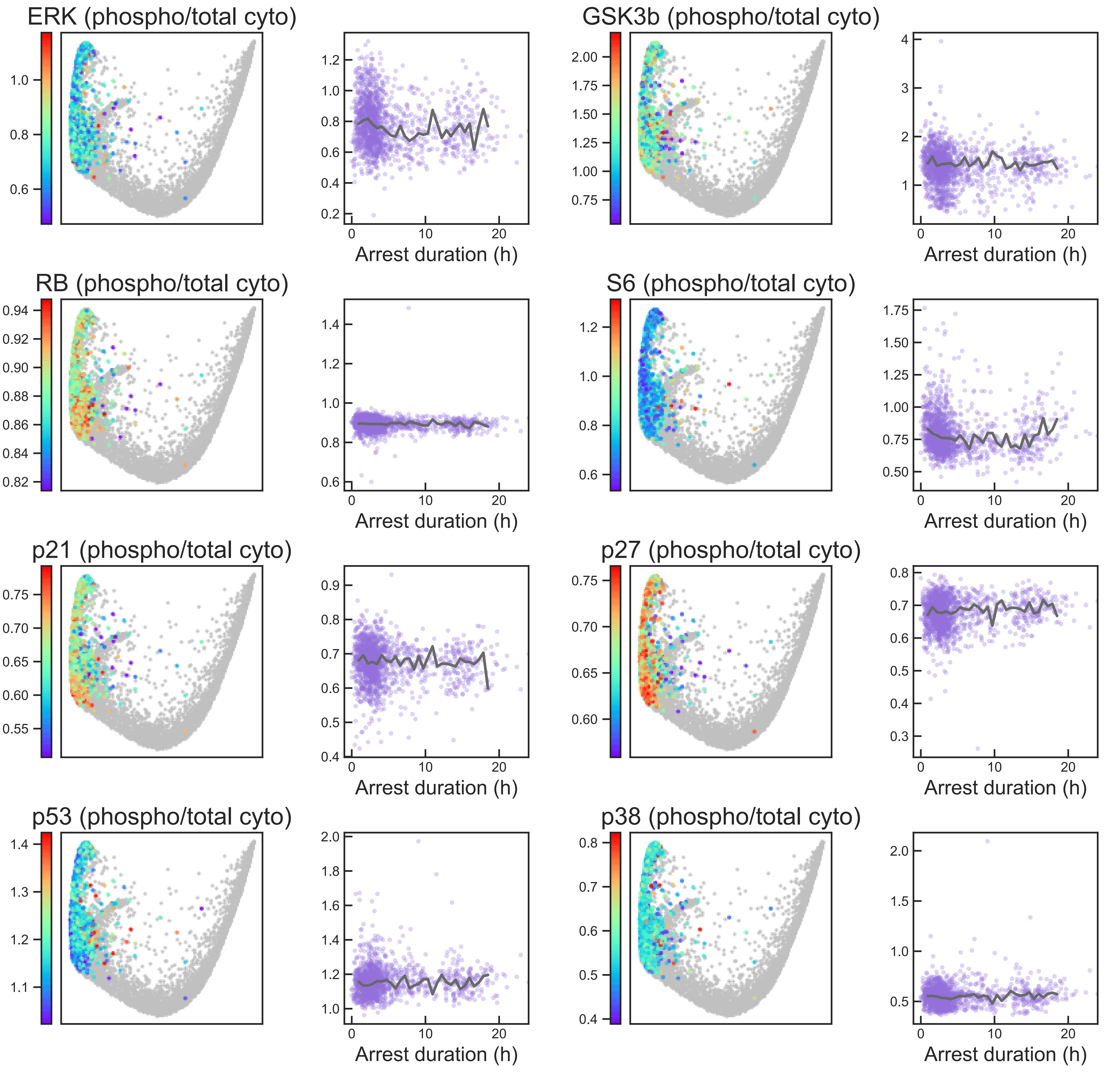

### Aux Fig. S2 - Arrest trajectory - Page 34

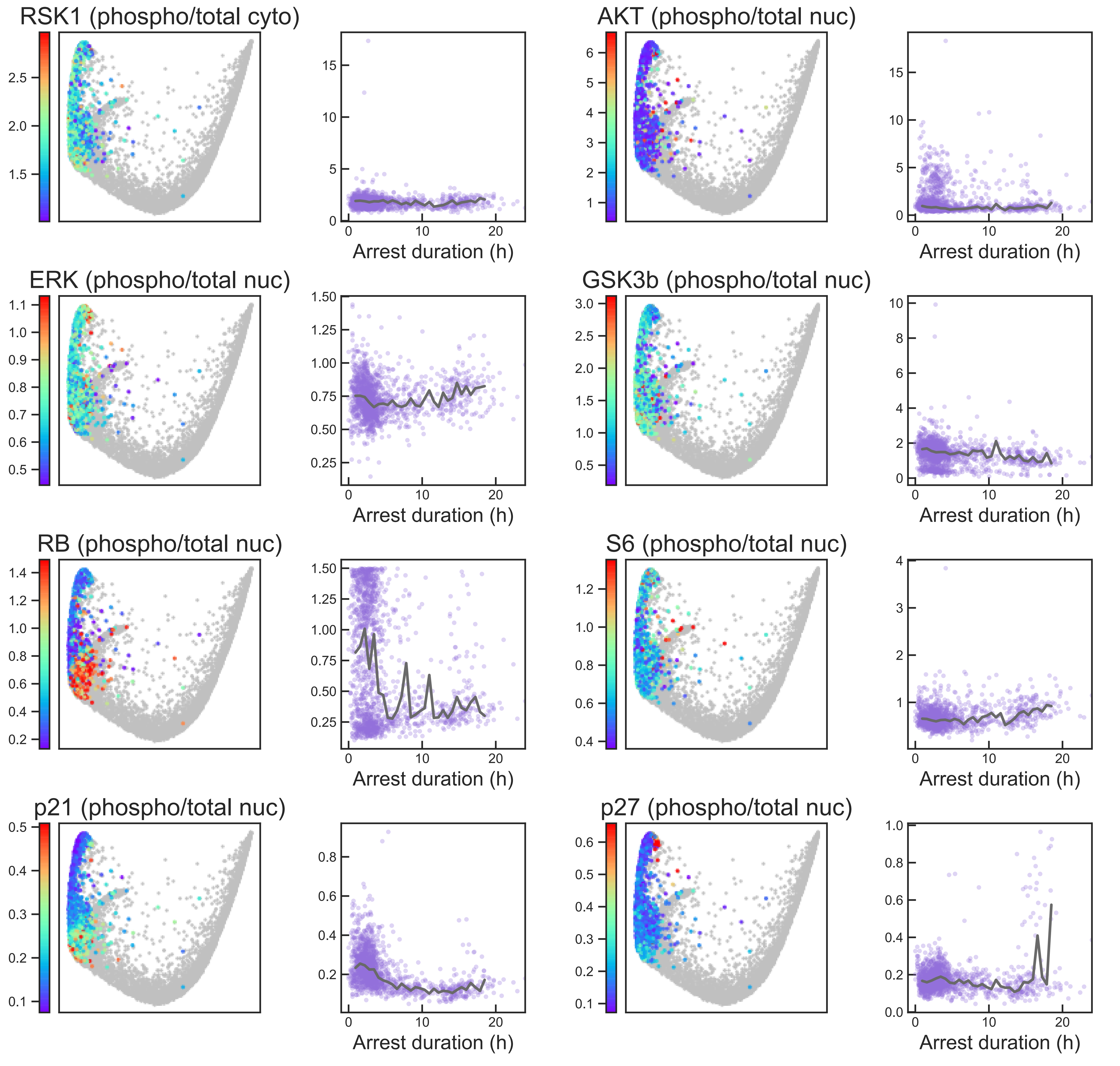

### Aux Fig. S2 - Arrest trajectory - Page 35

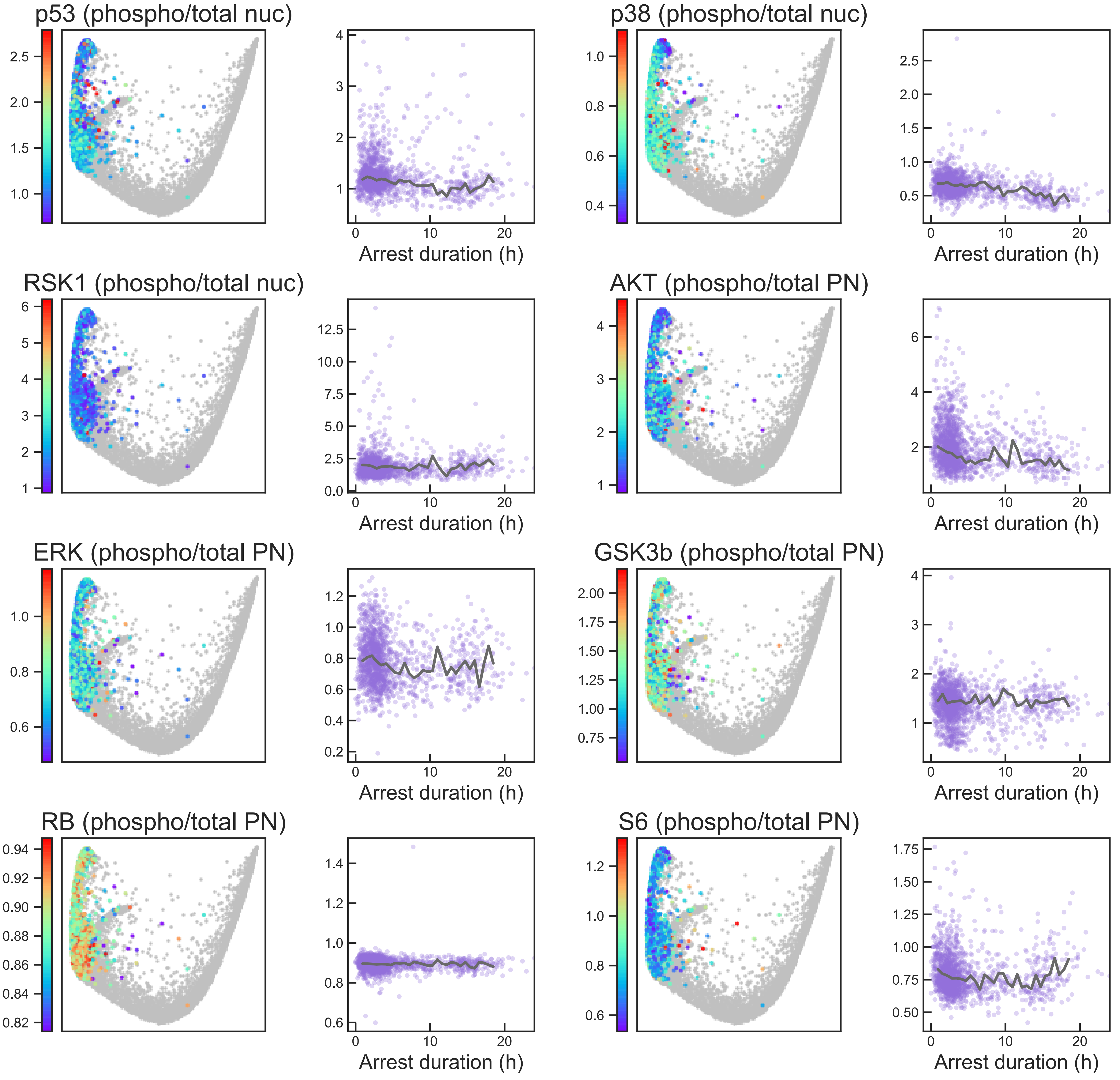

### Aux Fig. S2 - Arrest trajectory - Page 36

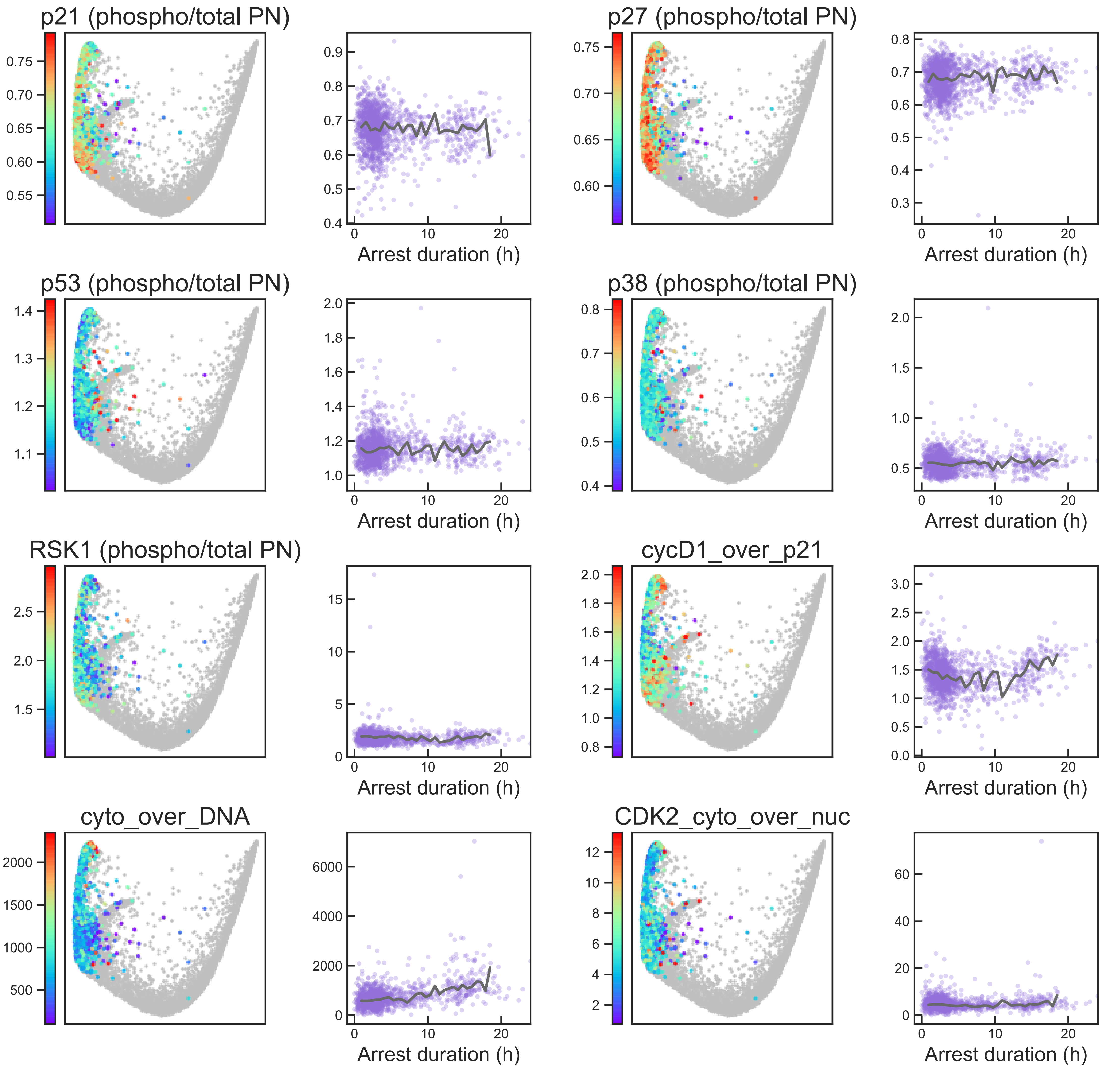
